# Automated community ecology using deep learning: a case study of planktonic foraminifera

**DOI:** 10.1101/2022.10.31.514514

**Authors:** Allison Y. Hsiang, Pincelli M. Hull

## Abstract

The development of deep learning methods using convolutional neural networks (CNNs) has revolutionised the field of computer vision in recent years. The automation of taxonomic identification using CNNs leads naturally to the use of such technology for rapidly generating large organismal datasets in order to study the evolutionary and ecological dynamics of biological communities across time and space. While CNNs have been used to train machine learning classifiers that can identify organisms to the species level for several groups, this vision of automated community ecology has yet to be thoroughly tested or fulfilled. Here, we present a case study of automated community ecology using a large dataset of Atlantic planktonic foraminifera for which the generation of species labels and morphometric measurements was completely automated. We compare standard community diversity metrics between the fully automated dataset and a “traditional” dataset with human-identified specimens. We show that there is high congruence between the results, and that machine classifications help avoid biases that can result in the inference of misleading biodiversity patterns. Our study demonstrates the viability and potential of fully automated community ecology and sets the stage for a new era of ecological and evolutionary inquiry driven by artificial intelligence.

## Introduction

The field of community ecology focuses on the organisation of and interactions between species that comprise an interacting biological network occupying a common geographic area. The study of ecological questions on a community level is essential for understanding the function, health, and dynamics of ecosystems and the relationship between the biosphere and the physical environment. Understanding these interactions has become crucial in light of rapid anthropogenic climate change and increasing evidence that we are currently experiencing the sixth mass extinction (Barnosky et al. 2011), with global biodiversity decreasing alarmingly across many groups. Conducting such studies requires large amounts of data in order to accurately represent the diversity and composition of a given biological community, and thus lend statistical rigour to the underlying analyses. Traditionally, collecting such data is a laborious, time-intensive task that requires preparation and curation of samples, identification of specimens by taxonomic experts, and manual measurement of morphological metrics of interest such as the length, width, and volume of individuals. The generation of these datasets is thus a rate-limiting factor in the study of community ecology, with the time and labor required becoming increasingly problematic in light of increasing rates of species turnover and ecological disruption in response to climate change (Barnosky et al. 2011; Ceballos et al. 2015; Ceballos et al. 2017), which in turn necessitate increased large-scale biodiversity monitoring.

The development of machine learning methods that are able to parse visual information with little to no human interaction in recent years provides a powerful new set of tools which may be fruitfully applied to the field of community ecology. Specifically, computer vision performance has experienced a renaissance thanks to the development of convolutional neural networks (CNNs), which are comprised of interconnected layers of “neurons” that perform convolutions (*i.e.,* mathematical transformations). Supervised deep learning algorithms using CNNs are able to achieve superhuman accuracies in classification tasks given appropriate hyperparameters and sufficient (in both quantity and quality) training data without the need for hand-engineered features. These networks require large training sets of image/label pairs as input in order to learn to correlate the visual information in an image with its associated identity. Careful preparation of the training data is necessary in order to avoid common issues such as overfitting and systematic bias. Automated classifiers using CNNs have been successfully trained for taxonomic identification of pollen grains (Sevillano et al. 2020; Olsson et al. 2021), insects (Valan et al. 2019; Hansen et al. 2020), planktonic foraminifera (Hsiang et al. 2019; Mitra et al. 2019; Marchant et al. 2020; Karaderi et al. 2022), among others. These models allow for rapid identification of large numbers of organisms with accuracies superior to those of human experts on average. Such datasets are essential for the study of community ecology and CNNs thus represent a revolutionary methodology for the future of this field.

Along these lines, several initiatives for generating such ecologically relevant data have begun operation in the last decade (*e.g.*, camera-trap datasets such as Snapshot Serengeti (Swanson et al. 2015)). However, while the generation of such datasets continues apace, to date the downstream analyses relevant to community ecology using such datasets have yet to be explicitly tested and validated (though see Marchant et al. (2020), which compares relative abundance, counts per sample, and fragmentation rates through time for eight benthic and six planktonic foraminifer species between their automated system and the Endless Forams (Hsiang et al. 2019) dataset). Here we present, to our knowledge, one of the first in-depth studies comparing relevant ecological measures and diagnostics between a traditional human-generated dataset and a CNN-generated dataset. We focus on planktonic foraminifera, a group of single-celled marine protists that play an essential role in oceanic carbon cycling and biological productivity and are used extensively to reconstruct paleoclimatic and oceanographic records. Using the Endless Forams database, we demonstrate that the ecological composition and diversity of both datasets is similar, while revealing certain biases and limitations of human-labeled datasets. We further evaluate the ecological characteristics of the combined human+machine dataset, thus paving the way for the use of machine-generated datasets for studies of community ecology.

## Materials & Methods

### Taxonomic and Shape Data

This study uses two datasets which are both derived from the Yale Peabody Museum Coretop Collection (YPMCC), which contains 61,849 planktonic foraminifer individuals above the 150-μm size fraction sourced from 34 Atlantic coretop samples (Elder et al. 2018). A subsample of 24,569 specimens were independently identified in quadruplicate by 24 taxonomic experts as part of the Endless Forams (http://endlessforams.org/) database (only identifications with ≥75% agreement between experts retained).

This subsample is not entirely random. All images were randomly selected from the available pool of images in batches of 5,000 or 10,000 images, the initial batch of 10,000 images were selected from all 34 sites. However, following feedback from the taxonomic experts, the remaining images were randomly selected from only those sites with good preservation (23 out of 34). In total, 3,020 images from the 11 sites with poor preservation were identified (12.3% of all identified images). Because the images from these 11 sites do not constitute a true random sample (as they were excluded from sampling rounds subsequent to the first), we removed these images from the dataset, thus restricting the original community to only the 23 sites with good preservation (Fig. S1). We also removed any individuals that were identified as not a planktonic foraminifer (9 individuals). This final set of 21,540 images and associated labels serves as our *human-identified dataset (HD)* and comprises 33 species.

The remaining 26,663 individuals from the YPMCC from the 23 sites with good preservation were not identified by human experts. We generated species labels for these individuals using a CNN (see details below). This set of images and CNN-predicted labels serves as our *machine-identified dataset (MD)*. These images are accessioned in the open-access data repository Zenodo (DOI: 10.5281/zenodo.7268451). As in the human dataset, we removed all objects identified as not a planktonic foraminifer (9 individuals), resulting in a final dataset of 26,654 individuals comprising 32 species (with *Globorotaloides hexagonus* being the single unrepresented species). As both of these datasets are derived from the same original populations, and their composition was randomly determined (after removing the individuals sampled from poorly preserved sites), they are representative of the same underlying biological community and should share ecological characteristics and properties. The relative composition of these datasets by site is shown in Fig. S2 in the Supplementary Materials.

The full dataset of 48,194 individuals is termed the *combined dataset (CD)*. Morphometric data (e.g., 2D area and perimeter, major and minor axis length, estimated volume using a dome base [see Hsiang et al. (2016)]) for these foraminifera were extracted from the measurements from Elder et al. (2018). A total of 2,410 individuals had no associated shape information, and these individuals were thus excluded from the analyses, resulting in 45,784 individuals. All labels and morphometric data are available in the Supplementary Materials.

### CNN-facilitated label prediction

We generated labels for the machine-identified dataset using the best-performing CNN described in (Hsiang et al. 2019). This CNN uses the VGG-16 (Simonyan & Zisserman 2014) architecture and achieves a top-1 accuracy of 87.4% (compared to an average human expert accuracy of 71.4%). We used this model to predict species labels for the foraminifera without human-assigned labels. The prediction script is available at https://www.github.com/palaeomachinist/foram-comm-ecol and is implemented using the Keras Python API (Chollet 2015) using the GPU-enabled TensorFlow backend (Abadi et al. 2016). The prediction was run on the Beella machine in the Hull Lab in the Department of Earth and Planetary Sciences at Yale University (6-core 2.4Ghz CPU, 32 GB RAM, 2x NVIDIA GeForce GTX 1080 GPUS [8 GB], Ubuntu 15.04.5 LTS).

### Measuring characteristics of community ecology

In order to assess whether machine-generated datasets are suitable for studies of community ecology, we measured several standard measures of community composition and biodiversity for the HD and the MD independently in order to determine whether they are equivalent. We measured the following characteristics: (1) α-diversity of all sites (species richness [Hill number *q*=0], Shannon index [Hill number *q*=1], and Simpson index [Hill number *q*=2]); (2) Pielou’s evenness; (3) ß-diversity between all sites; (4) Rank abundance; (5) Latitudinal abundance; (6) Latitudinal body size trends vs. sea-surface temperature (SST); and (7) Sitewise relative abundance. We use the Chao version of all α-diversity measures, i.e., richness *sensu* Chao (1984; 1987), transformed Shannon diversity (Chao et al. 2013), and transformed Simpson diversity (Good 1953; Chao et al. 2014). We also compared the HD and MD as independent assemblages following Chao et al.’s (2020) diversity comparison procedure, whereby we calculate for the HD and MD the sample completeness profile, asymptotic and empirical diversity estimates, sample-size- and coverage-based rarefaction/extrapolation curves, and the evenness profile (*sensu* Chao & Ricotta (2019)). These profiles allow us to compare diversity patterns between the HD and the MD while incorporating information about sample coverage integrated with the calculation of Hill numbers (Hill 1973) parameterised by order *q.*

All Chao diversity metrics were calculated using the “iNEXT” package (ver. 2.0.20) (Hsieh et al. 2016) with data classified as abundance data. Although these metrics assume sampling with replacement, it has been previously noted that results from samples taken without replacement (as is the case with our dataset) should be similar if the sample is small relative to the size of the underlying assemblage (Gotelli & Colwell 2011). Rank abundance curves were calculated using the *rankabundance* function from the “BiodiversityR” package (ver. 2.12-3) (Kindt & Coe 2005). Relative abundances were calculated using the *make_relative* function in the “funrar” package (Grenié et al. 2017). Jaccard indices were calculated using the *vegdist* function in the “vegan” package (ver. 2.5-7) (Oksanen et al. 2020). Body sizes were extracted from the Elder et al. (2018) dataset; for the volume estimates, the domed base measurement was used (see Hsiang et al. (2016)). The diversity comparison procedure was implemented using the “iNEXT.4steps” package described in Chao et al. (2020) with data classified as taxonomic abundance data. Table S1 shows calculated values for the special cases where *q* = 0, 1, 2 for all presented diversity metrics. All code and scripts used to process the data can be found in the GitHub repository available at https://github.com/palaeomachinist/foram-comm-ecol.

### Sea surface temperature

All SST data was taken from the NOAA Extended Reconstructed Sea Surface Temperature (ERSST) v5 dataset (Huang et al. 2017), which uses Empirical Orthogonal Teleconnections (EOTs) to reconstruct SSTs on a 2° × 2° grid using data from the International Comprehensive Ocean-Atmosphere Dataset (ICOADS). The monthly SST records for the time period between 2000 and 2018 were used to calculate the zonal mean SST (dashed green curve in Fig. 3) and plot the average SST spatial map for the Atlantic Ocean (Fig. S1) using CDO (Schulzweida 2019).

## Results

The rank abundance curves for the human (HD), machine (MD), and combined (CD) datasets are shown in Fig. 1. While the exact ordering of species in the rank abundance curves for the HD (Fig. 1a) and MD (Fig. 1b) differ, the same 10 species comprise the top 10 ranks. These top 10 ranks account for 84.6%, 81.9%, and 83.1% of the total abundance for the HD, MD, and CD, respectively. In both cases, *Globigerinoides ruber* occupies the top rank, comprising 29.7%, 26.4%, and 26.4% of the total abundance of each respective dataset. The rankings of *Globigerinoides sacculifer* and *Globigerina bulloides* (ranks 2 and 5) are switched between the two datasets, as are the rankings of *Globorotalia inflata* and *Neogloboquadrina pachyderma* (ranks 6 and 7) and *Globorotalia truncatulinoides and Globigerina falconensis* (ranks 8 and 10). The remaining rankings are the same between the HD and the MD. The rank abundance plot of the combined dataset (Fig. 1c) follows the same ordering as the ranking for the MD except for the positions of *G. inflata* and *N. pachyderma*, which are flipped (as in the ordering of the HD). All rank abundance data are available in the Supplementary Materials.

**Fig. 1.**
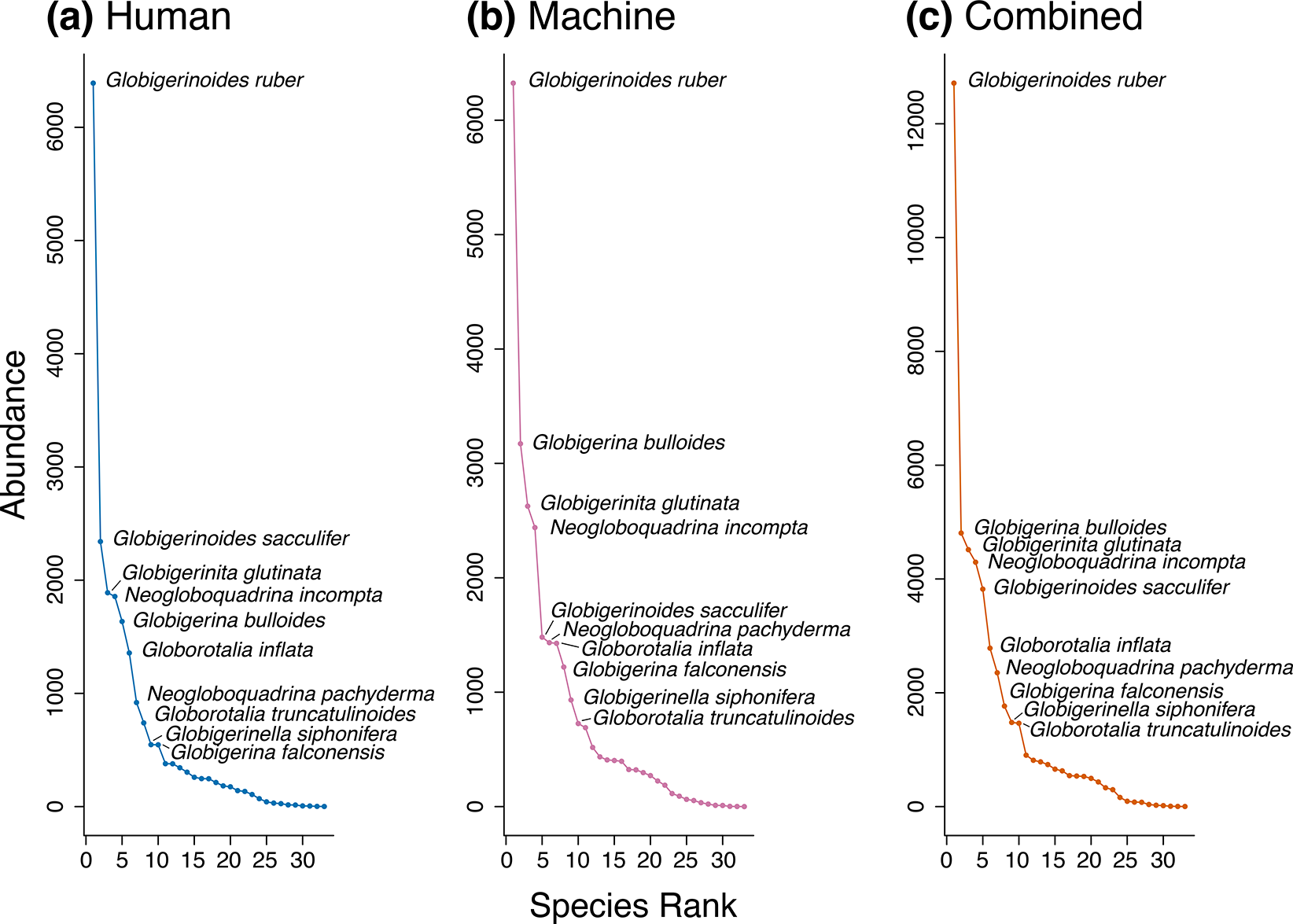
Rank abundance curves. (a) Human dataset (HD); (b) Machine dataset (MD); (c) Combined dataset (CD). The ten most abundant species for each dataset are labeled. Although the relative ordering shows some differences between the human (a) and machine (b) datasets (see text), the same ten species comprise the most abundant species in both datasets.

The normalized abundance maps (Fig. 2) show similar distribution patterns for each species amongst the datasets, with the maps being essentially identical when the number of samples (N) is large (*e.g.,* as with *Neogloboquadrina incompta*, which has N_HD_=1,856 and N_MD_=2,439 [Fig. 2a]). In general, marked deviations between the human and machine maps are only seen when N is small and clearly does not represent a statistically robust sample (*e.g.,* as with *Globigerinita uvula*, which has N_HD_=6 and N_MD_=11 [Fig. 2b]). Normalized abundance maps for all species can be found in the Supplementary Materials. In the species with high values of N, we observe two recurring inverse patterns of normalized abundance: either increasing normalized abundance from the equator to the poles (as exemplified by *Globorotalia inflata* [Fig. 2c]; this pattern is also seen in *N. incompta* [Fig. 2a]), or decreasing normalized abundance from the equator to the poles (as exemplified by *Globigerinoides ruber* [Fig. 2d]). Fig. S3 shows exemplars of species-specific latitudinal normalized abundance curves (*i.e.,* the best fit second degree polynomial) depicting the patterns of poleward-increase (*G. inflata;* Fig. S3a) and poleward-decrease (*G. ruber;* Fig. S3b). The exhibited patterns generally match between the HD and the MD within a species, with divergent patterns occurring primarily in species with N_CD_ < 300. Such a low N_CD_ results in only ~150 individuals per dataset, which is unlikely to be a statistically robust sample from which to reconstruct diversity patterns across the Atlantic. Some species, such as *Globigerina falconensis*, exhibit neither pattern (Fig. S3c). Instead, *G. falconensis* exhibits a gradual increase in abundance from the southern to the northern latitudes. While this may reflect a true biological pattern, such a pattern would also result from geographical sampling bias in the northern hemisphere, which may also explain the offset between the HD and MD curves observed in *Neogloboquadrina incompta* (Fig. S3d; see below for further discussion).

**Fig. 2.**
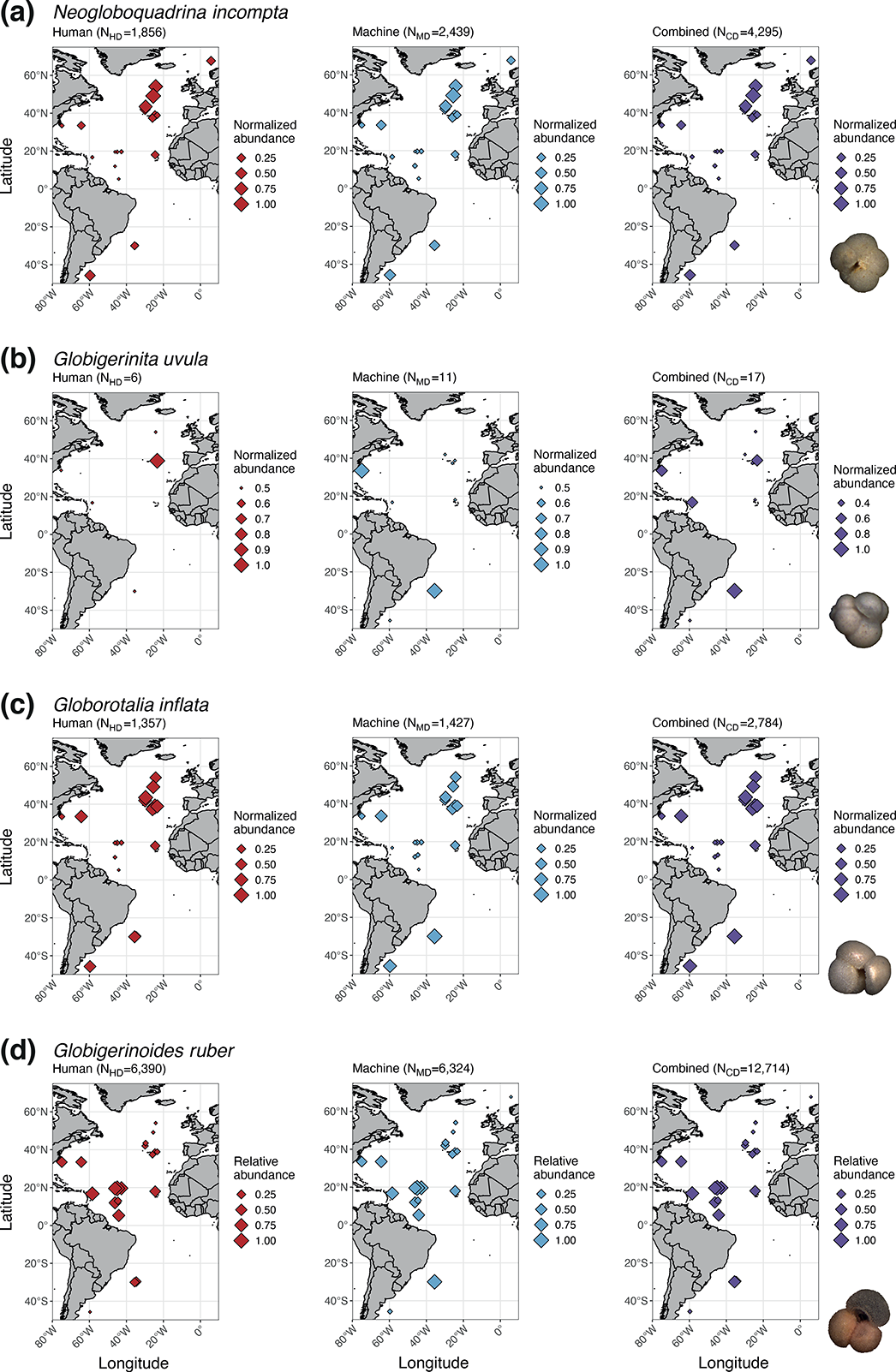
Species-specific latitudinal normalized abundance maps. The selected maps show various patterns observed in the datasets. In general, the normalised abundance patterns match for species with high N, as in (a) *Neogloboquadrina incompta* (N_CD_=4,295). Large differences in normalised abundance patterns are seen primarily in datasets with low N, as in (b) *Globigerinita uvula* (N_CD_=17). Many species exhibit either a pattern of increase in normalised abundance towards the poles, as in (c) *Globorotalia inflata* (N_CD_=2,784), or a pattern of decrease in normalised abundance towards the poles, as in (d) *Globigerinoides ruber* (N_CD_=12,714). Species exemplar images sourced from the Endless Forams database.

Fig. 3a shows the latitudinal (observed) species richness curves for the human, machine, and combined datasets. As seen from the best-fit second-order polynomial curves, all datasets show the same pattern of higher species richness near the equator and decreasing species richness towards the poles, *i.e.,* the latitudinal species diversity gradient that has long typified marine ecosystems and zooplankton diversity patterns (Yasuhara et al. 2012). We note that the richness curve for the HD is negatively offset from the curves for the MD and the CD, and that this offset appears to be greater towards the poles. This pattern of lower diversity in the HD is also observed in the α-diversity metrics (Fig. S4). While this offset may be caused in part by the smaller number of individuals in the HD compared to the MD (by ~5,000 specimens), it is unlikely to be the only cause given that the individuals were randomly sampled (see Materials & Methods). We suggest two possible related contributing factors: (1) Human bias against identification of rare species and (2) Sampling bias in the northern vs. southern hemispheres.

**Fig. 3.**
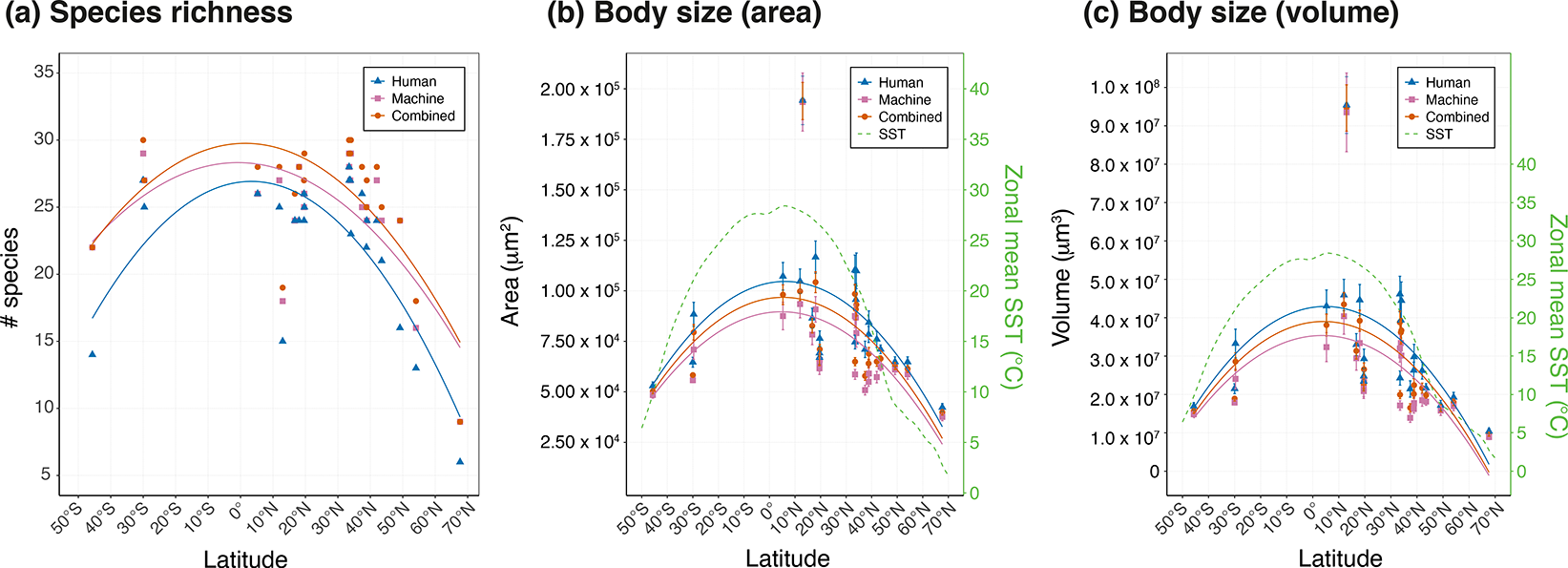
Latitudinal patterns in species richness and body size. (a) Observed species richness for the HD, MD, and CD plotted against site latitude. The curves represent best-fit second-order polynomials for each dataset. (b) Average 2D enclosed area (μm^2^) of all individuals at a site with 95% confidence intervals plotted against site latitude. (c) Average estimated volume (μm^3^) of all individuals at a site with 95% confidence intervals plotted against site latitude. Sea-surface temperatures (SSTs) are zonal means between 2000 and 2018 (see Materials & Methods).

As discussed in Hsiang et al. (2019), human identifiers tend to preferentially assign species identities that they perceive to be more numerous to ambiguous individuals. That is, if a community includes a common taxon A and a rare taxon B that closely resembles taxon A, then human classifiers will often incorrectly identify taxon B as taxon A. The same “pull towards the common” bias does not affect machine classifiers to the same extent. This effect may be pronounced at the poles, where there are fewer species overall, and planktonic foraminifer communities are relatively dominated by *Neogloboquadrina pachyderma* and *N. dutertrei.* Indeed, when comparing species relative abundances at the highest latitude site in our dataset (IPE.08295; Fig. 4), we see that the relative abundance of *N. pachyderma* and *N. dutertrei* are higher for the HD than for the MD and the CD. The same is true for the site with the lowest latitude (IPE.08248), although less pronounced (though we note that the community at site IPE.08248 is also relatively less dominated by these two species compared to IPE. 08295). This pattern can be seen in the exemplar sitewise species distribution plots (Fig. S5; all plots available in Supplementary Materials), where for sites where one or two species predominate, the number of individuals assigned to those species by human identifiers is much higher than the number assigned by the trained CNN (*e.g.,* site IPE.08171 and site IPE.08316; Fig. S5a-b). In contrast, sites with more even communities tend to have more individuals per species in the machine dataset (*e.g.,* site IPE.08378 and IPE.08262; Fig. S5c-d), as would be expected given that the machine dataset contains more specimens overall.

**Fig. 4.**
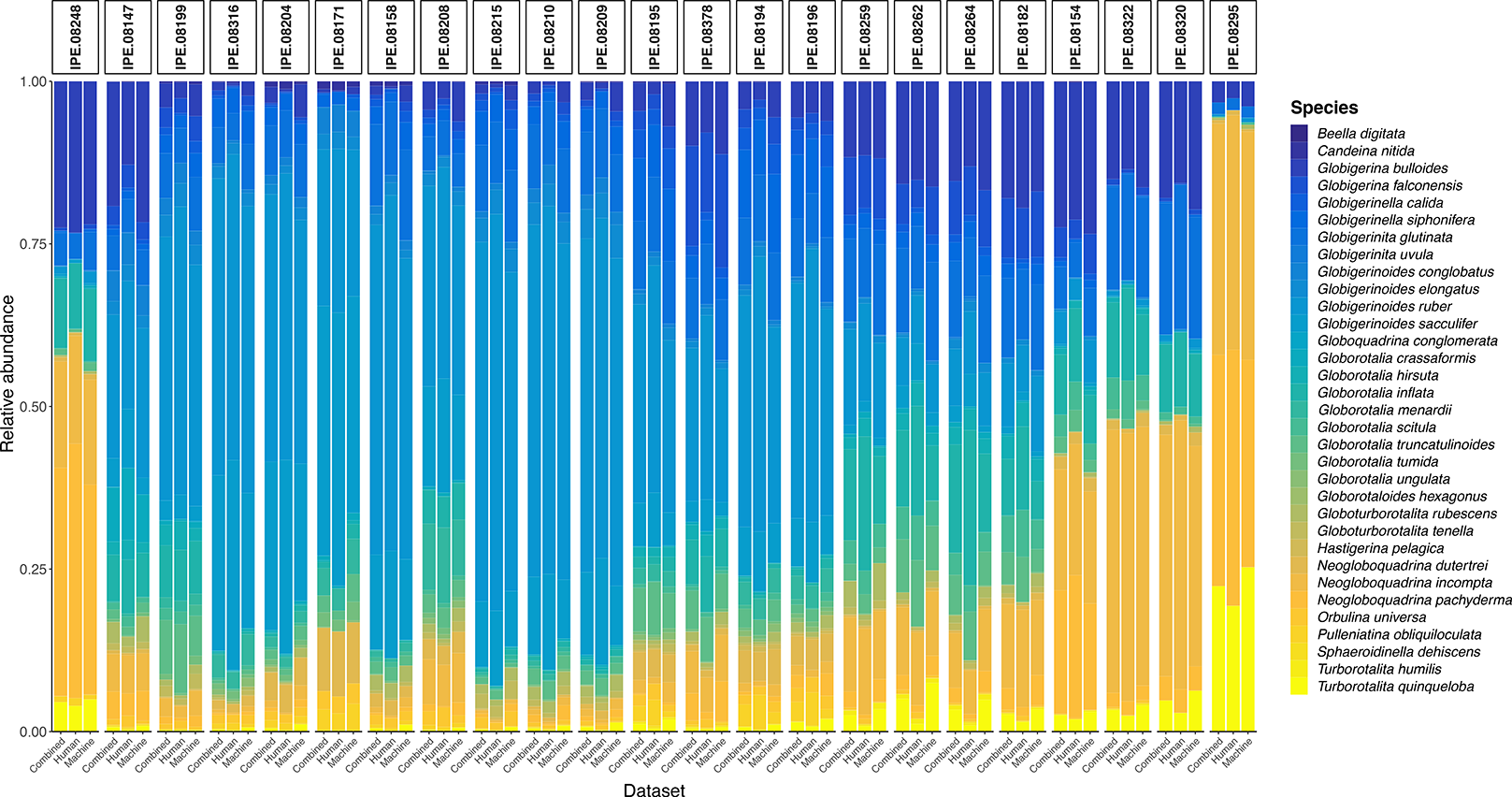
Sitewise relative abundance. Within each site, relative abundances are compared between the combined, human, and machine datasets. In general, relative abundance profiles match well between datasets. Sites are ordered from southernmost (left; 45.675°S) to northernmost (right; 67.65°N) latitude, which makes diversity patterns — such as the relative abundance of *Neogloboquadrina dutertrei* and *N. incompta* increasing towards the poles — apparent.

It is well known that biodiversity studies are affected by sampling bias, such as the far greater number of inventories taken from the northern compared to the southern hemisphere. For instance, Garraffoni et al. (2021) describe a pervasive inventory completeness bias in marine meiobenthic fauna, with the difference between observed and expected species richness being much higher in the southern vs. northern hemisphere. A corollary of this bias is that reconstructed diversity patterns in the northern latitudes are likely to be more complete and accurate than those in the southern latitudes. In our dataset, only three sites are found in the southern hemisphere, and the latitudinal patterns we describe must be evaluated in light of this bias, particularly for species with relatively low abundance. In addition, the meta-effects of geographical sampling bias on the performance of human taxonomists must also be considered. As inventories from the southern hemisphere are less numerous, taxonomists will also be less familiar with the extant diversity in sites from southern latitudes, particularly rare species. Given the observed human bias against identifying rare species, these biases are likely to have additive effects that result in human-classified datasets consistently underestimating species diversity in the southern hemisphere, a pattern we see in Fig. S3a-b, d. In the case of *N. incompta* (Fig. S3d), we actually see this discrepancy between the HD and the other two datasets disappear as we move towards the northern hemisphere. In our dataset, there are 653 specimens of *N. incompta* from the southern hemisphere (N_S_), compared to 3,642 in the north (N_N_). This pattern where N_N_ ≫ N_S_ is exhibited throughout our dataset.

We observe the familiar pattern of small body sizes towards the poles and increasing body size towards the equator in all datasets, for both 2D enclosed area (Fig. 3b) and estimated volume (Fig. 3c). This pattern follows zonal mean sea-surface temperatures. The body sizes exhibited by the HD are positively offset from the MD, which may be the effect of human taxonomists preferentially identifying larger specimens, as smaller individuals are more difficult to classify. The HD is comprised of only individuals with high-quality species labels (see Materials & Methods); since smaller individuals are more likely to engender disagreement between human classifiers, they have a higher chance of being excluded from the HD.

Fig. 5 shows the Jaccard community dissimilarity indices for all sitewise comparisons for the HD (a), MD (b), and CD (c). In the HD, the sites that appear to have relatively elevated Jaccard indices are IPE.08295, IPE.08320, IPE.08248, and IPE.08171. These sites also show relatively elevated Jaccard index values the MD (though much less markedly for IPE.08248). In particular, sites IPE.08295 and IPE.08171 appear to be the most distinct from one another in all datasets. IPE.08295 is the northernmost site in our sample at 67.65°N, and is highly dominated by *N. pachyderma* and *N. dutertrei* (Fig. 4). In contrast, IPE.08171 is a tropical site (12.973°N) dominated by *Globigerinoides ruber* and *G. sacculifer* (Fig. S5a), and also contains many large individuals (outlier site in Fig. 3b-c). In effect, these two sites represent the poleward and equatorial community extremes, respectively. IPE. 08320 is the site with the second-most northern latitude (54.067°N) and thus exhibits higher Jaccard indices against lower latitude sites (and a relatively low Jaccard index against IPE.08295). Finally, IPE.08248 is the southernmost site in our sample (45.675°S), and is relatively dominated by *N. pachyderma*. We note that the Jaccard indices for IPE.08248 in the MD are not as elevated as those for the HD, as highlighted in Fig. 5d, which shows the absolute differences of the sitewise Jaccard indices for the HD (J_HD_) and the MD (J_MD_). This may be caused by the commonality bias exhibited by human classifiers, as examination of the species abundances at IPE.08248 (see Supplementary Materials) reveals that the MD contains eight rare species that are unrepresented in the HD. (The converse, where a rare species is represented in the HD and not the MD, does not occur at this site.) This may also explain why the difference in Jaccard indices for the comparison between site IPE.08295 and IPE.08248 is high – there are three rare species represented in the MD and not the HD for site IPE.08295, which are also represented in the MD but not the HD for site IPE.0248.

**Fig. 5.**
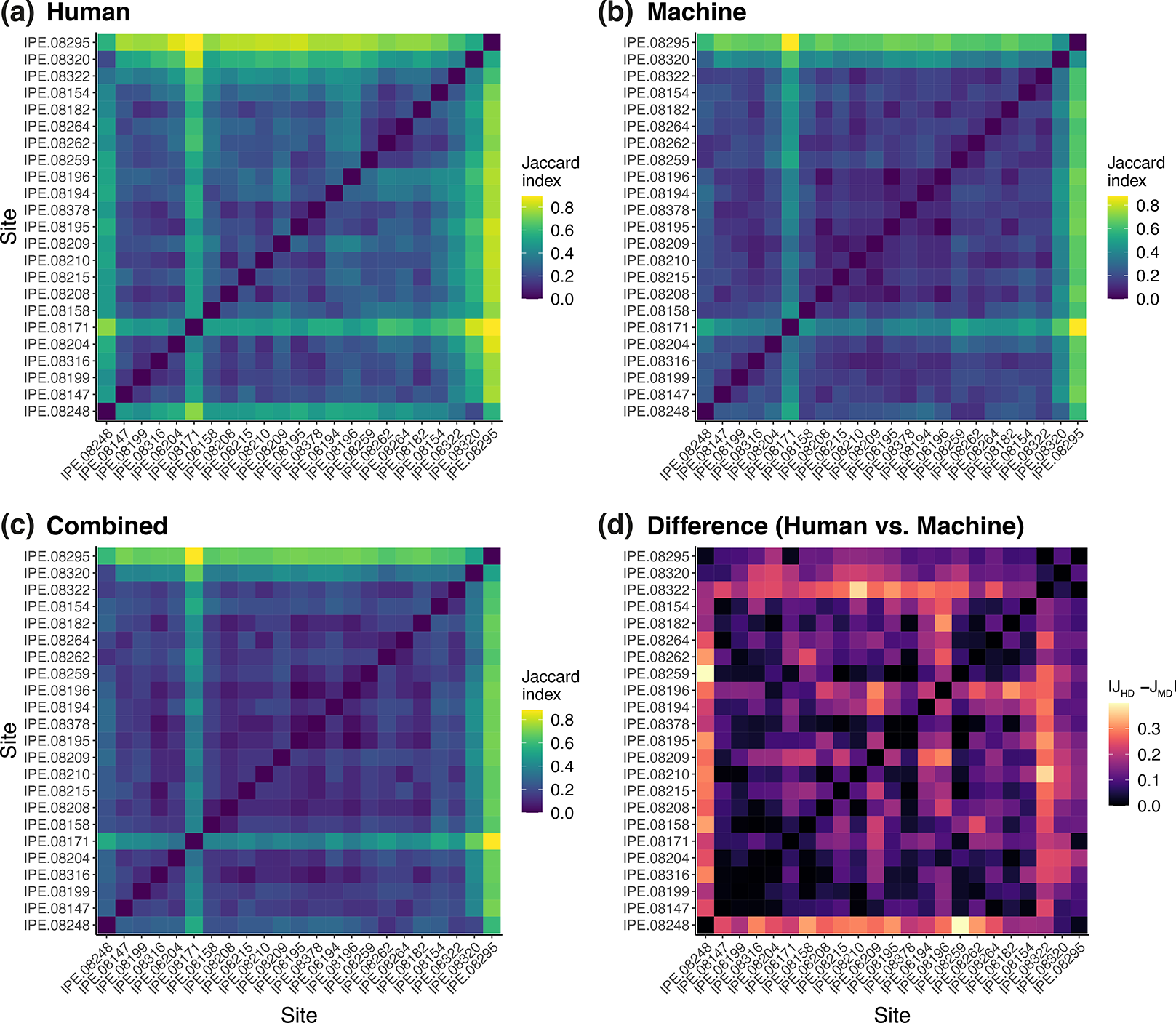
ß-diversity heatmaps. The Jaccard community indices calculated for each sitewise comparison for all 23 sites sampled in this study are shown for the (a) human dataset, (b) machine dataset, and (c) combined dataset. The absolute element-wise differences between the Jaccard indices for the human (J_HD_) and machine datasets (J_MD_) is shown in (d). Axes are ordered from southernmost (left/bottom) to northernmost (right/top) sites.

Finally, we calculate the assemblage comparison diversity estimates proposed by Chao et al. (2020) to compare diversity patterns across assemblages. In general, the patterns exhibited by the HD and the MD are similar as the order *q* (which quantifies sensitivity to species abundances in our diversity metrics; see Chao (2020)) changes for the sample completeness (Fig. 6a), diversity (Fig. 6c), and evenness (Fig. 6e) profiles. The sample-size- and coverage-based rarefaction/extrapolation curves are also similar for the HD and the MD across the values of *q* (e.g., species diversity decreases relatively as *q* increases). However, in every diversity metric, we observe that the diversity in the MD is higher than in the HD, except in some cases at low values of *q* (e.g., when *q* = 0 in the size-based rarefaction curve [Fig. 6b] — that is, when relative abundances are not taken into account). Evenness also exhibits a positive offset for all values of *q* for the MD vs. the HD (Fig. 6e). These results corroborate our other analyses and previous studies demonstrating the rarity bias present in human-identified datasets, and suggests that the offset in species richness between the HD and the MD (Fig. 3a) is not an artefact resulting from differences in sample size or coverage between the two datasets.

**Fig. 6.**
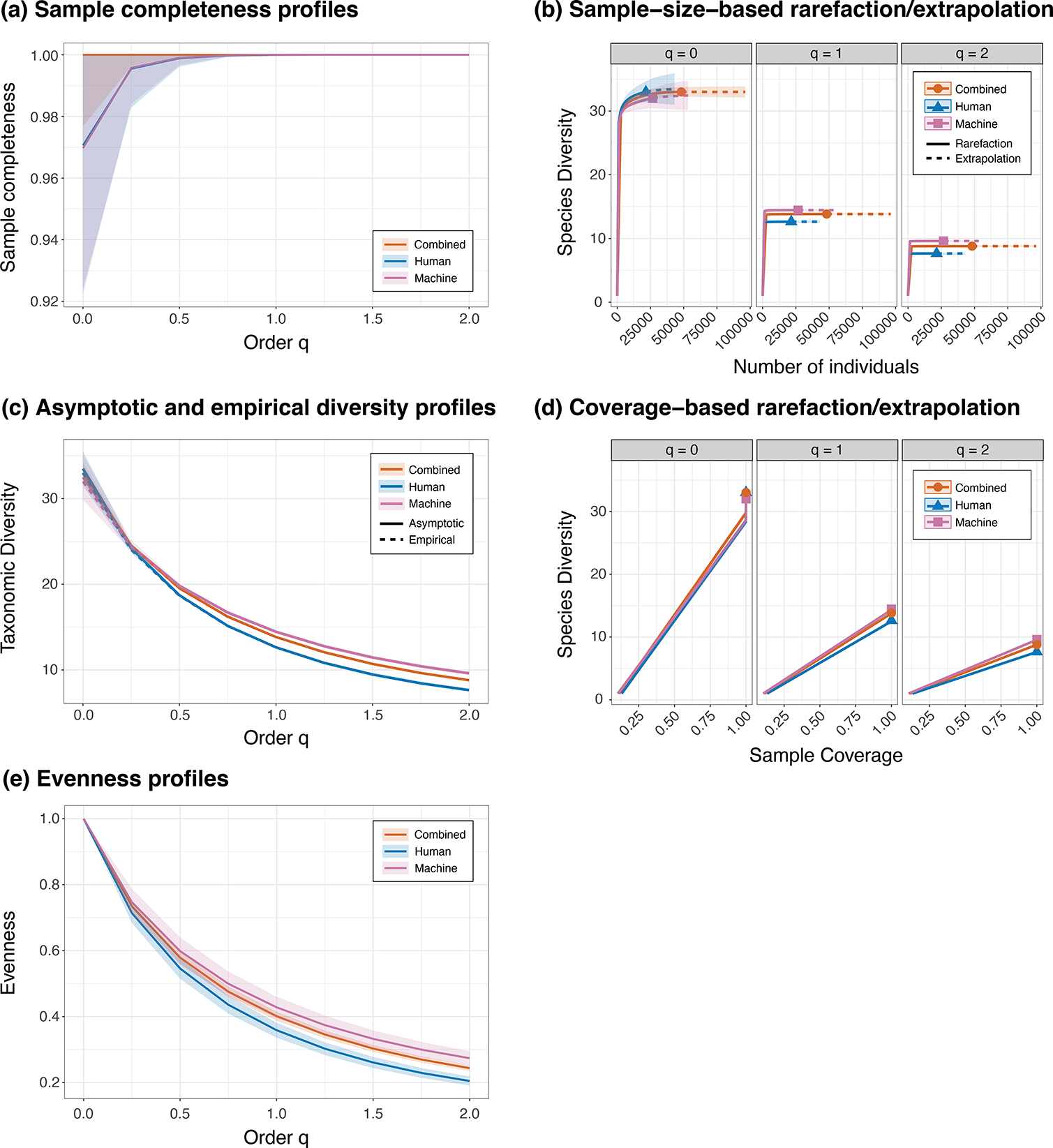
Assemblage comparison diversity estimates. We treat the HD (N_HD_=21,540), MD (N_MD_=26,654), and CD (N_CD_=48,194) as different assemblages and calculate the (a) sample completeness profiles, (b) sample-sized-based rarefaction/extrapolation curves, (c) asymptotic and empirical diversity profiles, (d) coverage-based rarefaction/ extrapolation curves, and (e) evenness profiles. Shaded regions represent 95% confidence intervals calculated using bootstrapping with 50 replicates.

## Discussion

We have shown that CNN-driven community ecology is not only feasible and accurate, but also avoids biases that can affect datasets generated by human classifiers that may result in misleading conclusions about biodiversity patterns. Aside from the commonality and sampling biases discussed above, human classifiers are also subject to practical limitations that a well-trained CNN could avoid. For instance, smaller individuals are harder to identify for human classifiers, and thus human-identified datasets are more likely to omit smaller members of a community (as exhibited by the latitude-size relationship curves in Fig. 3b-c). Machine classifiers are not affected by this size bias, assuming the images used are taken at high enough resolution. We note that the omission of smaller specimens by human classifiers can occur both at sample preparation stage and at identification stage, particularly given that current standard practices for building planktonic foraminifer inventories involves hand-picking individuals and affixing them to microscope slides. Thus, in order to truly take advantage of computer vision methods for large-scale studies of community ecology and biodiversity, there must be concurrent advances in high-throughput imaging and digitisation of specimens that limit or remove the likelihood of human biases affecting dataset composition.

## Supporting information

Rank abundance data (CSVs)

Normalized abundance maps (PDFs)

Sitewise species distributions (PDFs)

Machine-assigned species labels

Shape measurements for human- and machine- labeled foraminifera

## Acknowledgements

We would like to thank Srinath Krishnan for discussion and assistance with netCDF processing. AYH is supported by Vetenskapsrådet (VR) grant 2020-03515.

