## Supplementary Figures S1-S5 and Table S1 for "Automated community ecology using deep learning: a case study of planktonic foraminifera"

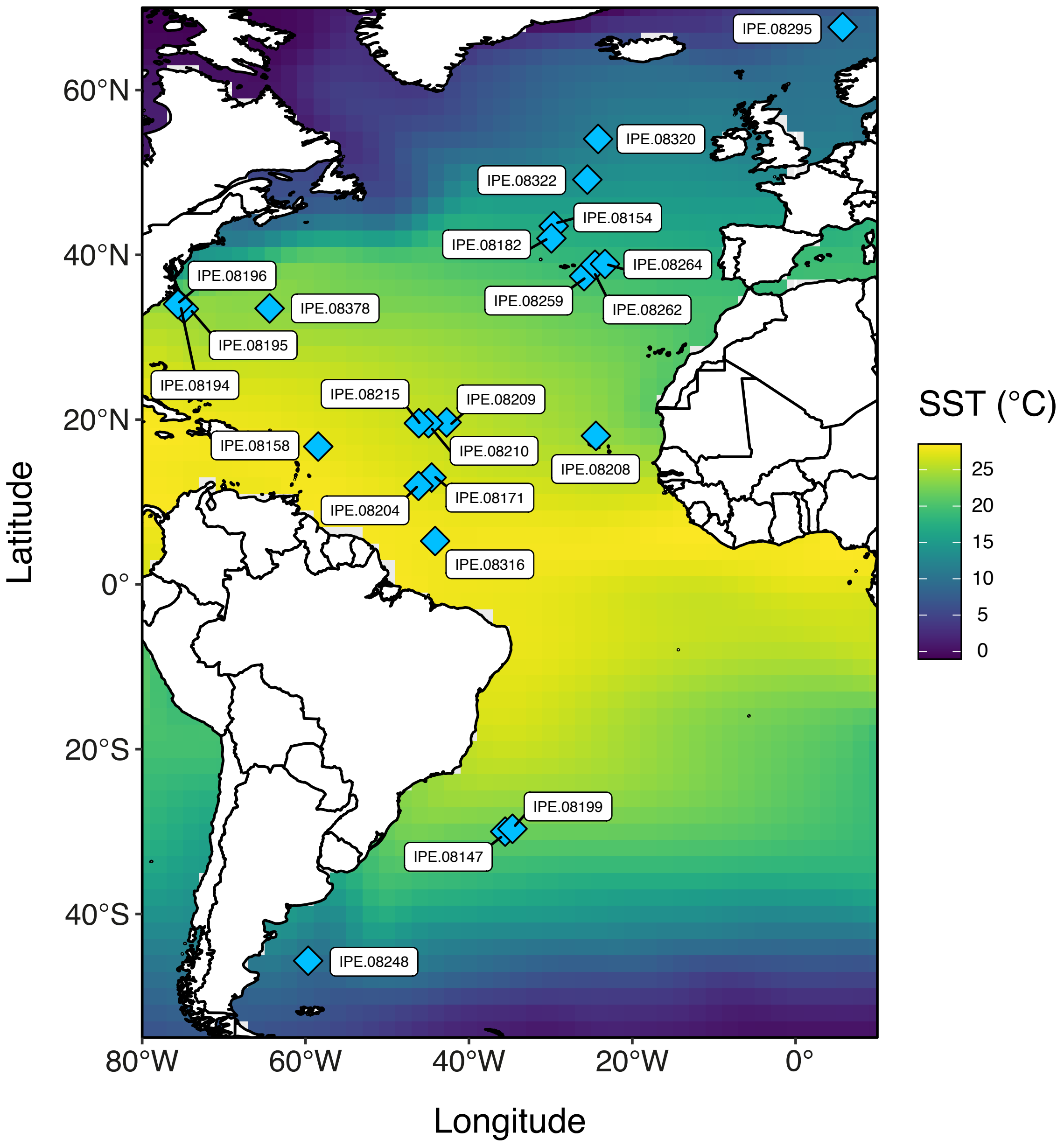


**Fig. S1.** *Site map of Atlantic coretop sites with good preservation sampled in this study*. Sea-surface temperatures (SSTs) shown are averaged over all 2° x 2° grid data points in ERSSTv5 from January 2000 to December 2018 between 54°S-70°N and 80°W-10°E.

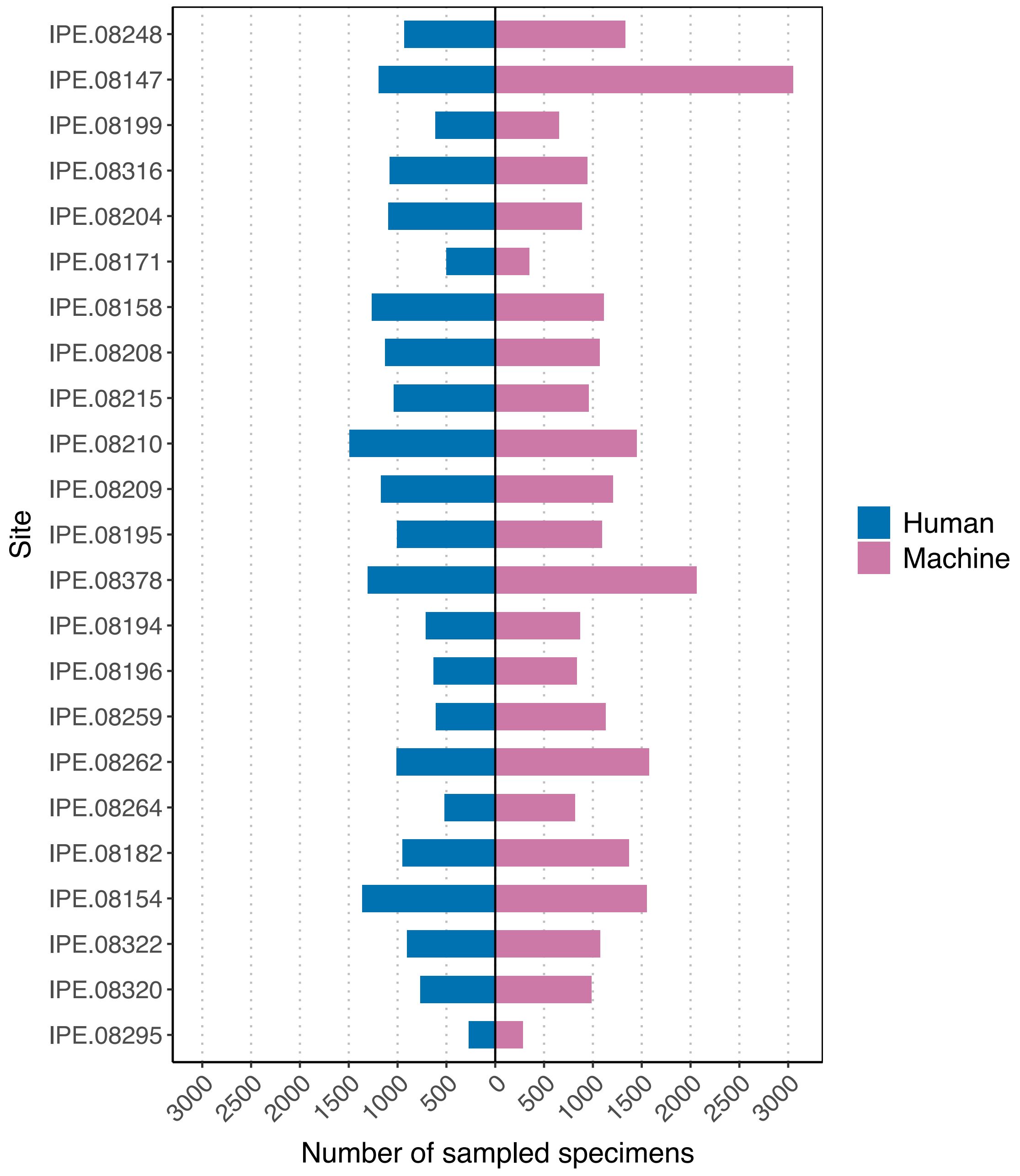


**Fig. S2.** *Number of samples per site by dataset*. Sites are listed in order of southernmost (top; 45.675˚S) to northernmost (bottom; 67.65˚N) latitude.

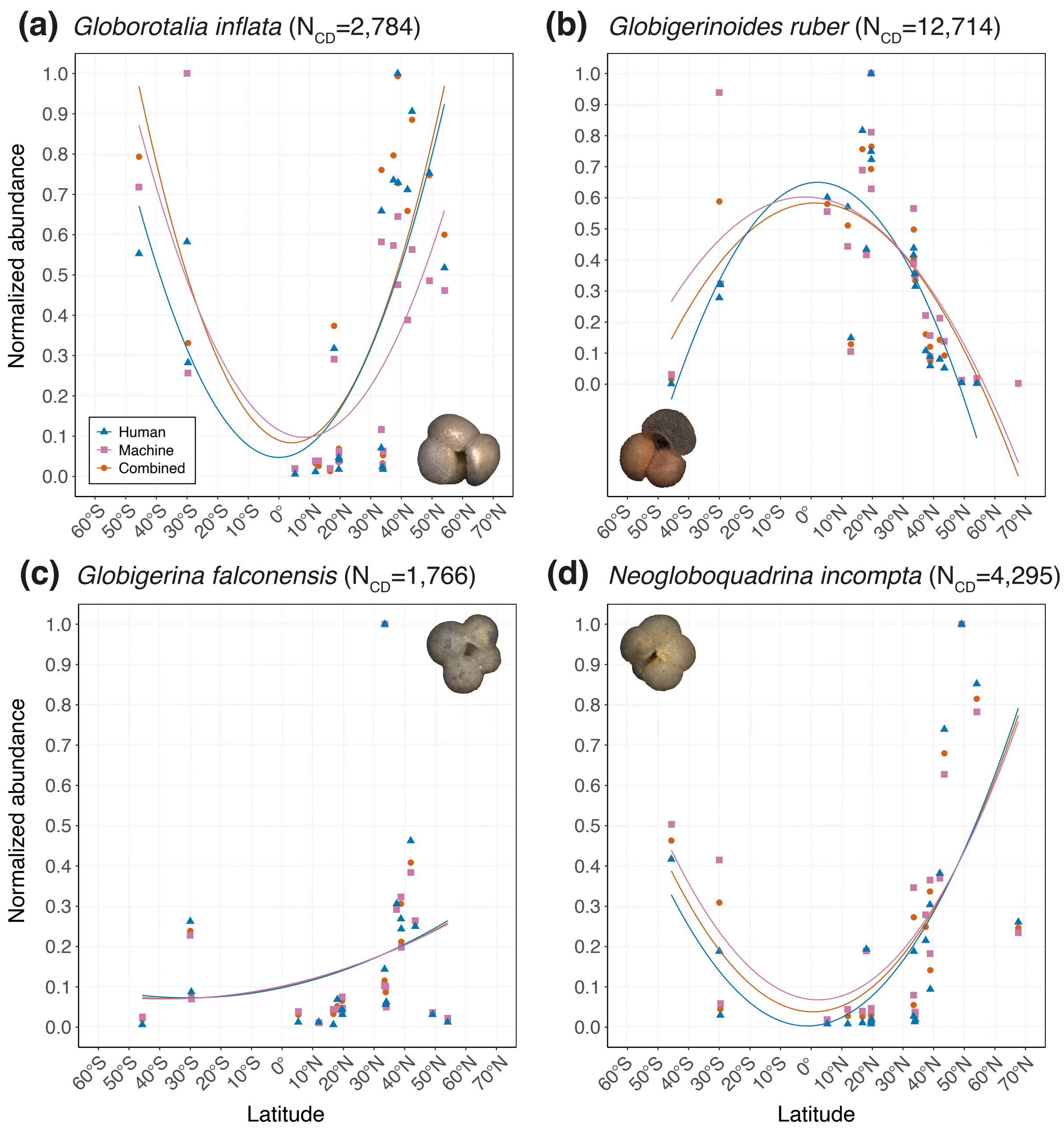


**Fig. S3.** *Species-specific latitudinal normalised abundance curves.* These alternate visualisations of the normalised abundance data show clearly the latitudinal patterns in normalised abundance observed. Normalised abundance is plotted against latitude, with the curves representing best-fit second-order polynomial curves for each dataset. (a) *Globorotalia inflata* shows a clear poleward increase in normalised abundance in all datasets. (b) *Globigerinoides ruber* shows the opposite pattern, with normalised abundance decreasing towards the pole for all datasets. (c) Some taxa, such as *Globigerina falconensis*, show neither pattern; the slight upward trajectory towards the northern latitudes may reflect a true biological pattern or may result from geographical sampling bias in the northern hemisphere. (d) *Neogloboquadrina incompta* exhibits a pattern that may be explained by sampling bias, whereby the discrepancy between the HD, MD, and CD decreases as the sites sampled move from the southern to the northern latitudes. Species exemplar images sourced from the Endless Forams database.

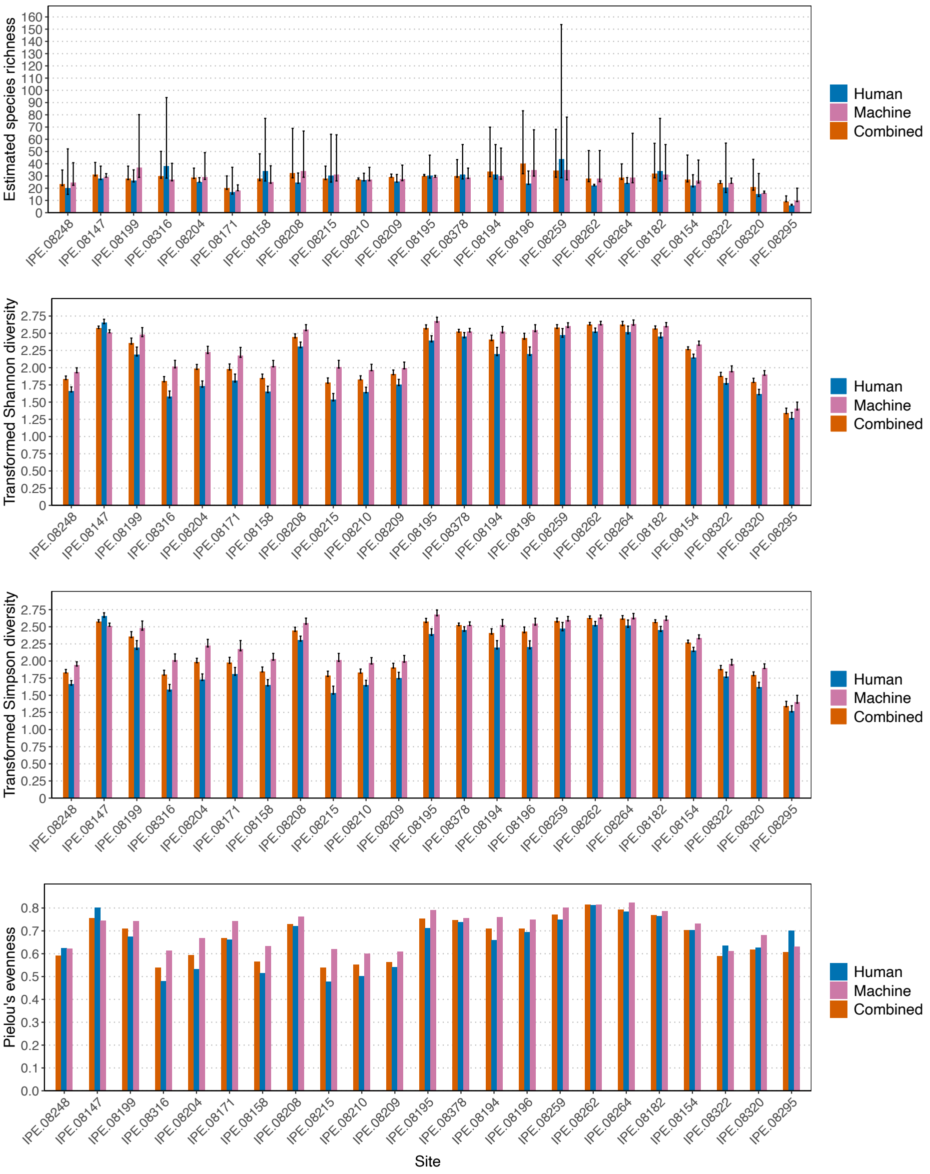


**Fig. S4.** *α-diversity metrics*. (a) Estimated species richness (*sensu* Chao 1985; 1987) with 95% confidence intervals; (b) Transformed Shannon diversity with 95% confidence intervals; (c) Transformed Simpson diversity with 95% confidence intervals; (d) Pielou's evenness. Sites are ordered from southernmost latitude (left; 45.675°S) to northernmost latitude (right; 67.65°N).

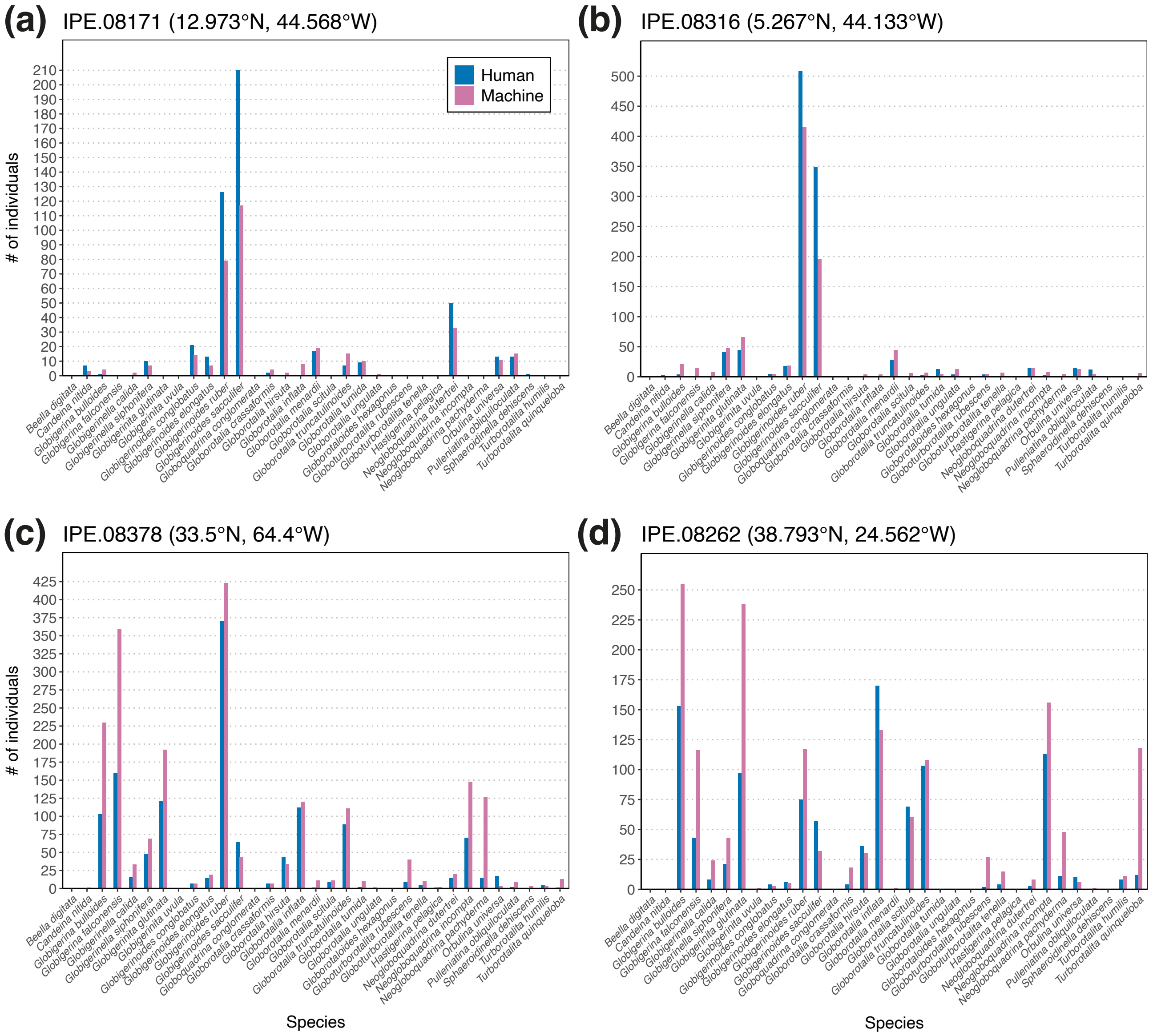


**Fig. S5.** *Species distribution by site*. Distribution profiles from exemplar sites, with y-axis showing number of representatives of a species for human and machine datasets. (a-b) show the recurring pattern where, in sites dominated by only a few species, the number of those species in the HD tends to be larger than in the MD. In contrast, (c-d) show that in sites where the distribution is more evenly spread across a number of species, the number of individuals in the MD tends to outnumber those in the HD for a particular species, as would be expected given that the MD contains ~5,000 more specimens than the HD. These examples illustrate the commonality bias observed in human classifiers (see text and Hsiang et al. (2019)).

**Table S1.** *Assemblage comparison metrics for Hill number special cases.*

| Analysis | Human | | | Machine | | | Combined | | |
|---|---|---|---|---|---|---|---|---|---|
| | $q = 0$ | $q = 1$ | $q = 2$ | $q = 0$ | $q = 1$ | $q = 2$ | $q = 0$ | $q = 1$ | $q = 2$ |
| (a) Sample completeness | 0.9706 | 0.9999 | 1.000 | 0.9697 | 0.9999 | 1.000 | 1.000 | 1.000 | 1.000 |
| (b,c) Sample-sized-based rarefaction/extrapolation | | | | | | | | | |
| (b,c.1) Asymptotic | 33.4999 | 12.6431 | 7.6485 | 32.4999 | 14.4560 | 9.6075 | 33.000 | 13.8425 | 8.8013 |
| (b,c.2) Empirical | 33.000 | 12.6334 | 7.6461 | 32.000 | 14.4473 | 9.6044 | 33.000 | 13.8378 | 8.7999 |
| (d) Coverage-based rarefaction/extrapolation[1] | 33.000 | 12.6334 | 7.6461 | 32.000 | 14.4473 | 9.6044 | 32.9967 | 13.8376 | 8.7998 |
| (e) Evenness[2] | 1.000 | 0.3589 | 0.2050 | 1.000 | 0.4282 | 0.2739 | 1.000 | 0.4014 | 0.2439 |

[1] At maximum standardized coverage of 99.99% (see Chao *et al.* 2020)

[2] At 99.99% coverage
