## Supplementary figures and images for "Automated community ecology using deep learning: a case study of planktonic foraminifera"

### beella_digitata_global_range_plot.pdf

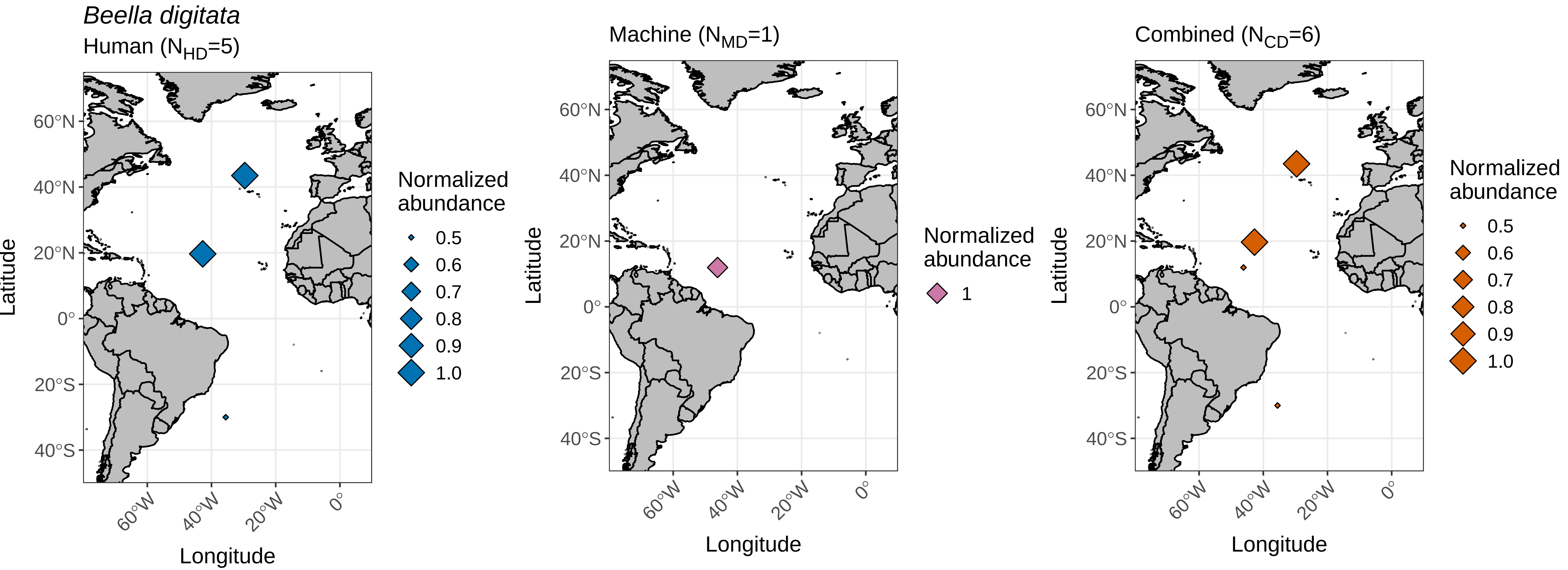

### candeina_nitida_global_range_plot.pdf

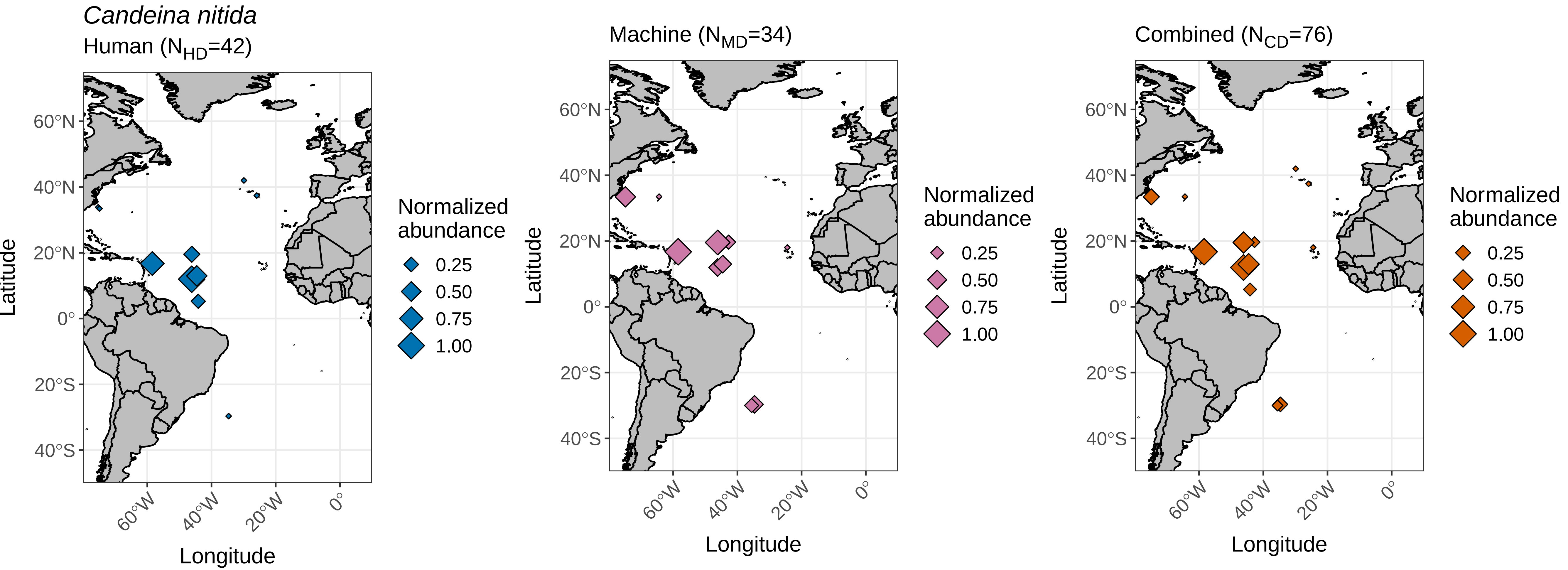

### globigerina_bulloides_global_range_plot.pdf

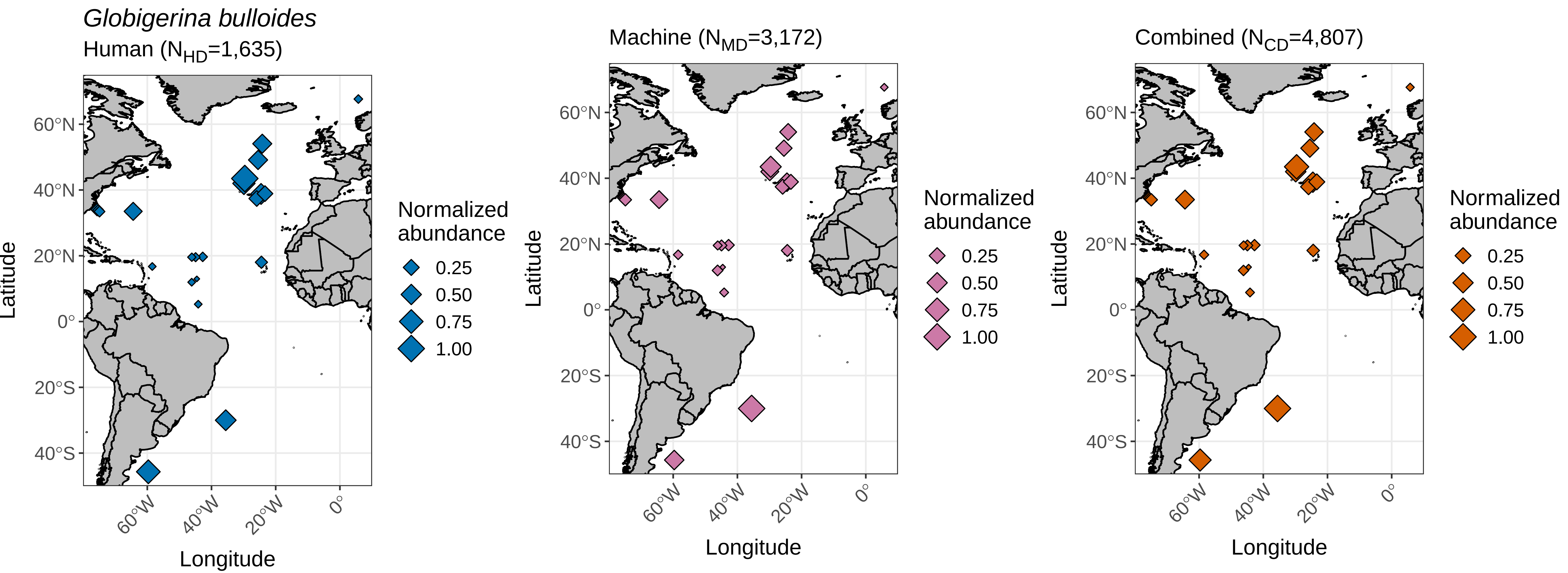

### globigerina_falconensis_global_range_plot.pdf

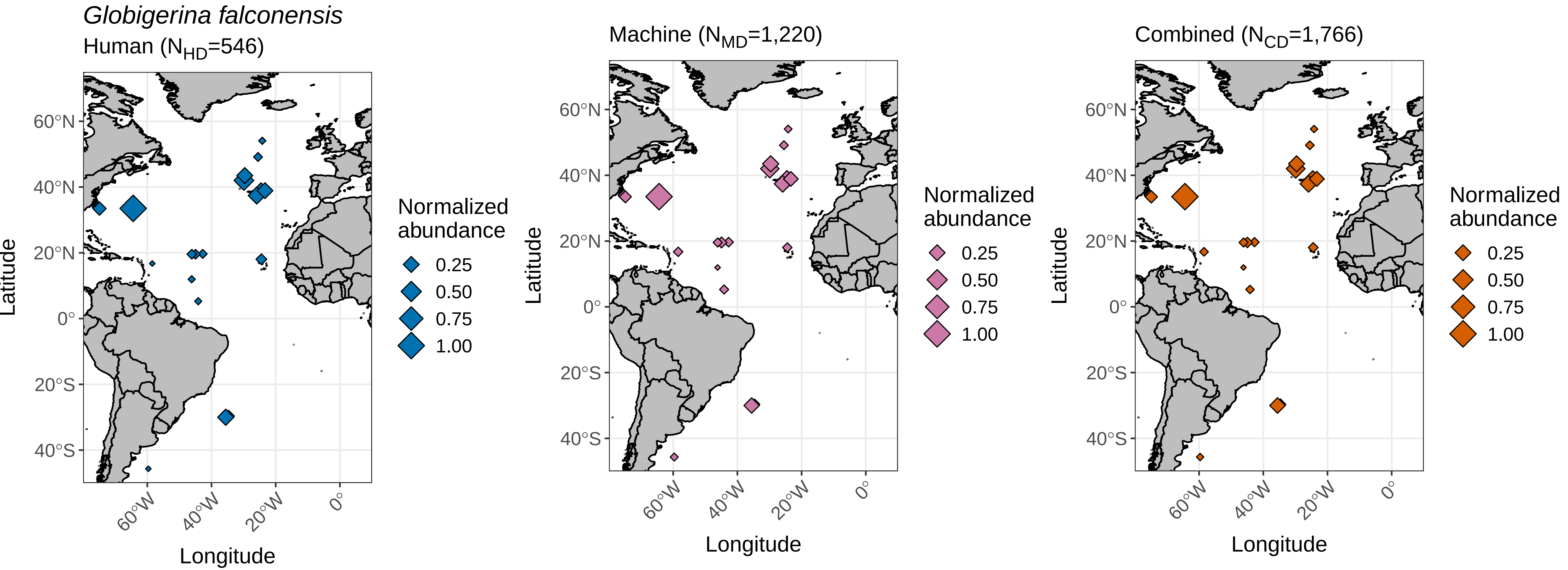

### globigerinella_calida_global_range_plot.pdf

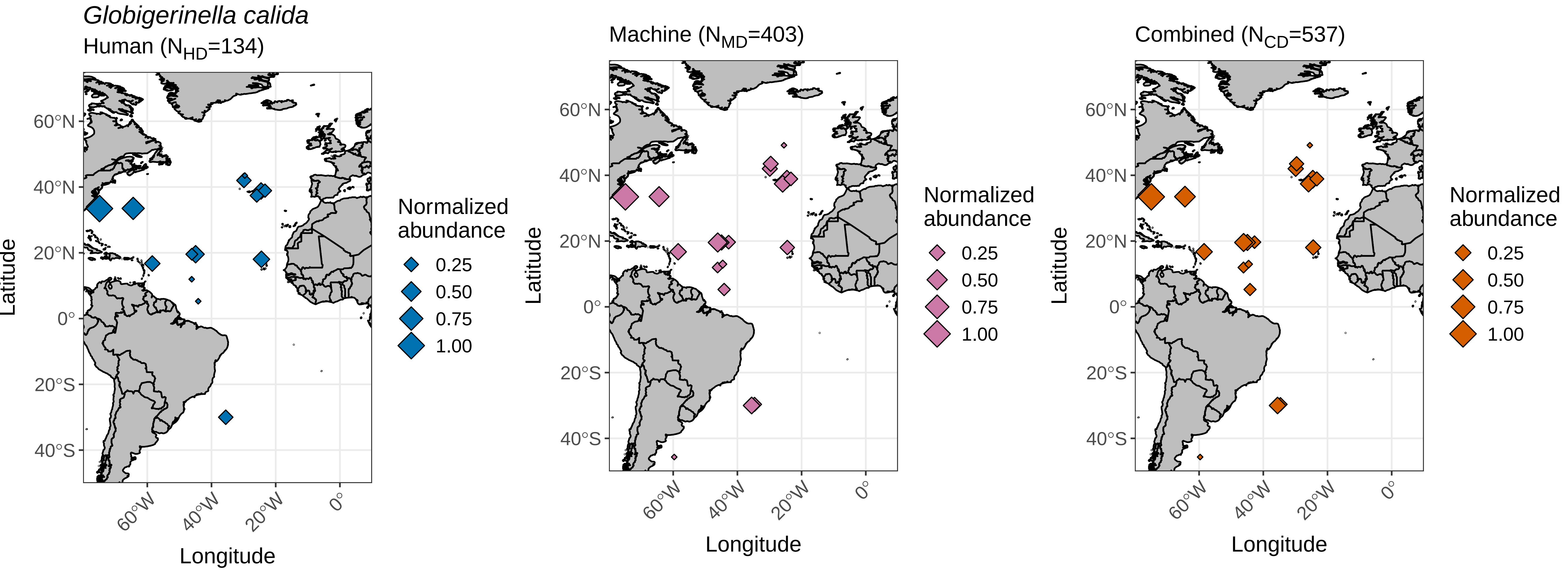

### globigerinella_siphonifera_global_range_plot.pdf

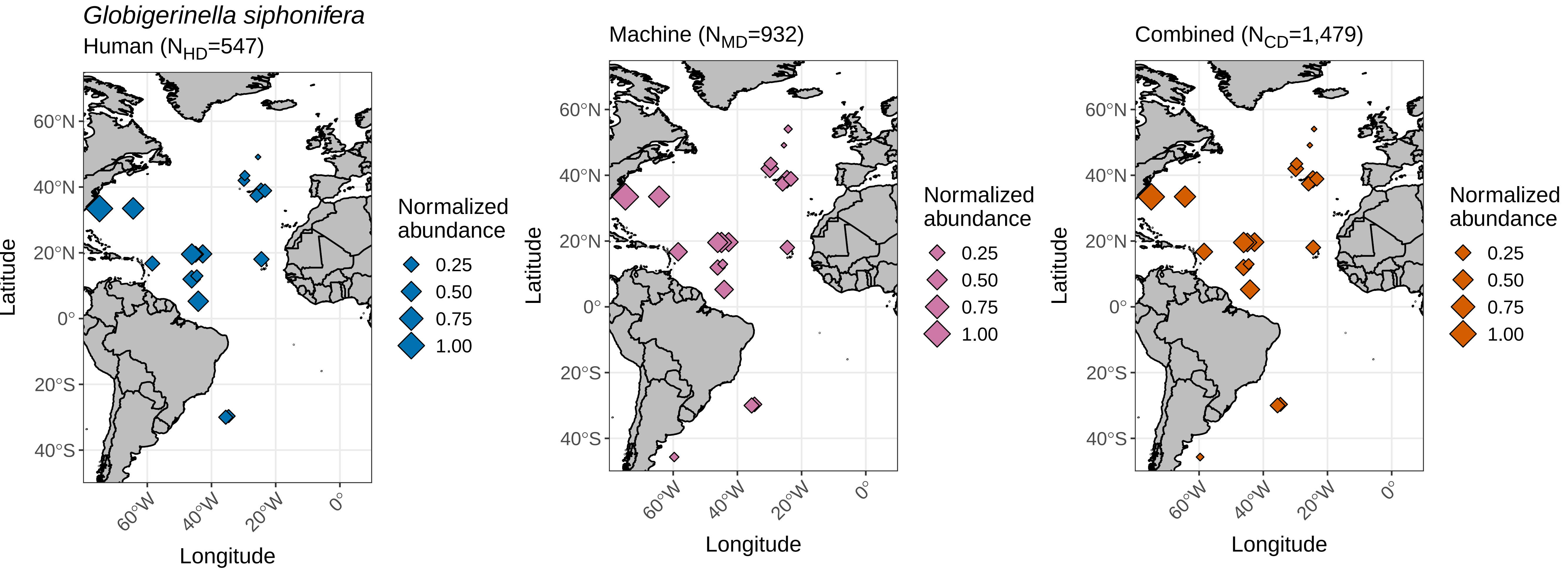

### globigerinita_glutinata_global_range_plot.pdf

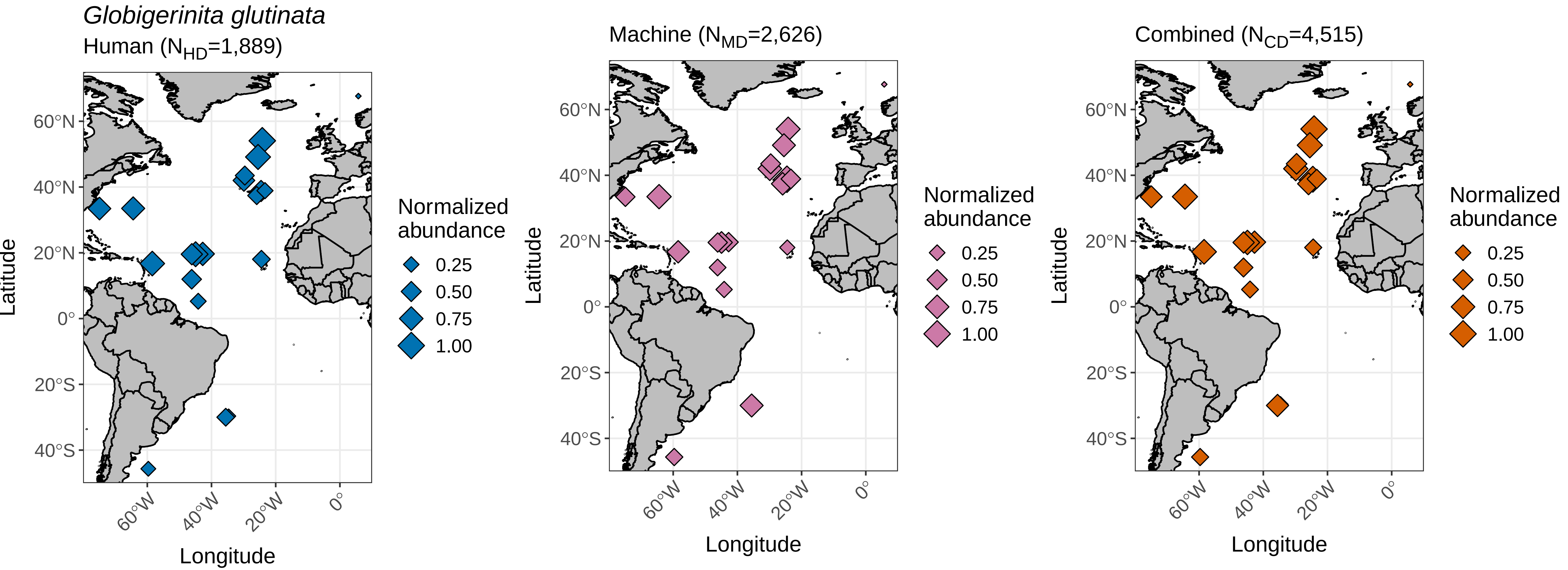

### globigerinita_uvula_global_range_plot.pdf

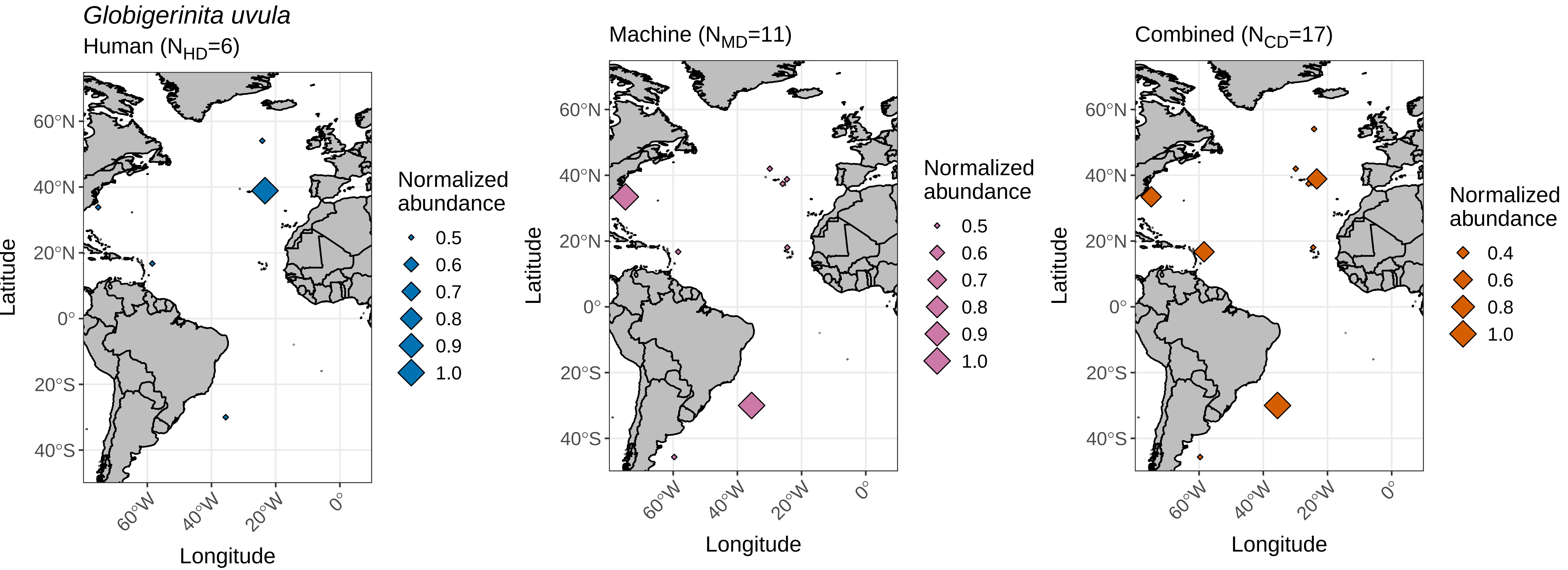

### globigerinoides_conglobatus_global_range_plot.pdf

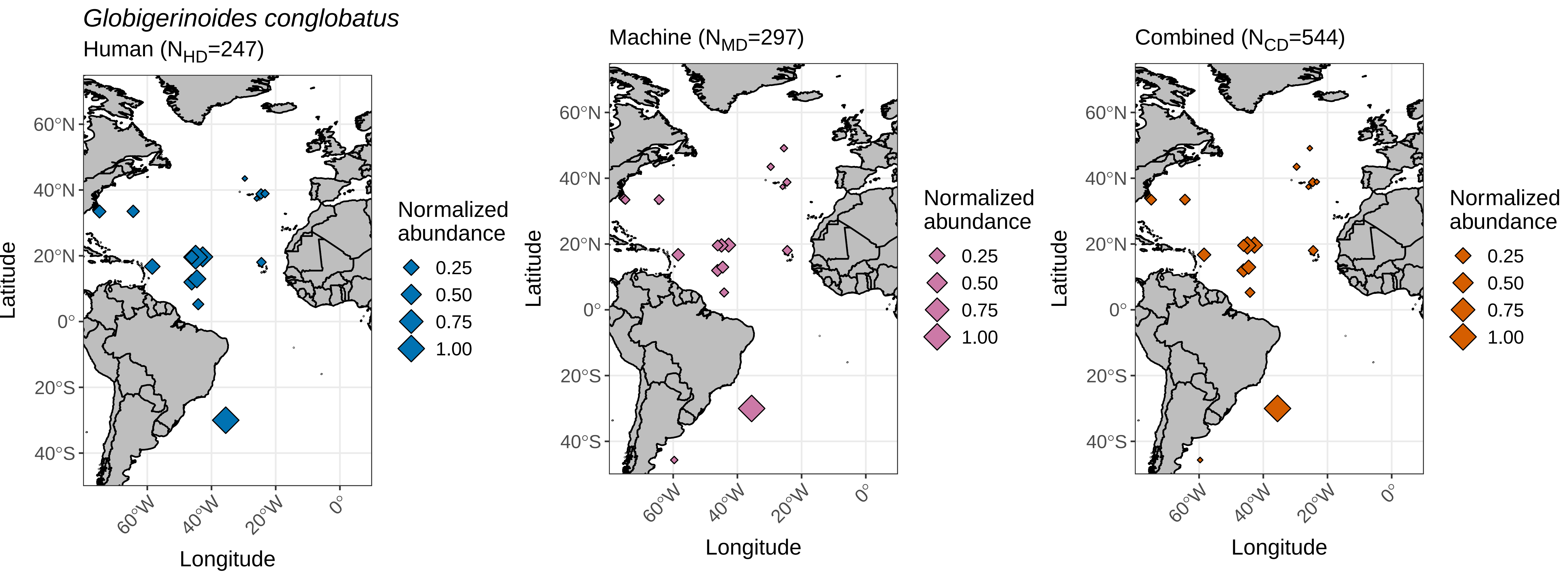

### globigerinoides_elongatus_global_range_plot.pdf

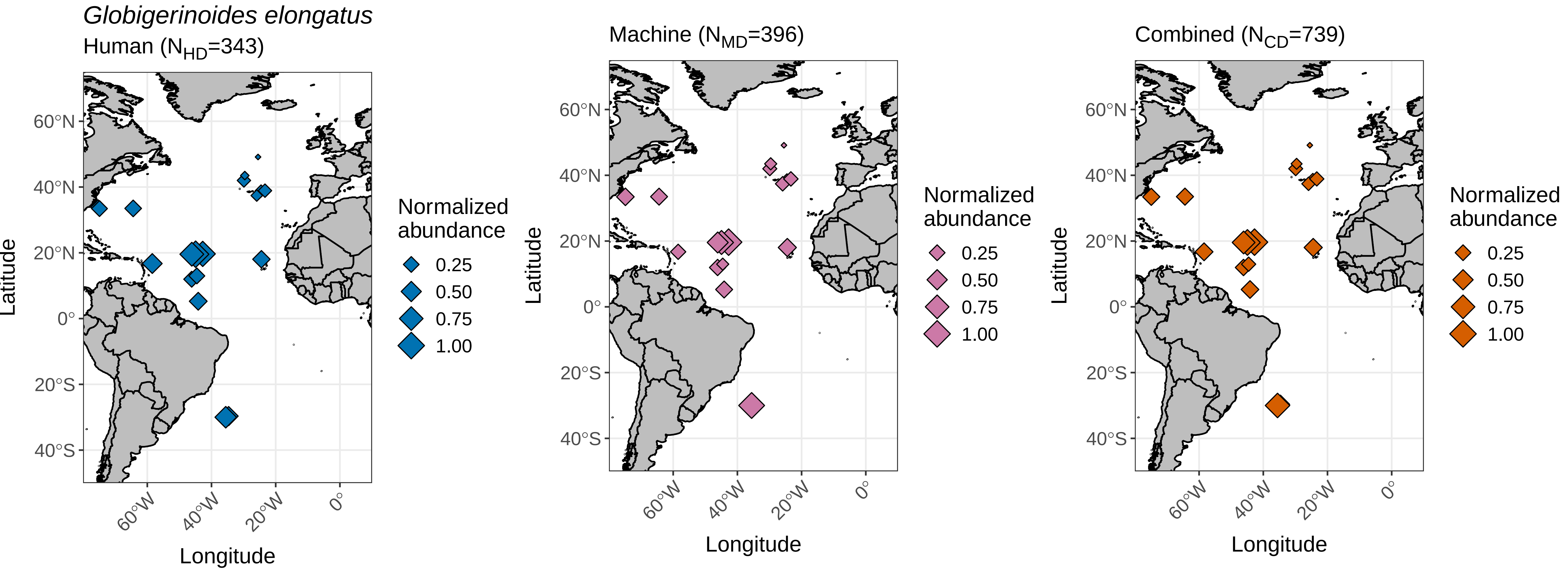

### globigerinoides_ruber_global_range_plot.pdf

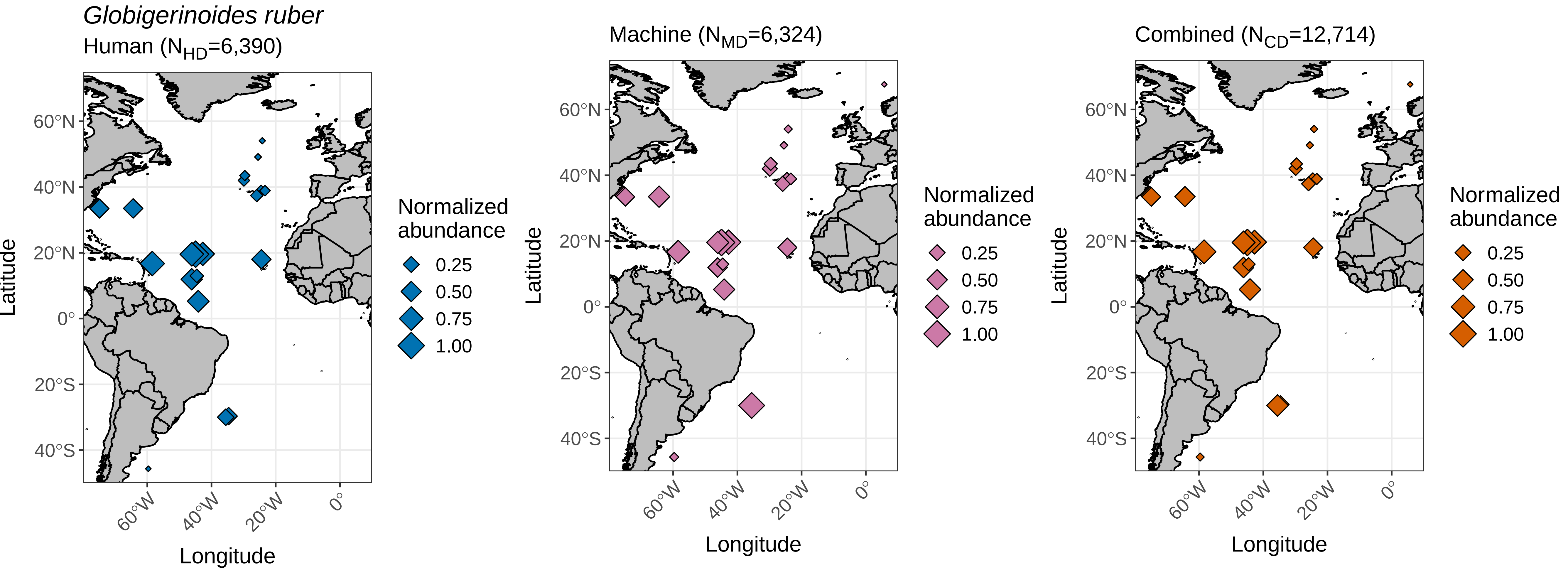

### globigerinoides_sacculifer_global_range_plot.pdf

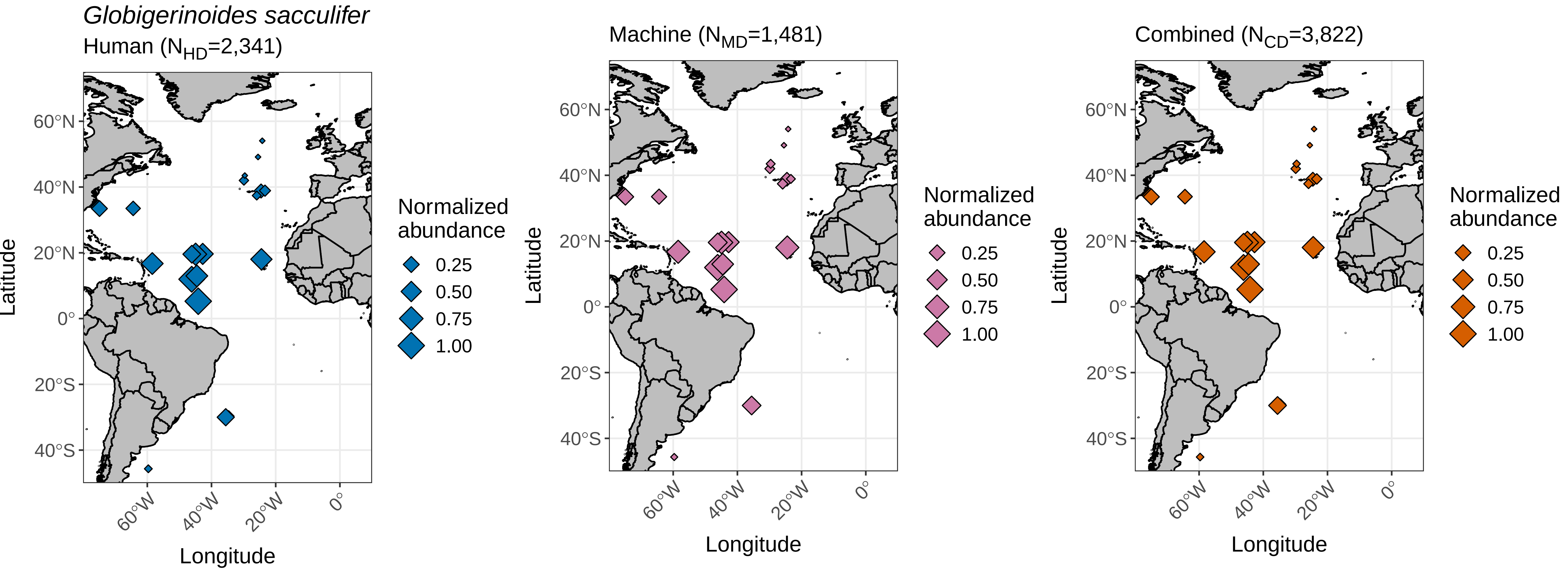

### globoquadrina_conglomerata_global_range_plot.pdf

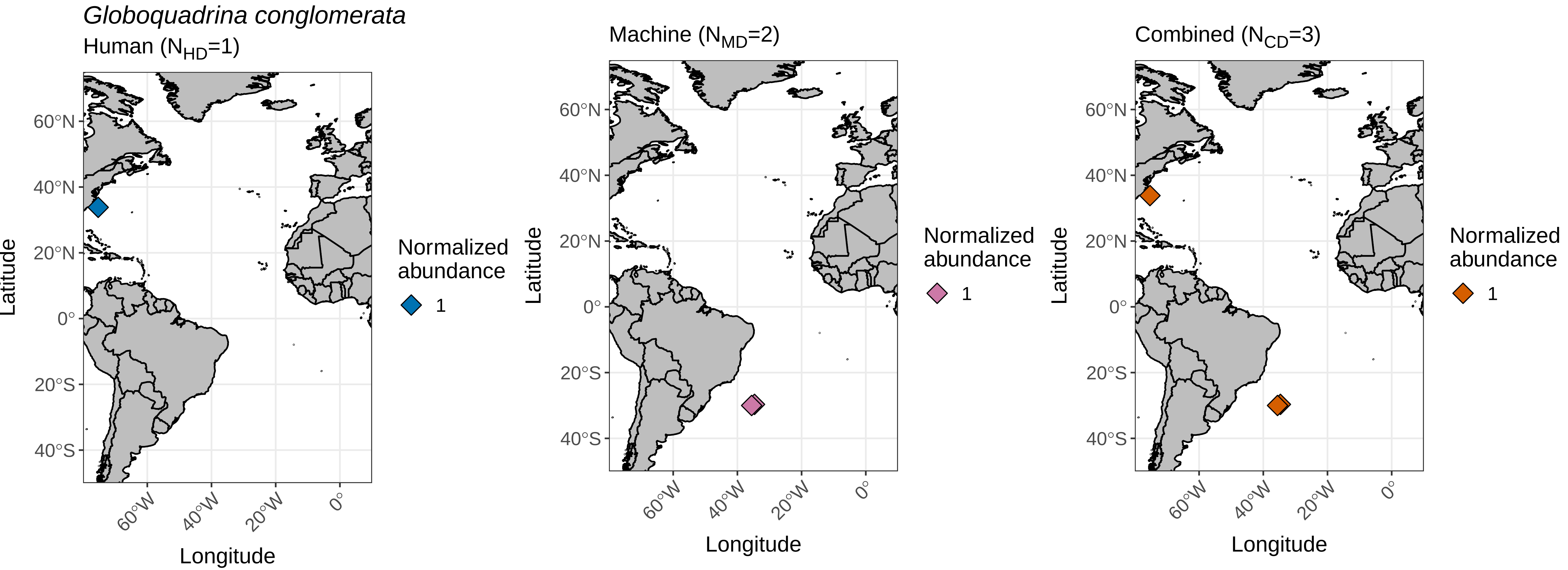

### globorotalia_crassaformis_global_range_plot.pdf

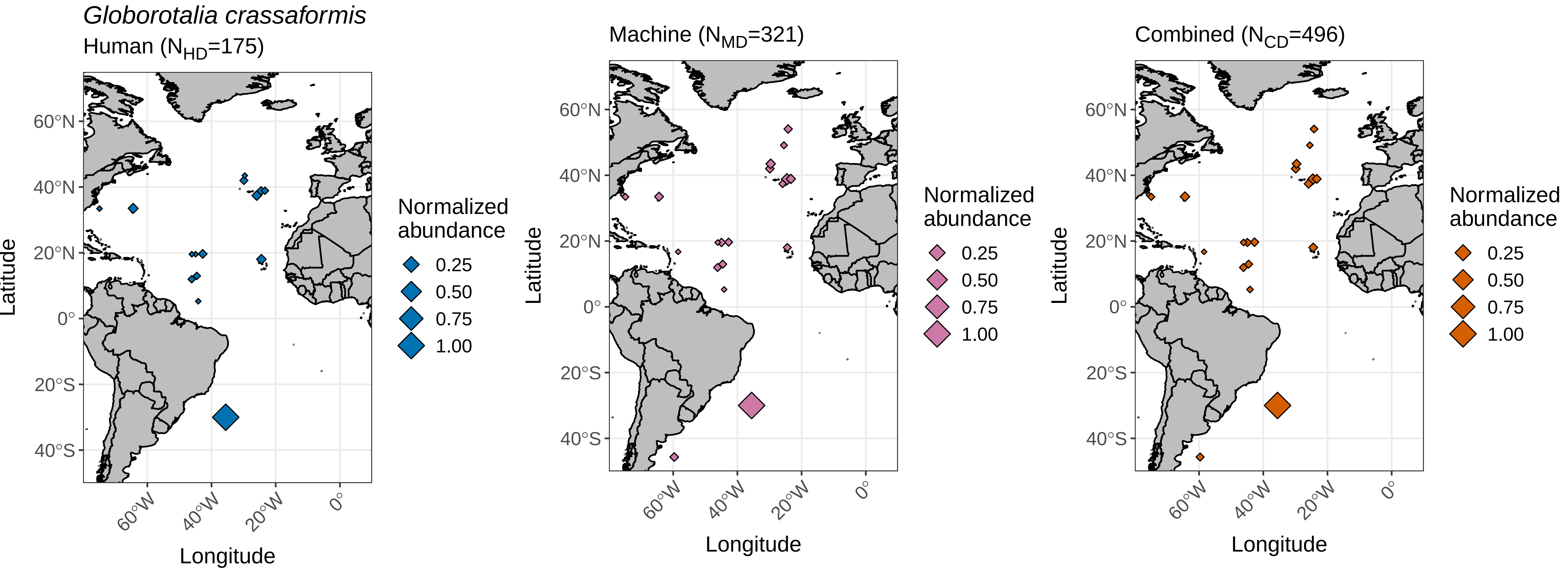

### globorotalia_hirsuta_global_range_plot.pdf

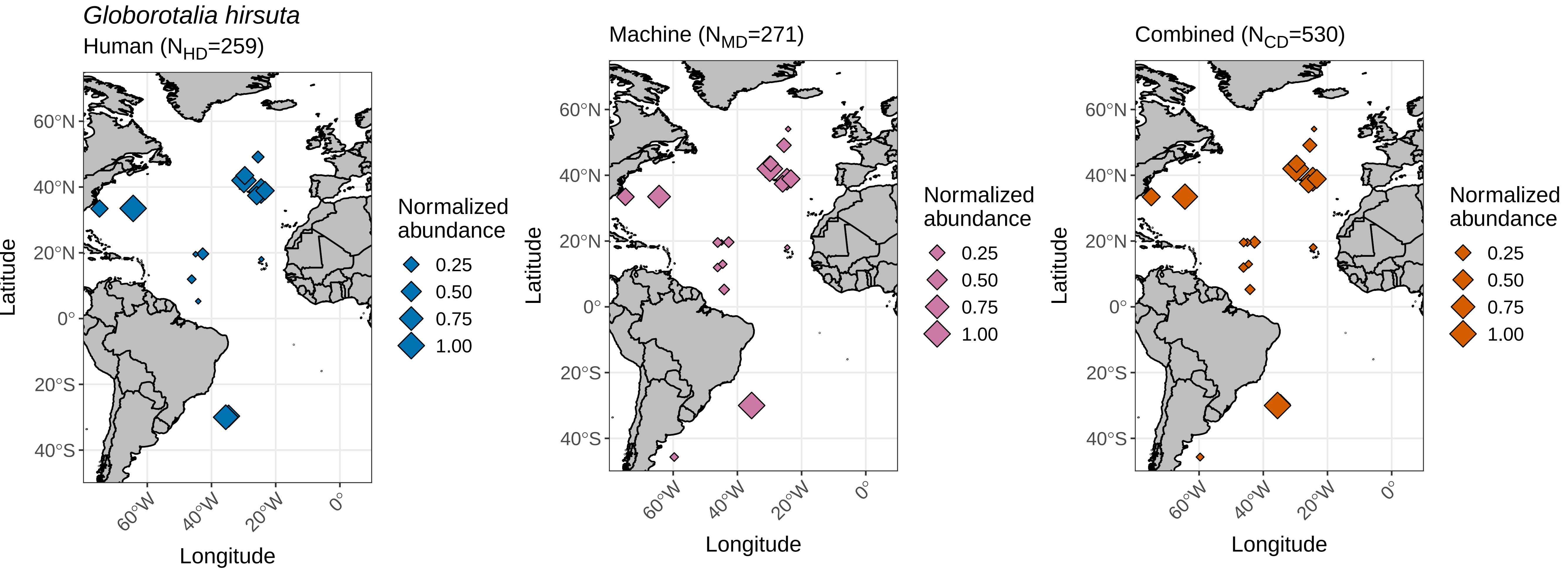

### globorotalia_inflata_global_range_plot.pdf

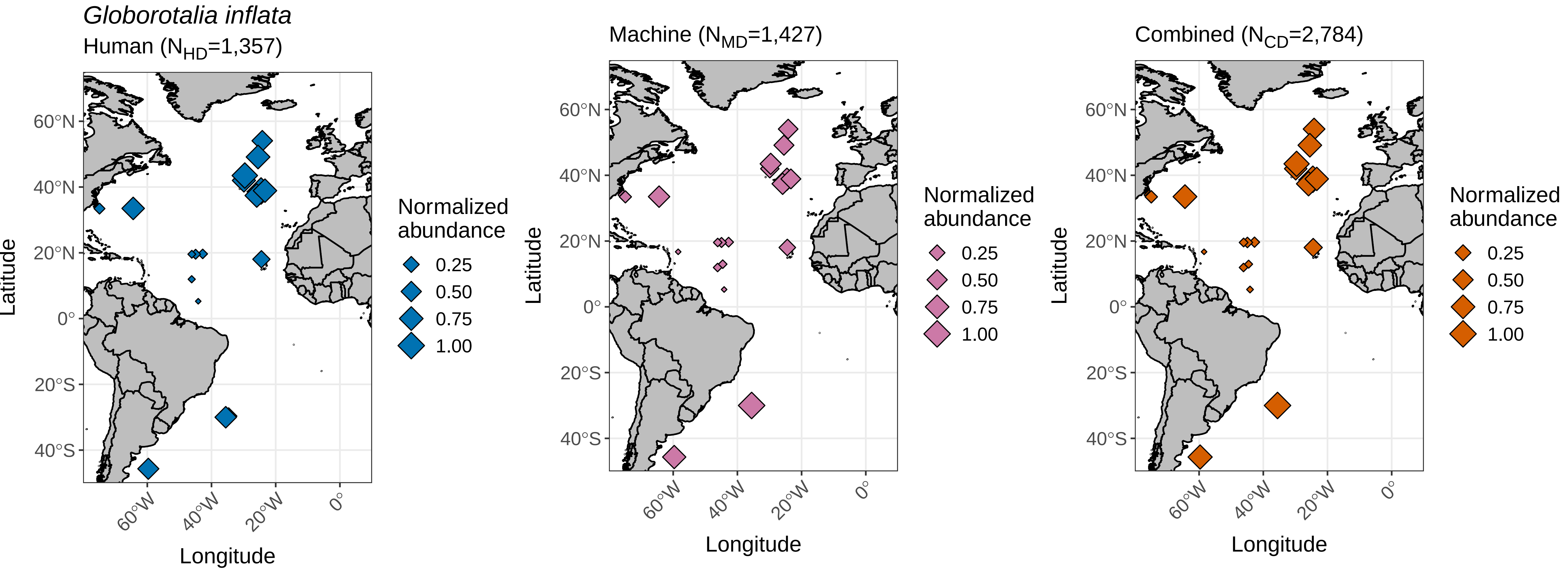

### globorotalia_menardii_global_range_plot.pdf

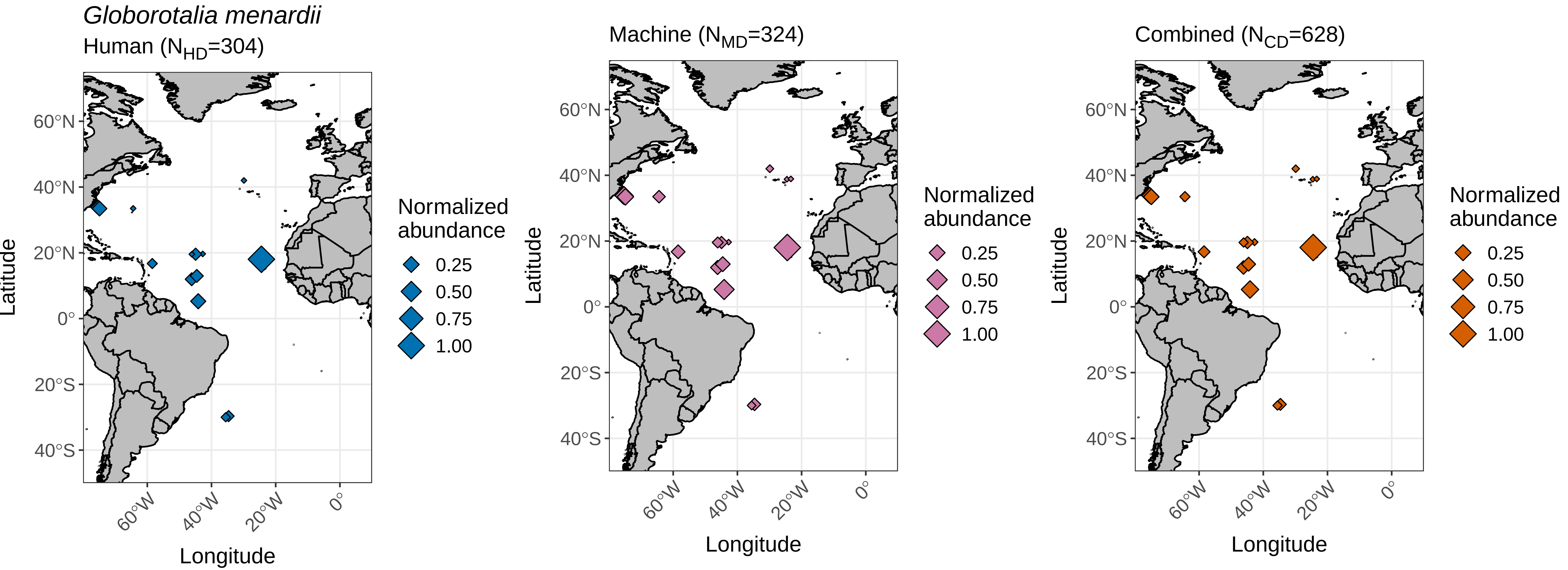

### globorotalia_scitula_global_range_plot.pdf

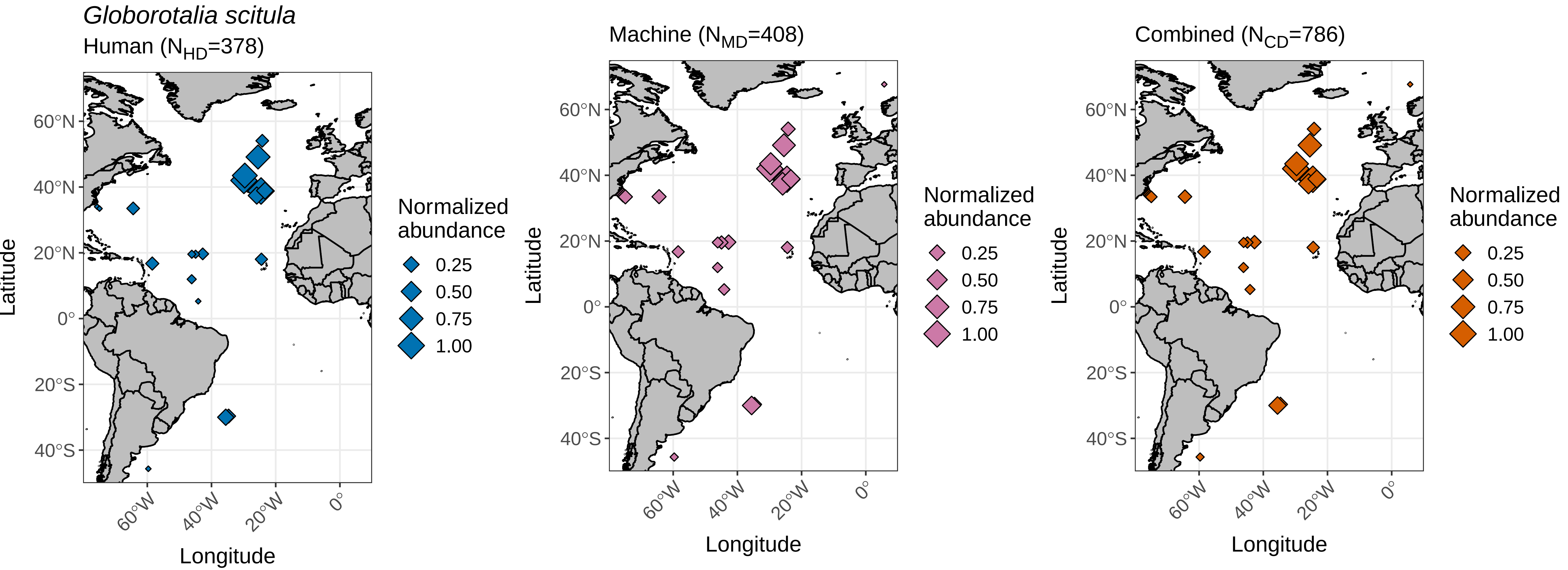

### globorotalia_truncatulinoides_global_range_plot.pdf

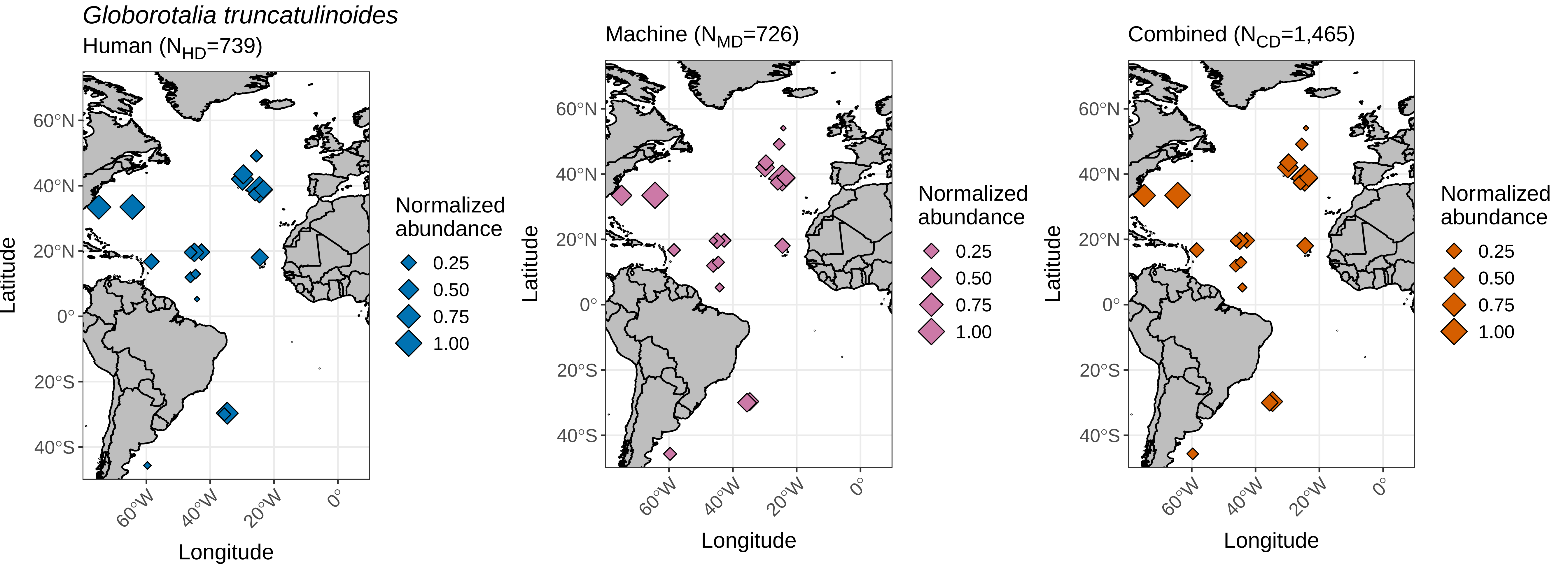

### globorotalia_tumida_global_range_plot.pdf

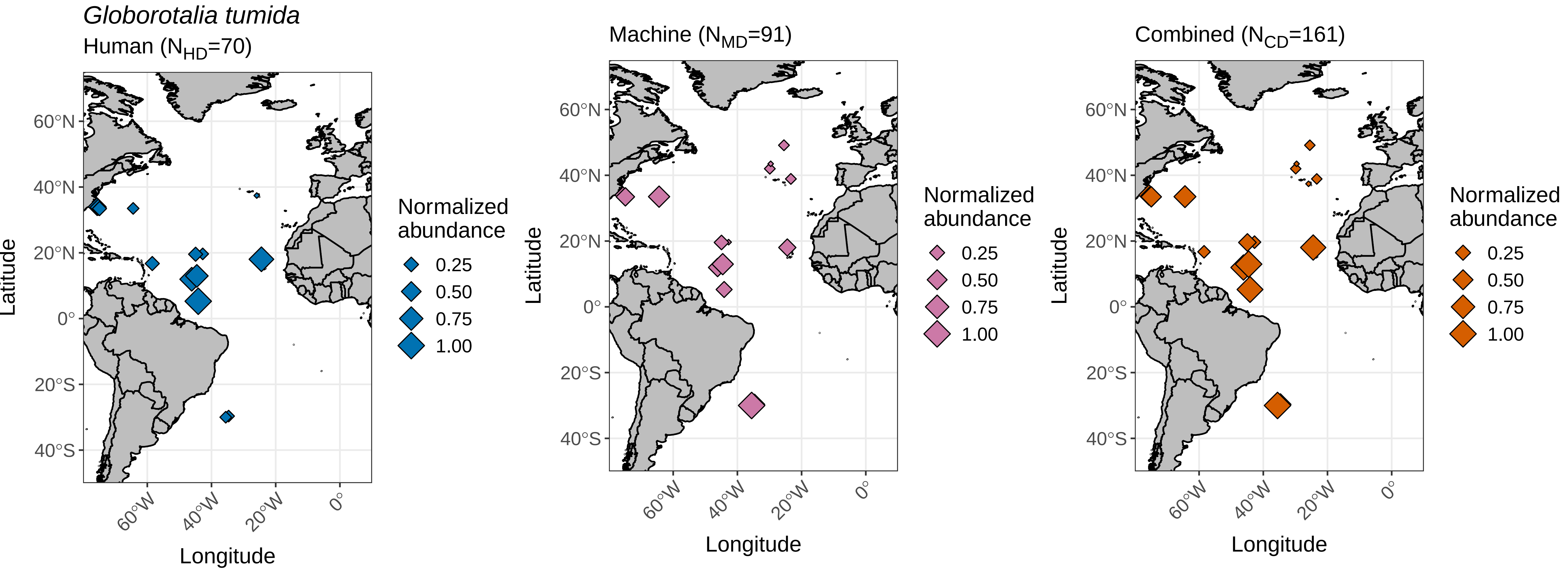

### globorotalia_ungulata_global_range_plot.pdf

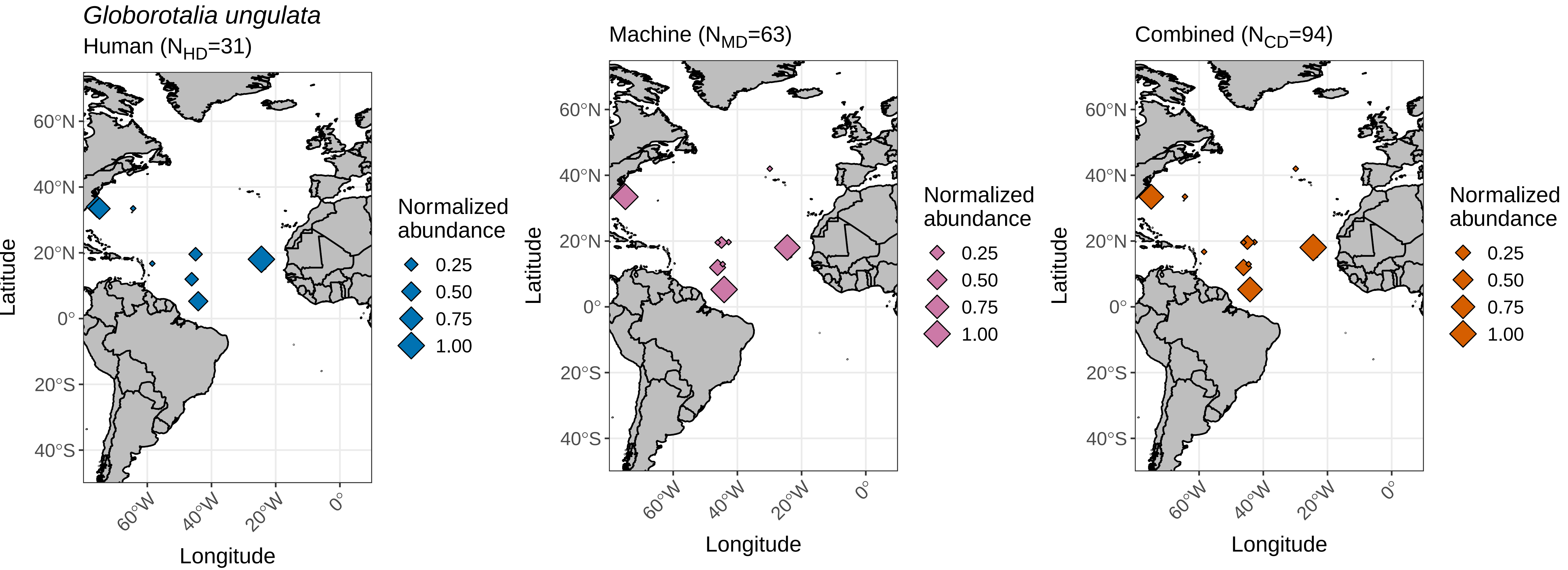

### globorotaloides_hexagonus_global_range_plot.pdf

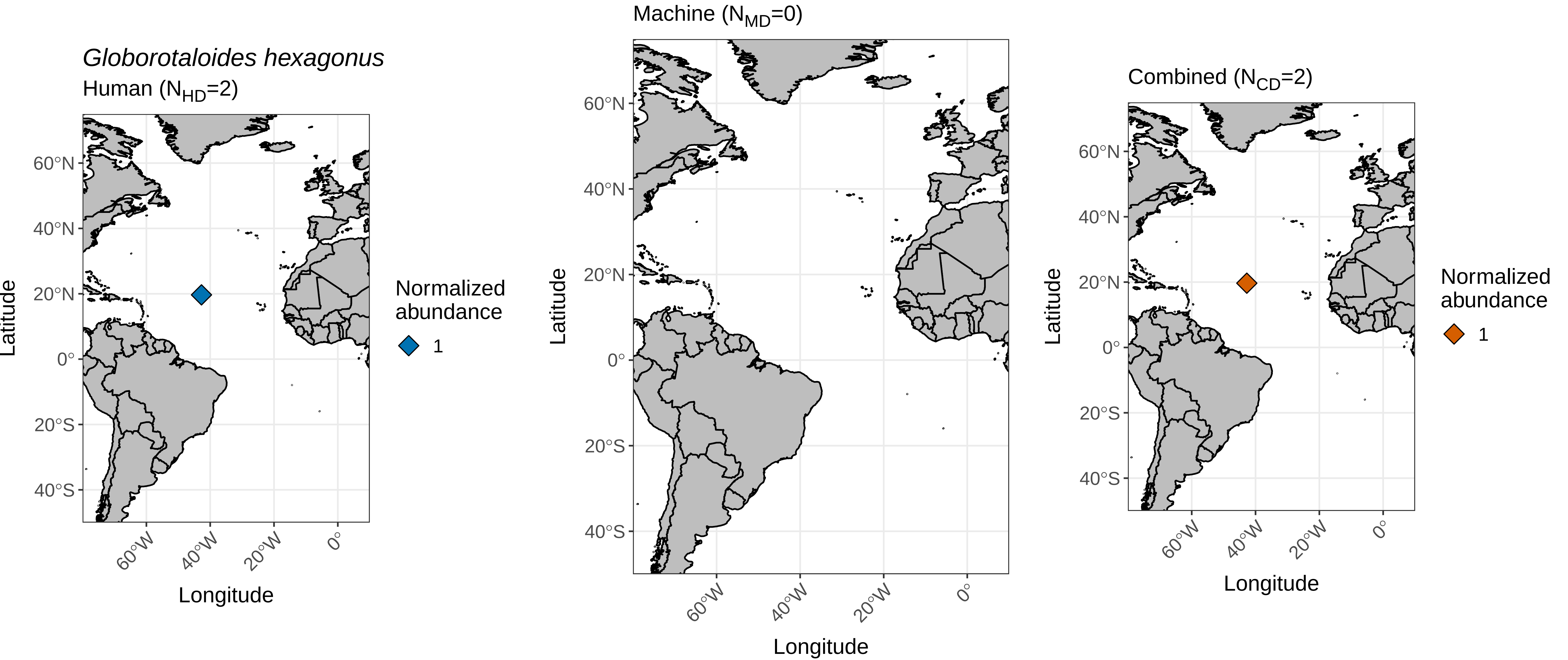

### globoturborotalita_rubescens_global_range_plot.pdf

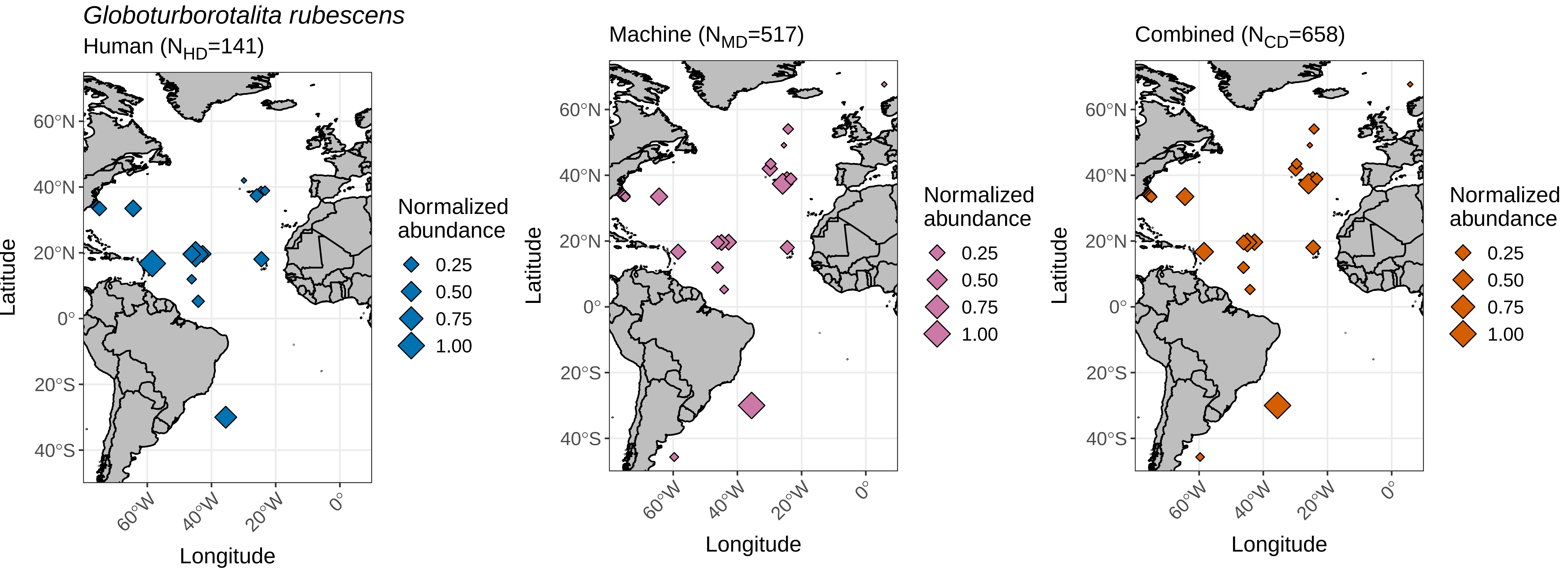

### globoturborotalita_tenella_global_range_plot.pdf

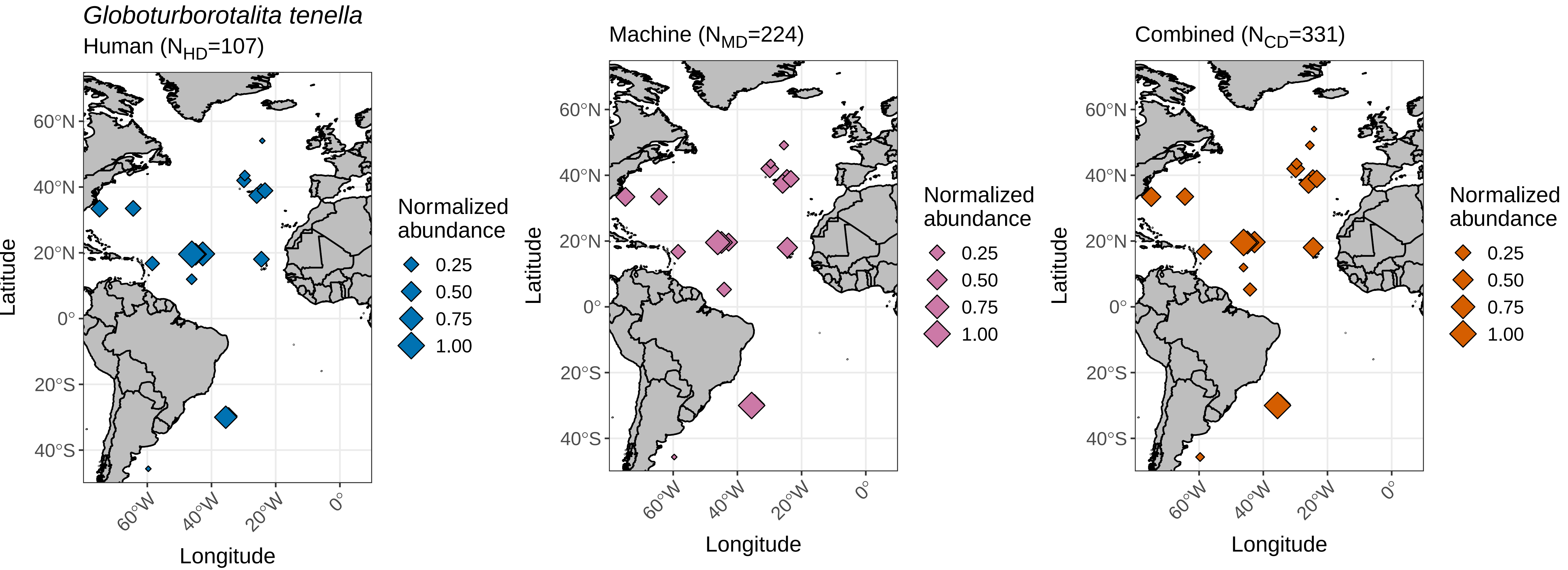

### hastigerina_pelagica_global_range_plot.pdf

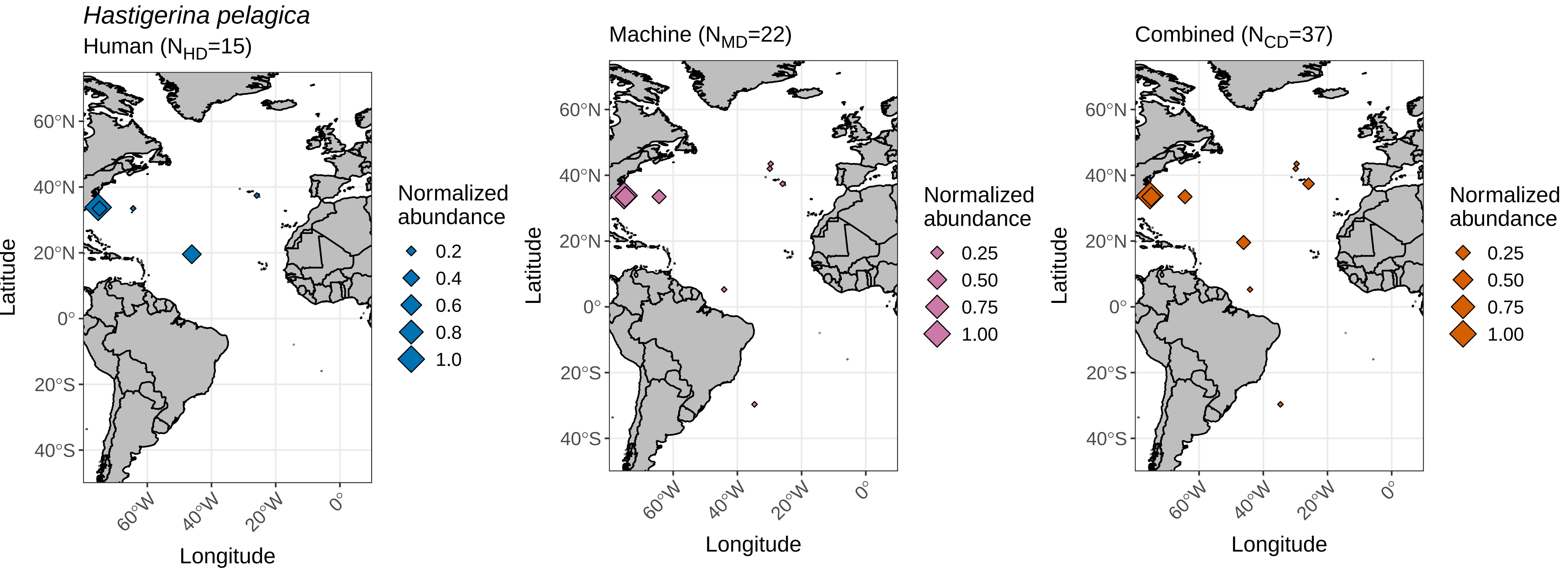

### IPE.08147.pdf

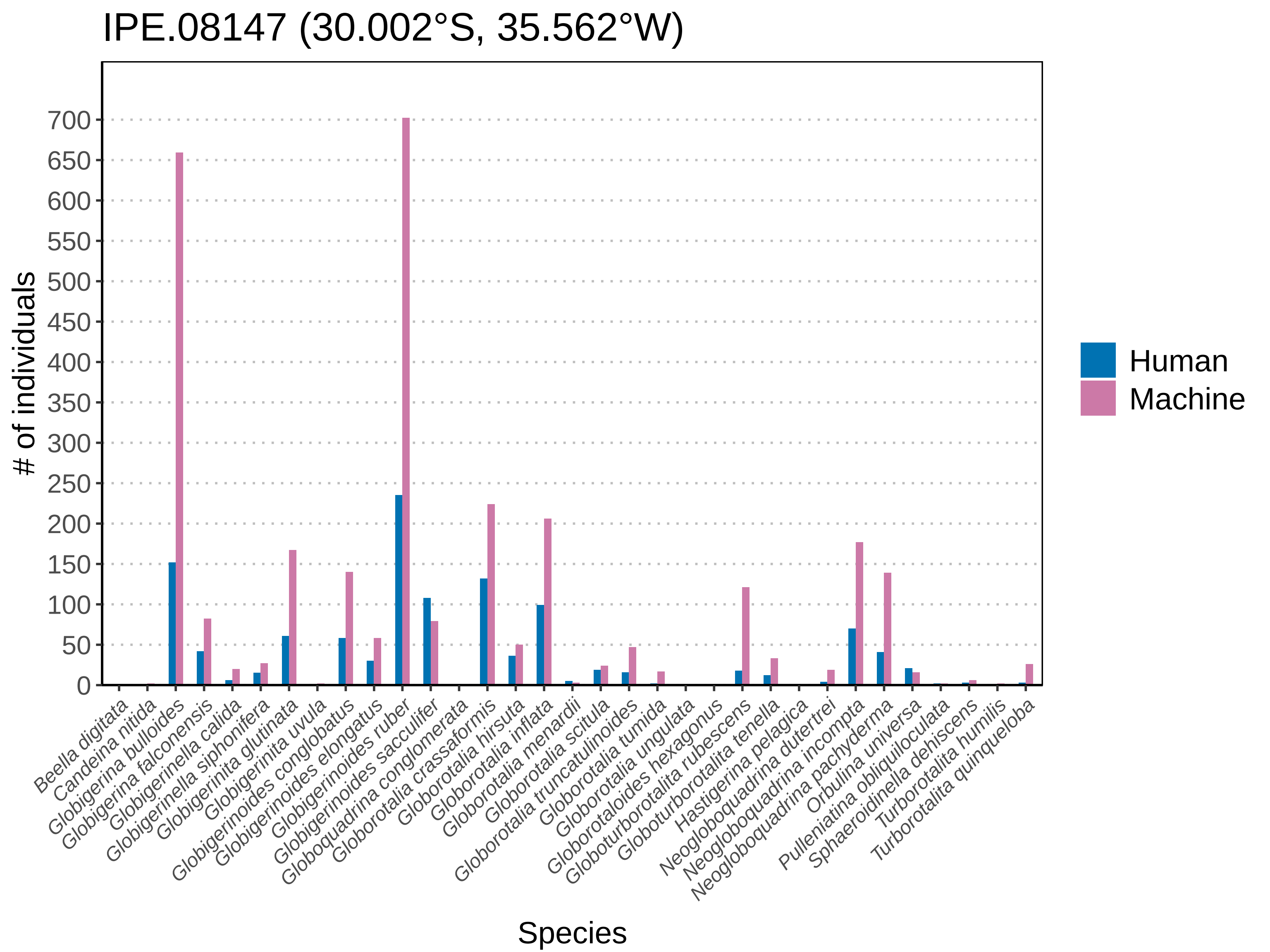

### IPE.08154.pdf

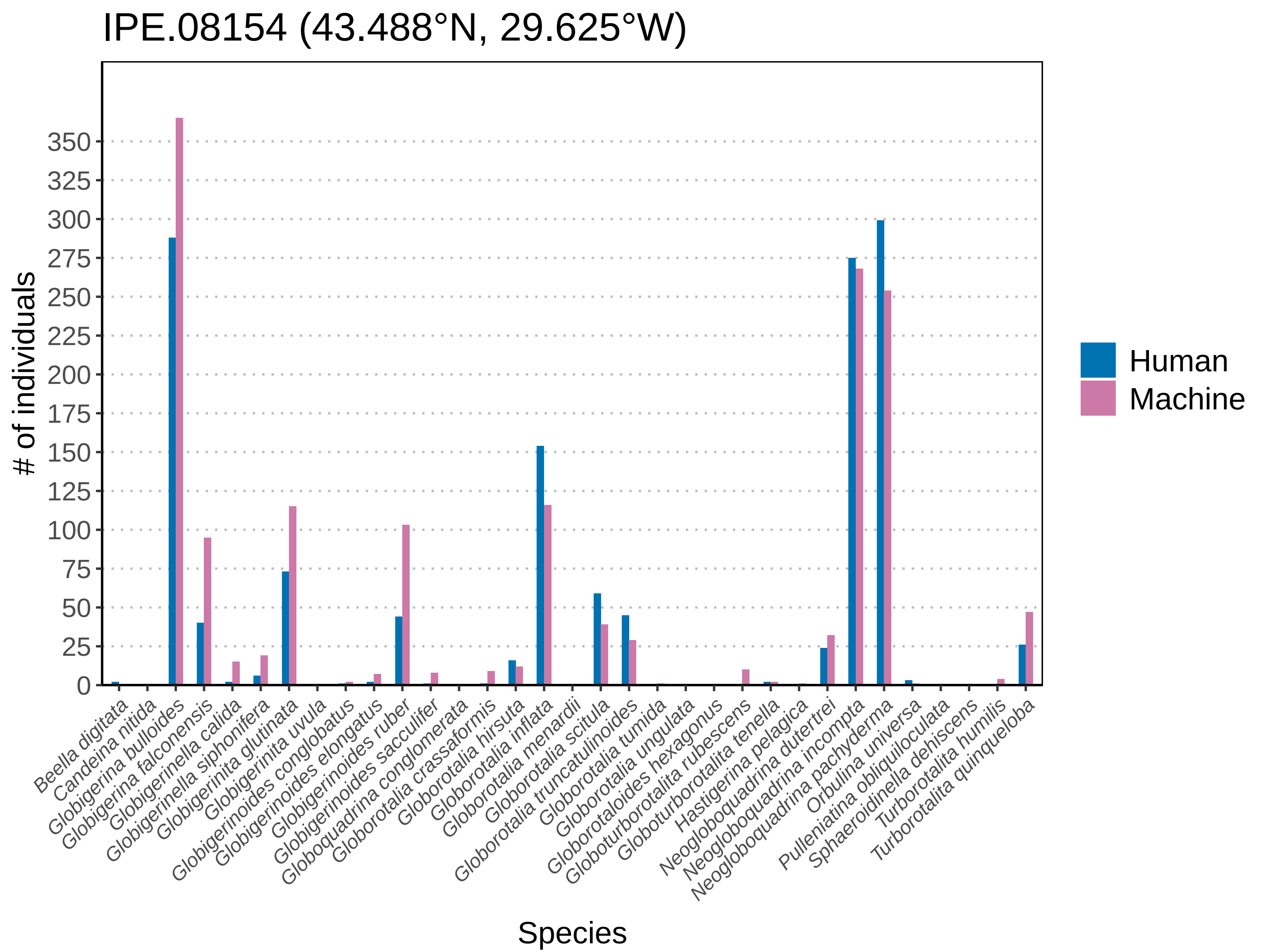

### IPE.08158.pdf

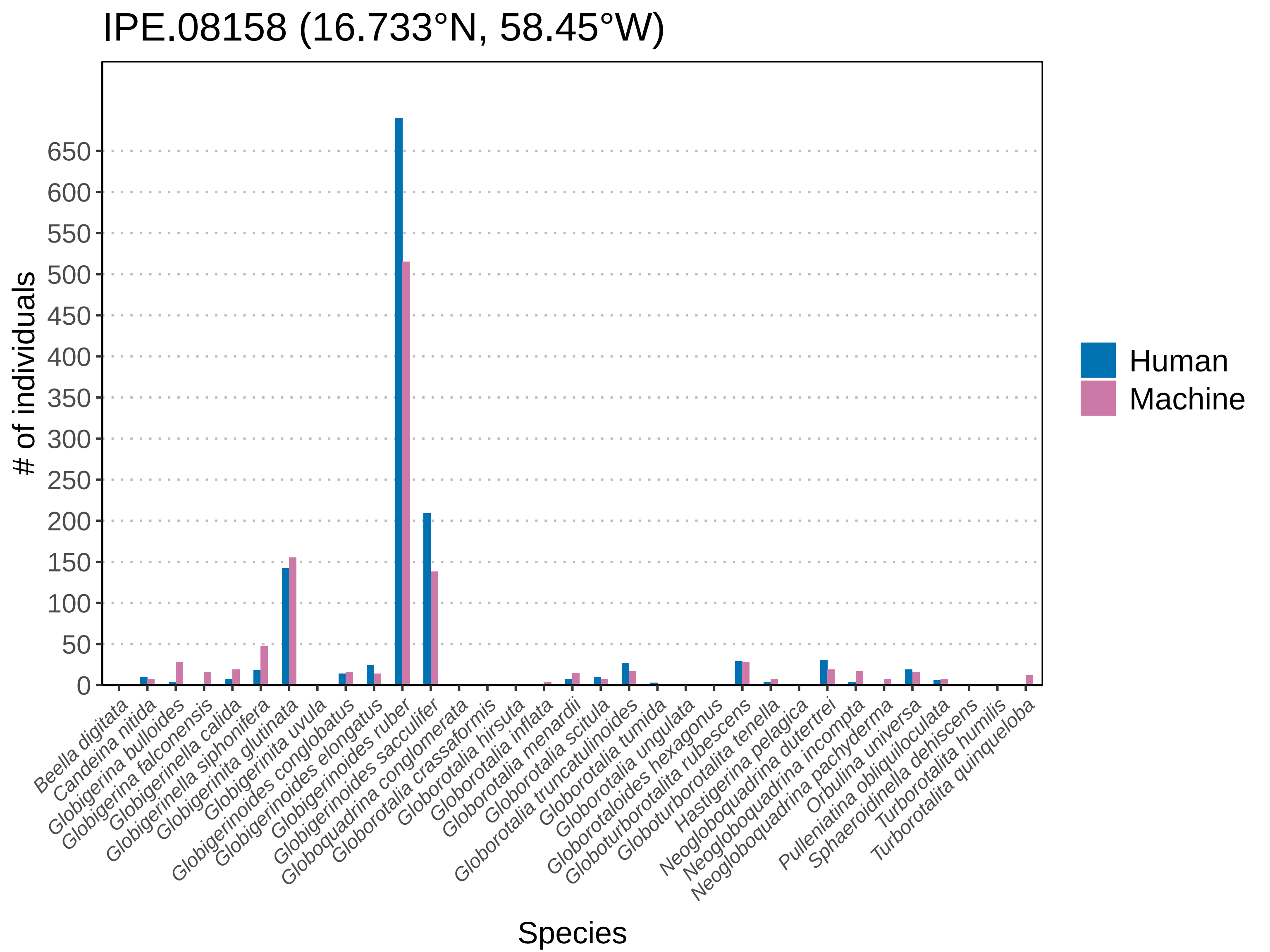

### IPE.08171.pdf

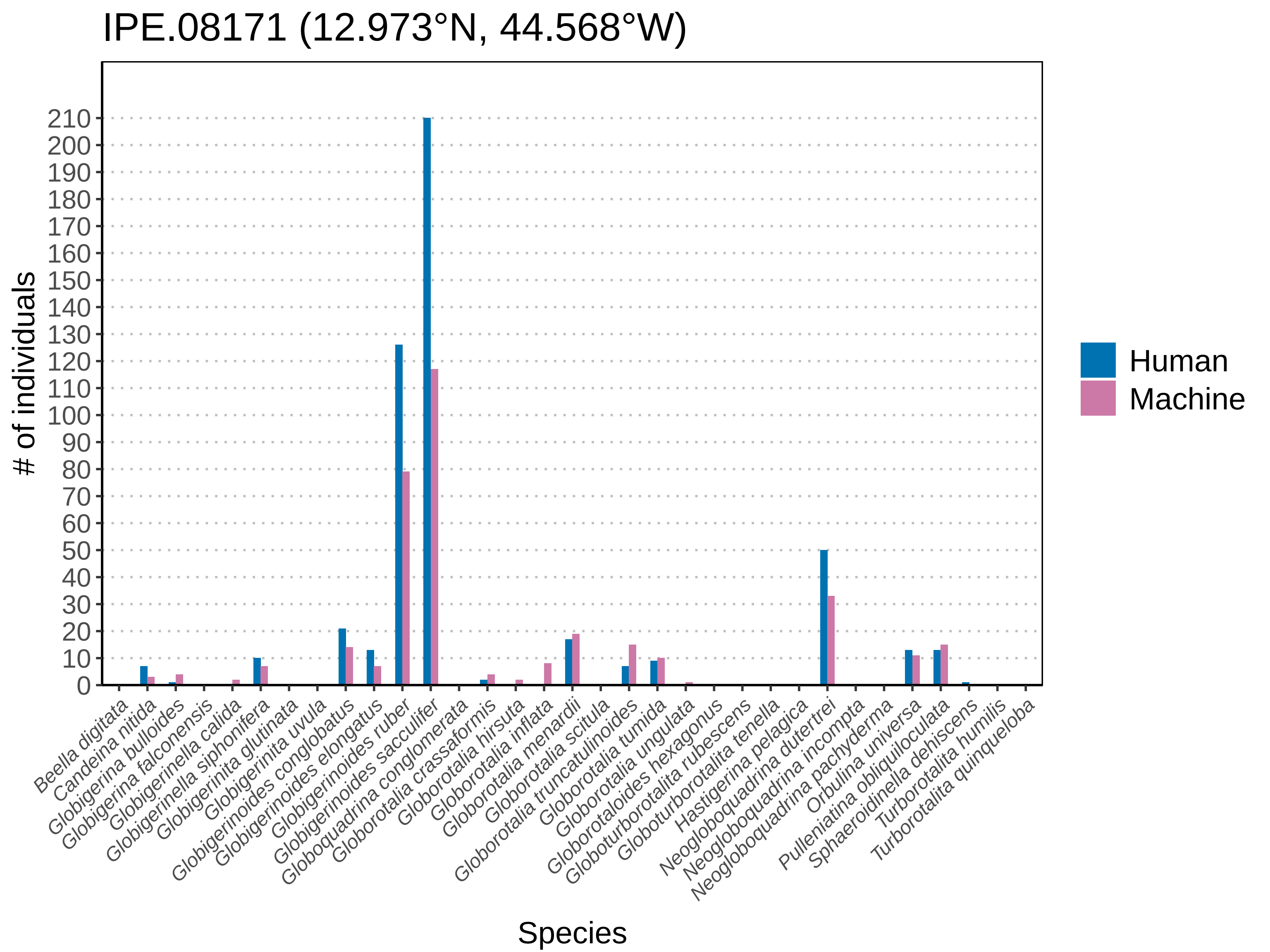

### IPE.08182.pdf

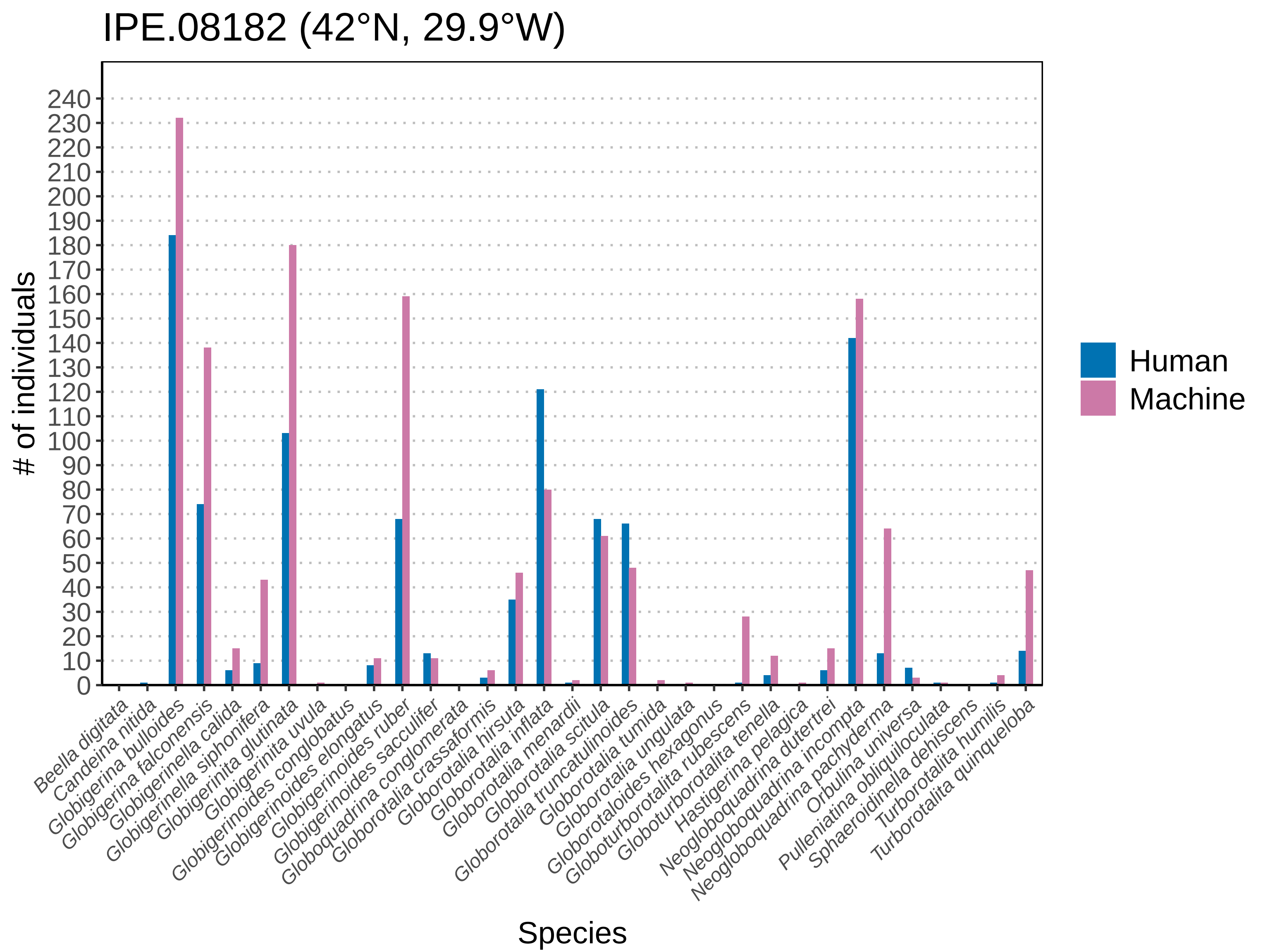

### IPE.08194.pdf

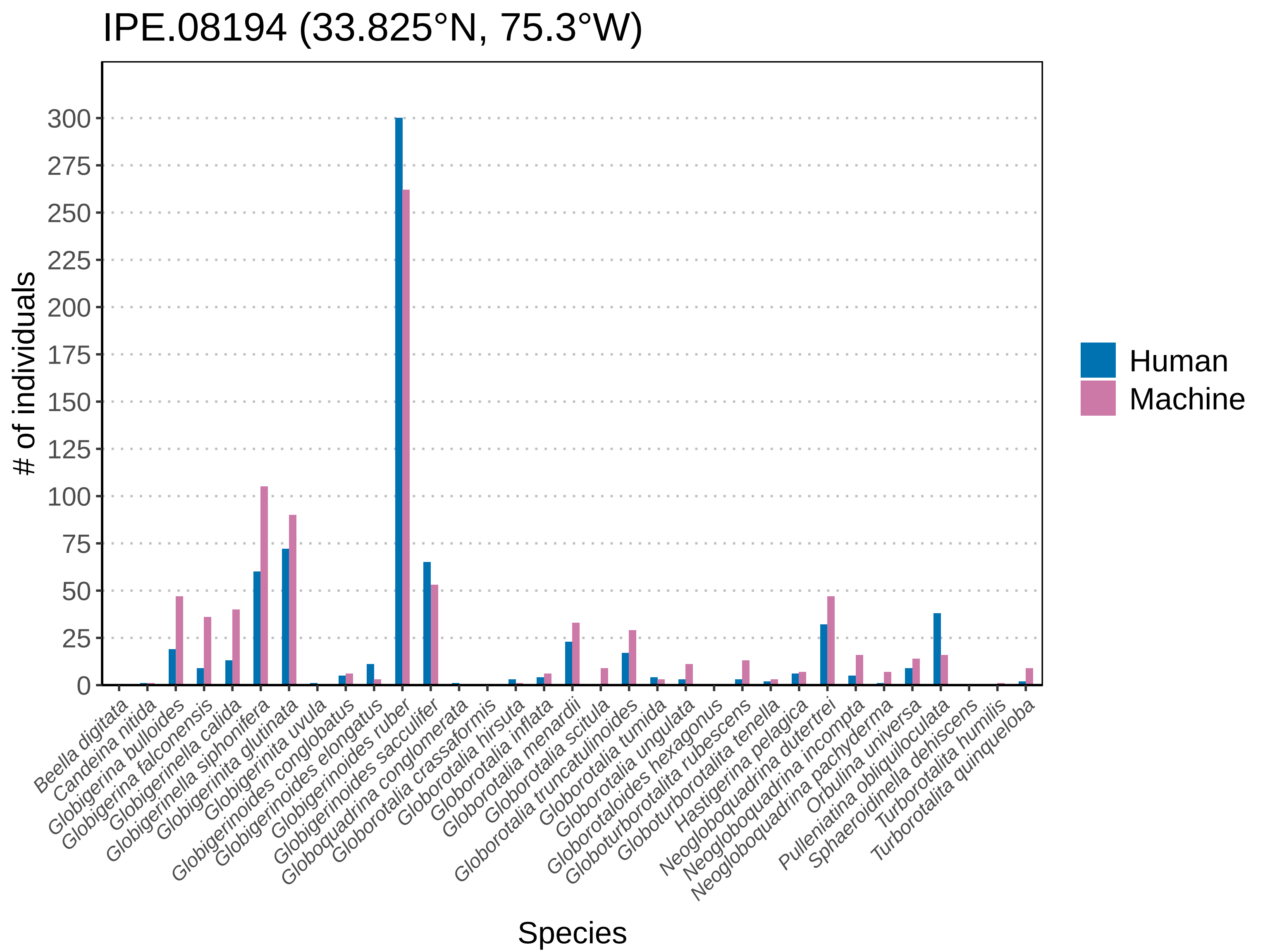

### IPE.08195.pdf

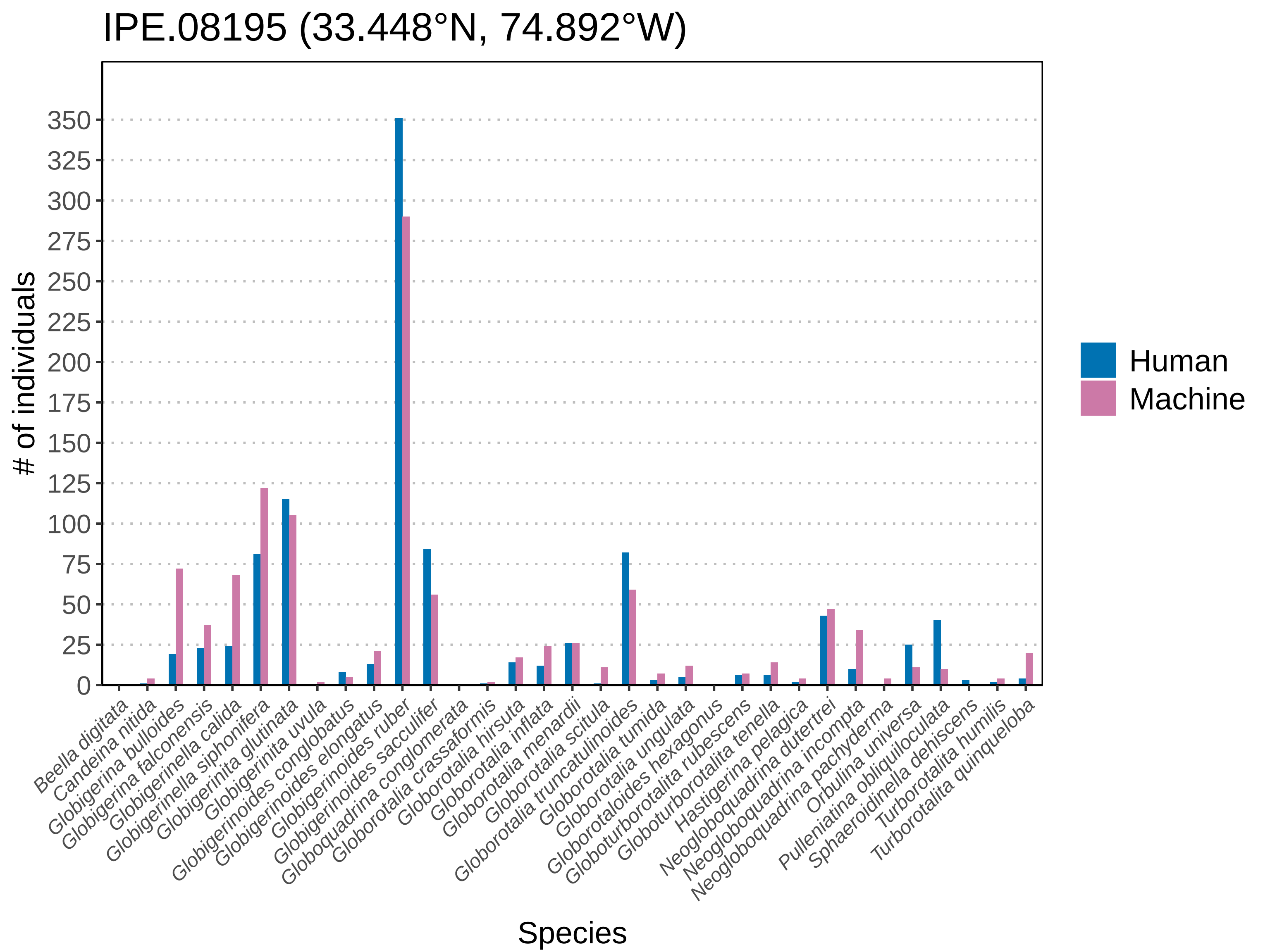

### IPE.08196.pdf

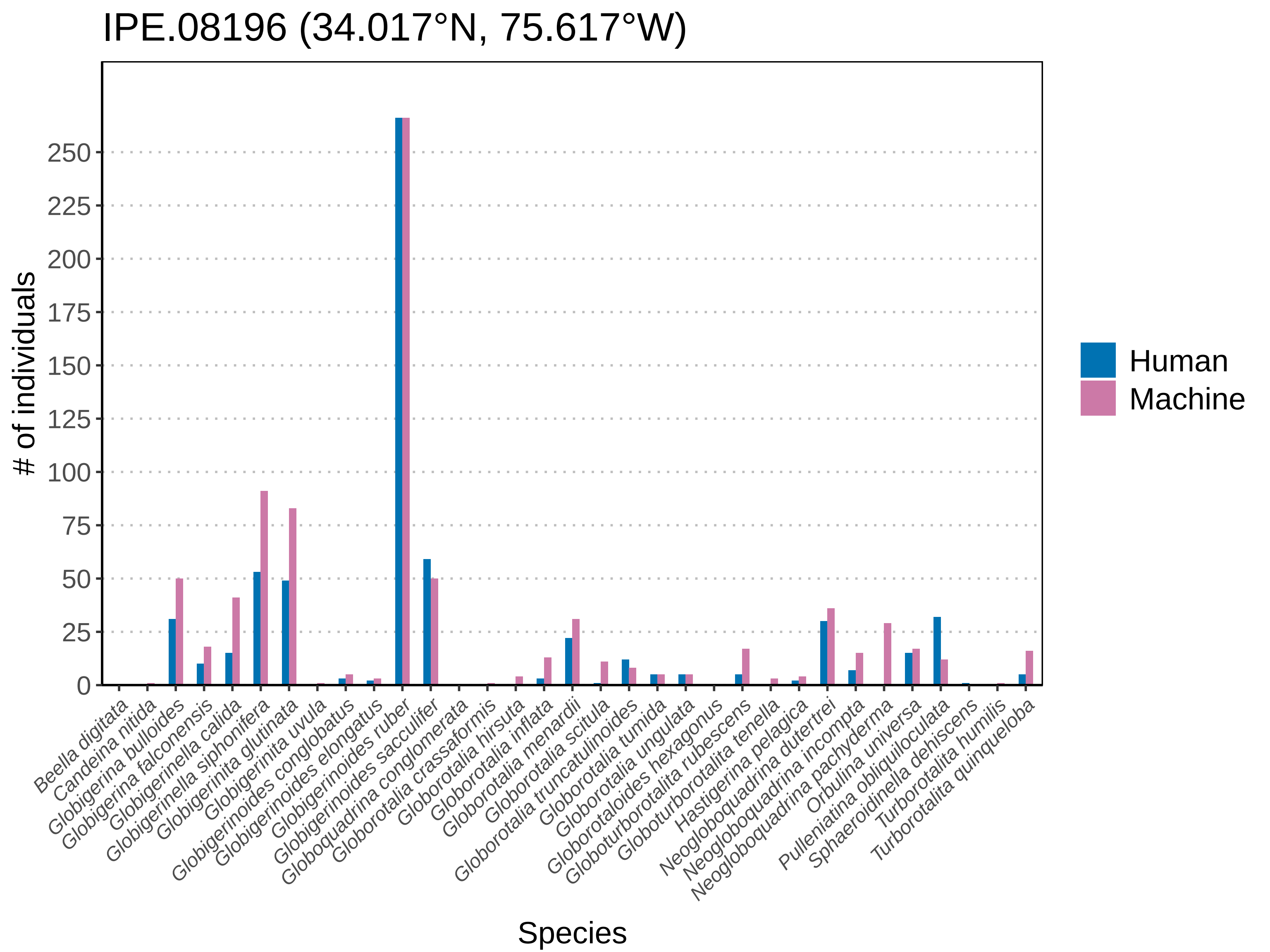

### IPE.08199.pdf

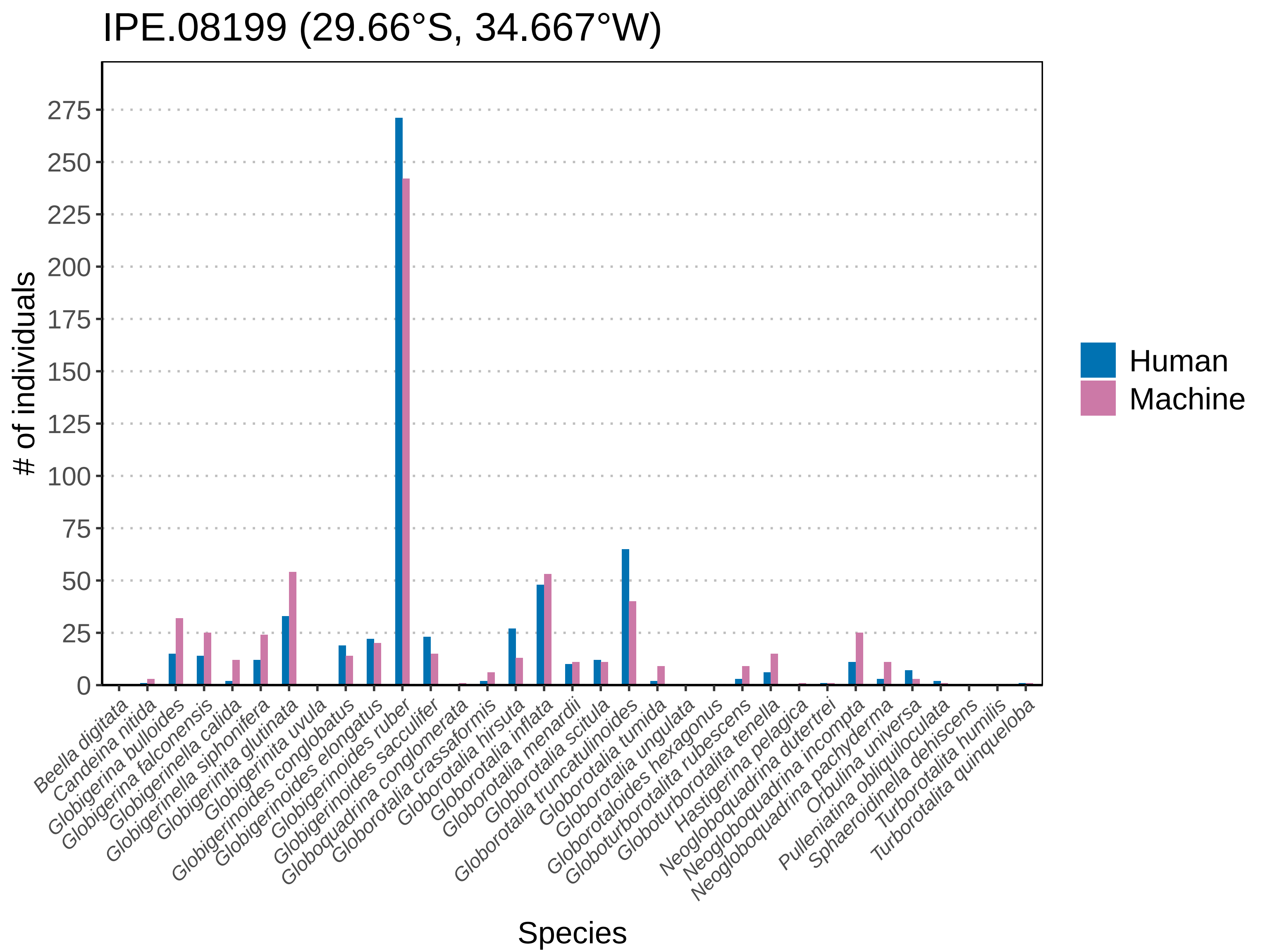

### IPE.08204.pdf

IPE.08204 (11.958°N, 46.167°W)

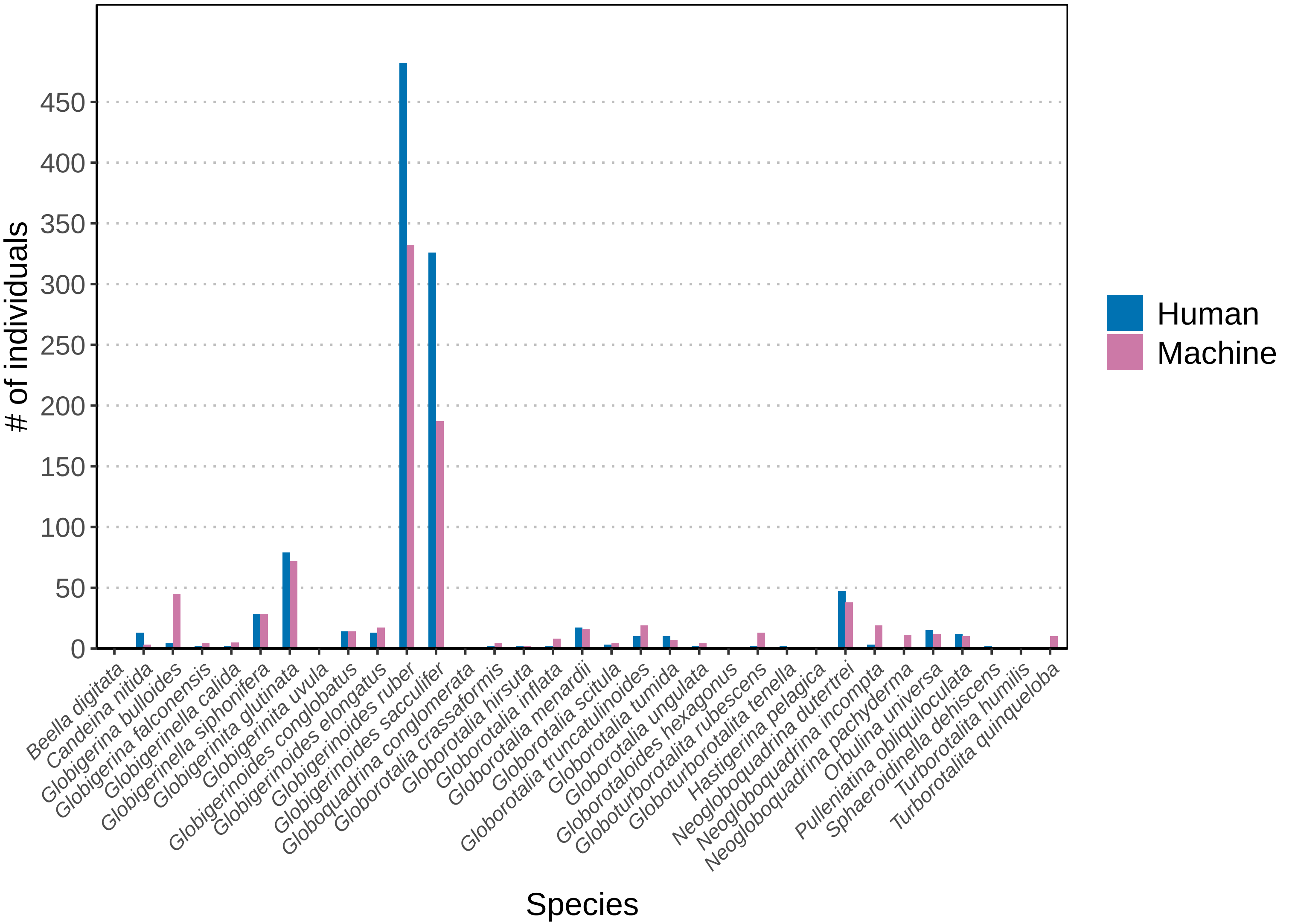

### IPE.08208.pdf

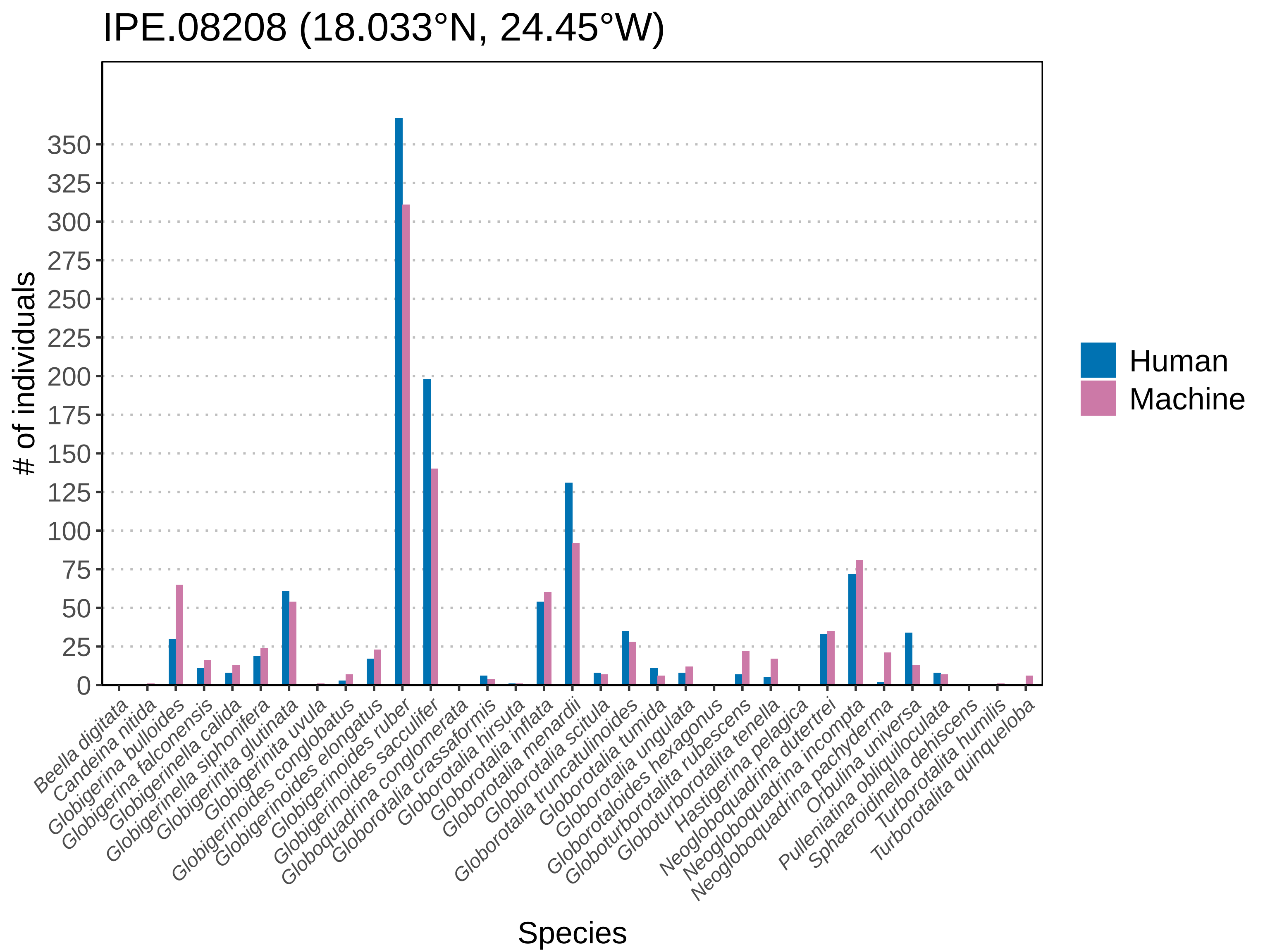

### IPE.08209.pdf

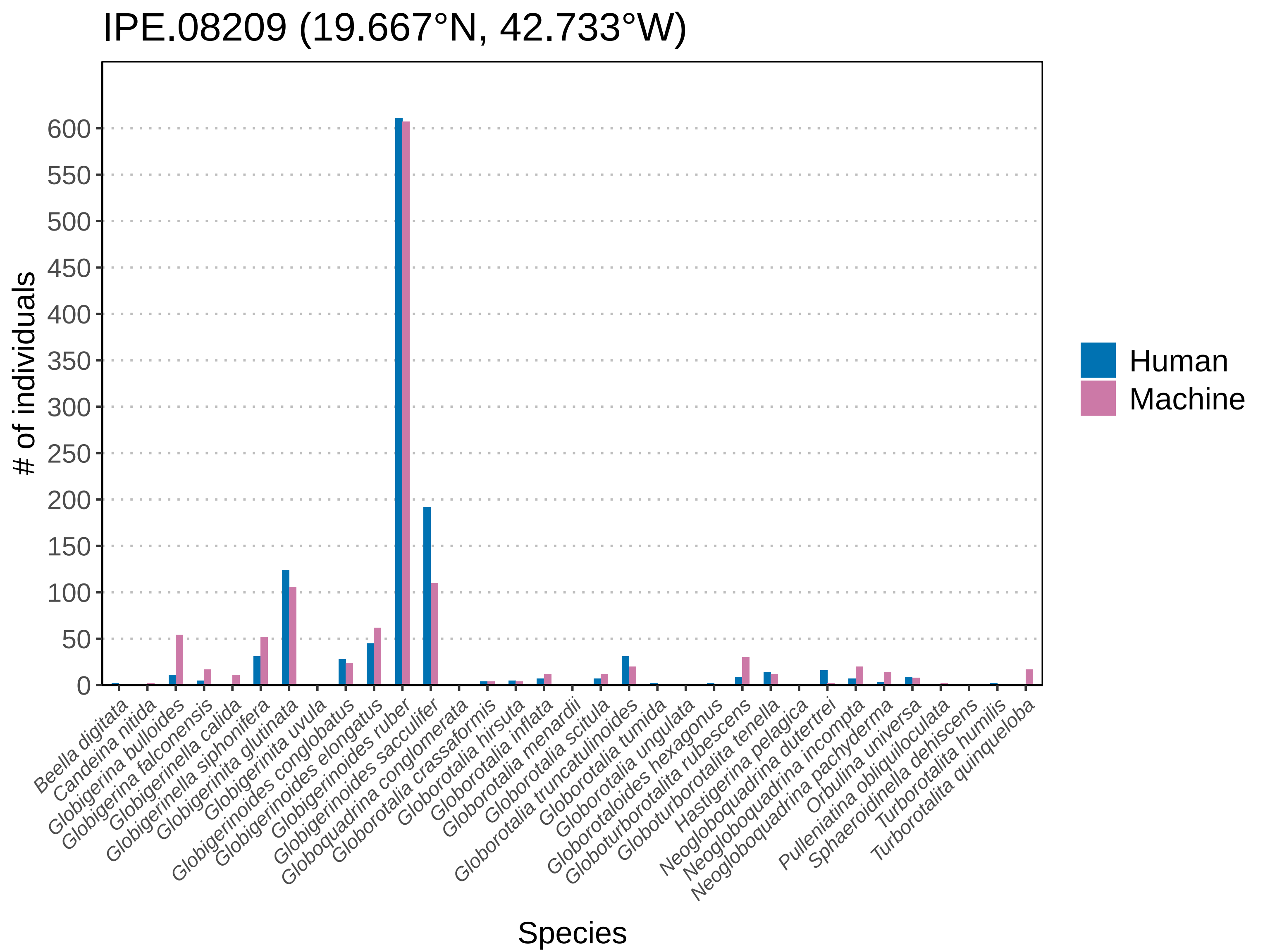

### IPE.08210.pdf

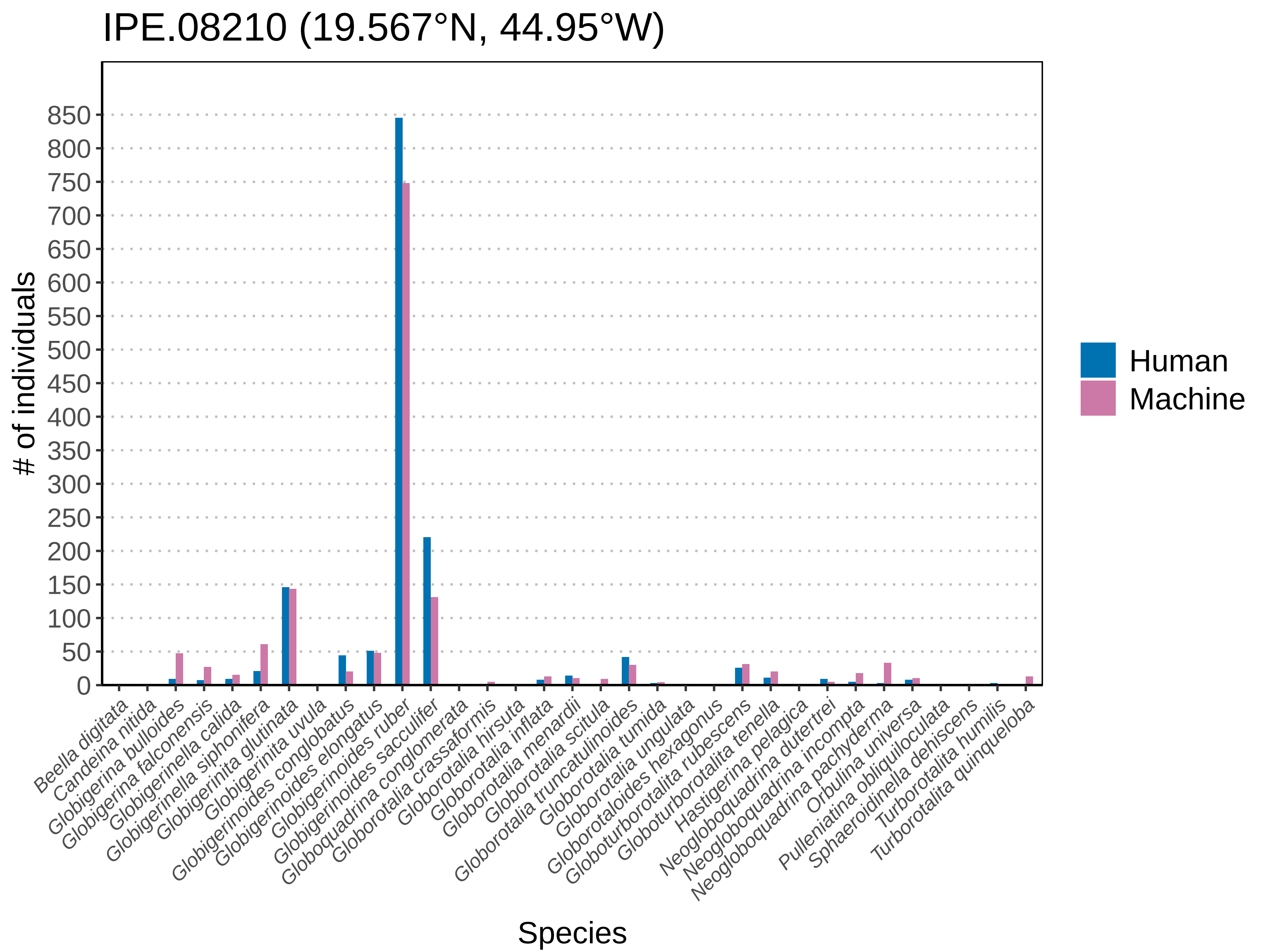

### IPE.08215.pdf

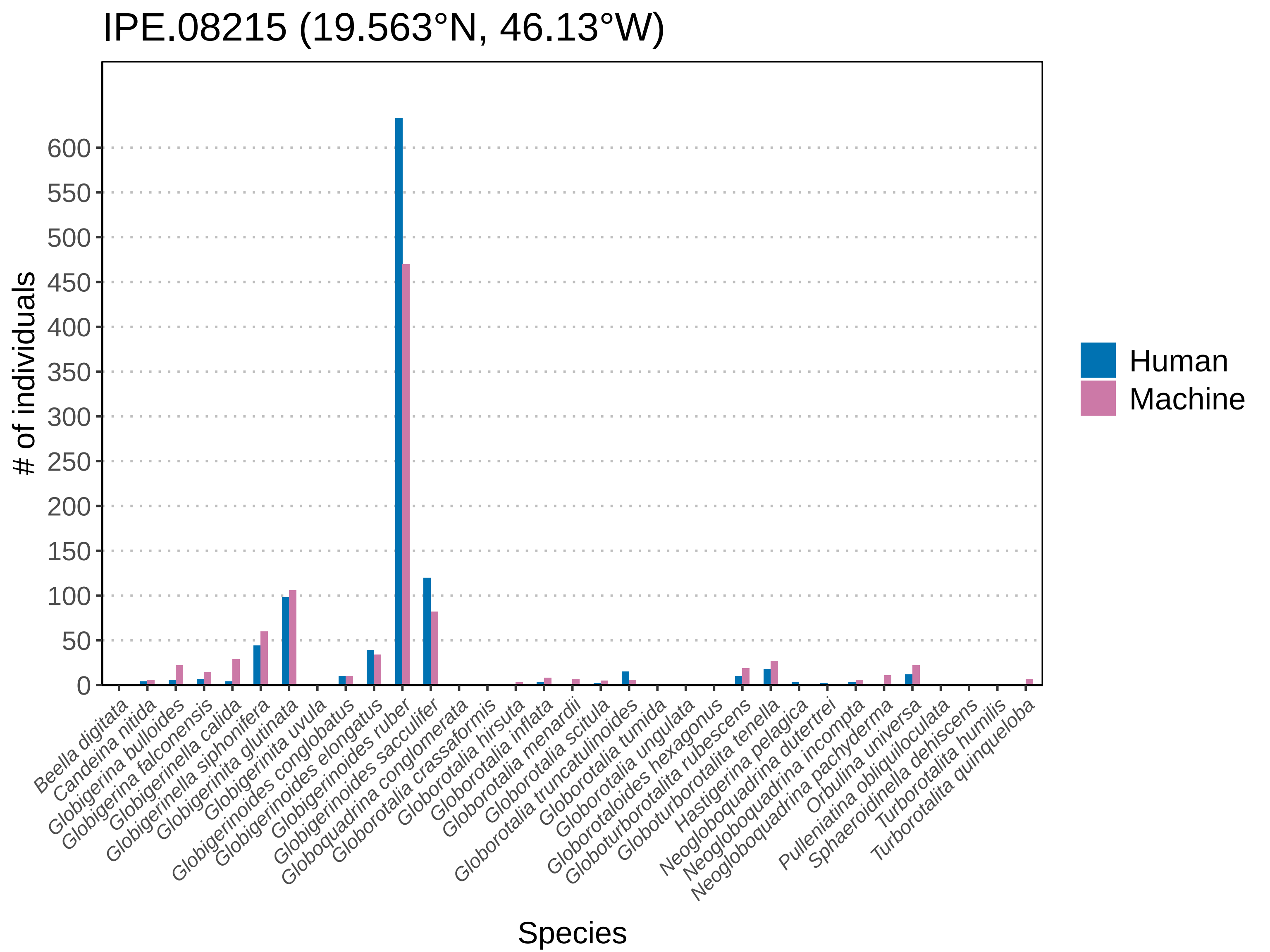

### IPE.08248.pdf

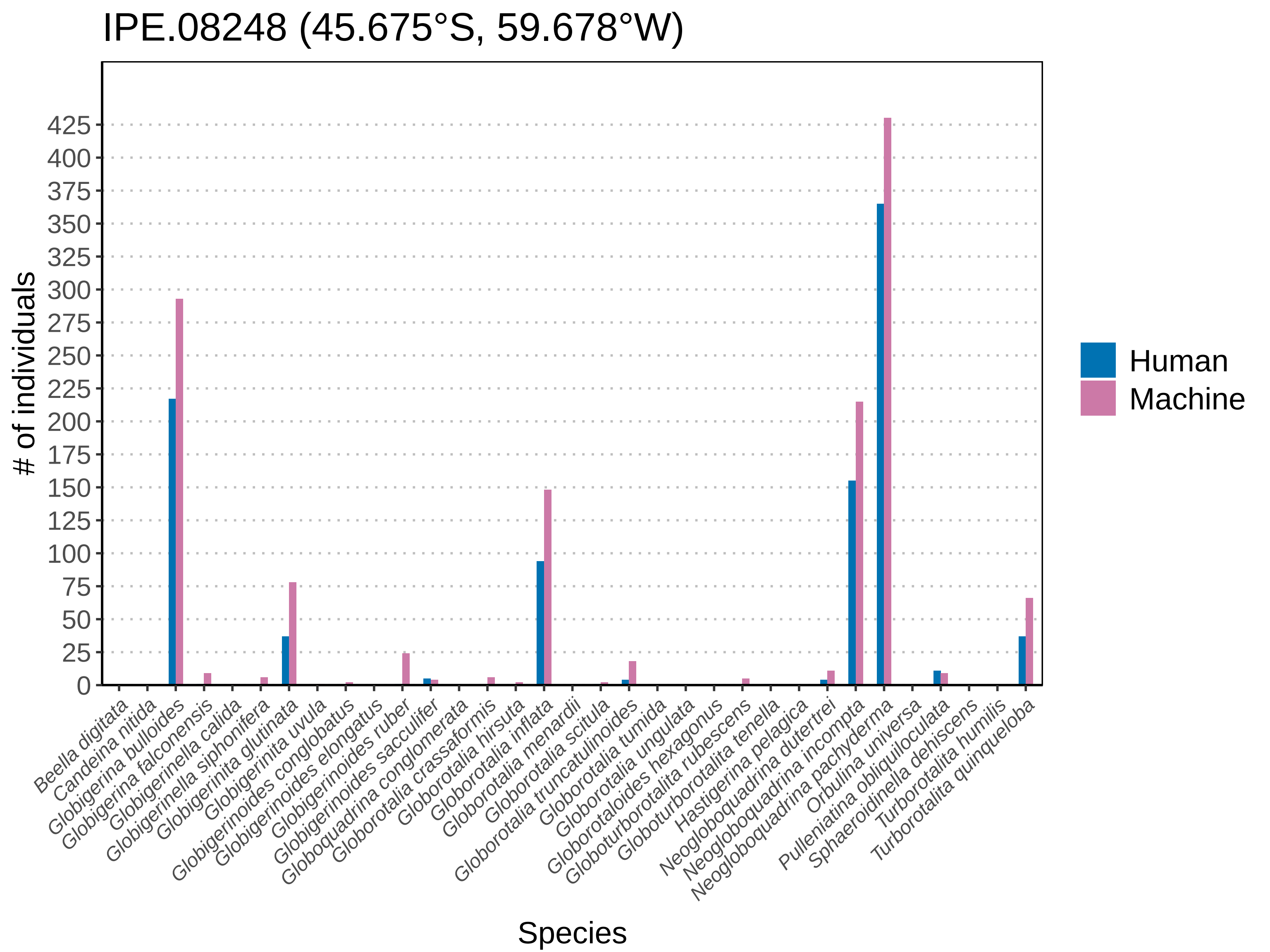

### IPE.08259.pdf

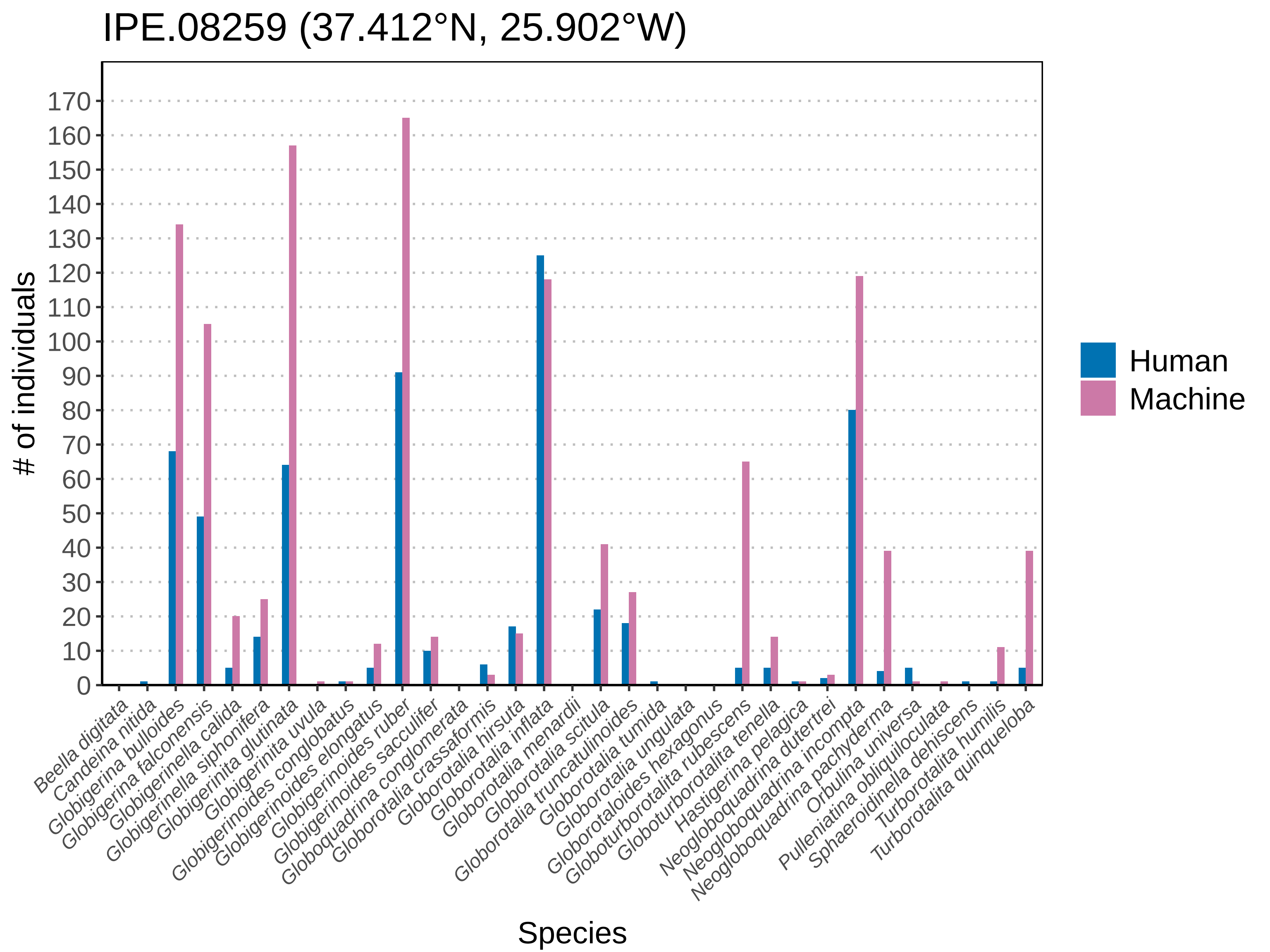

### IPE.08262.pdf

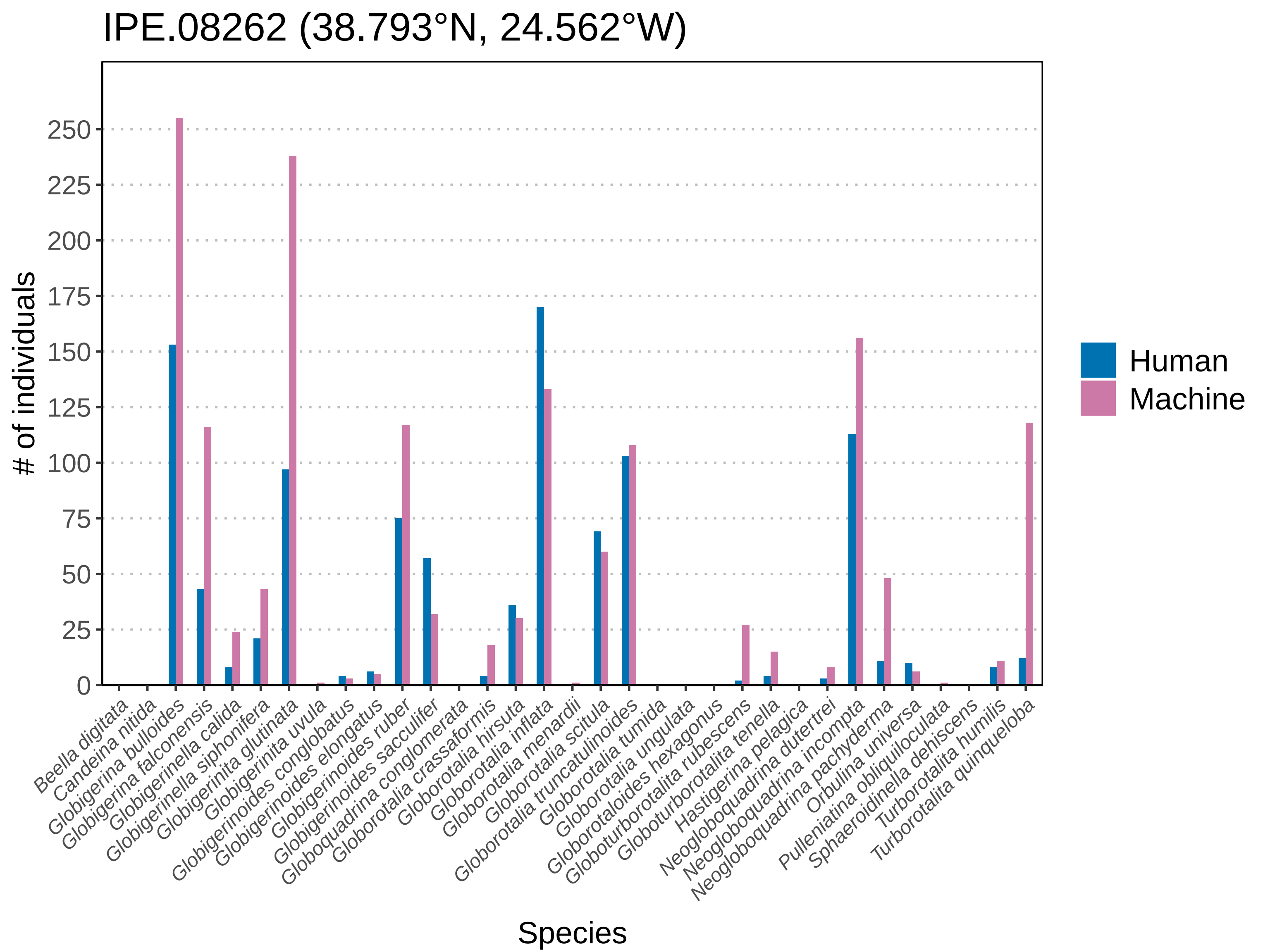

### IPE.08264.pdf

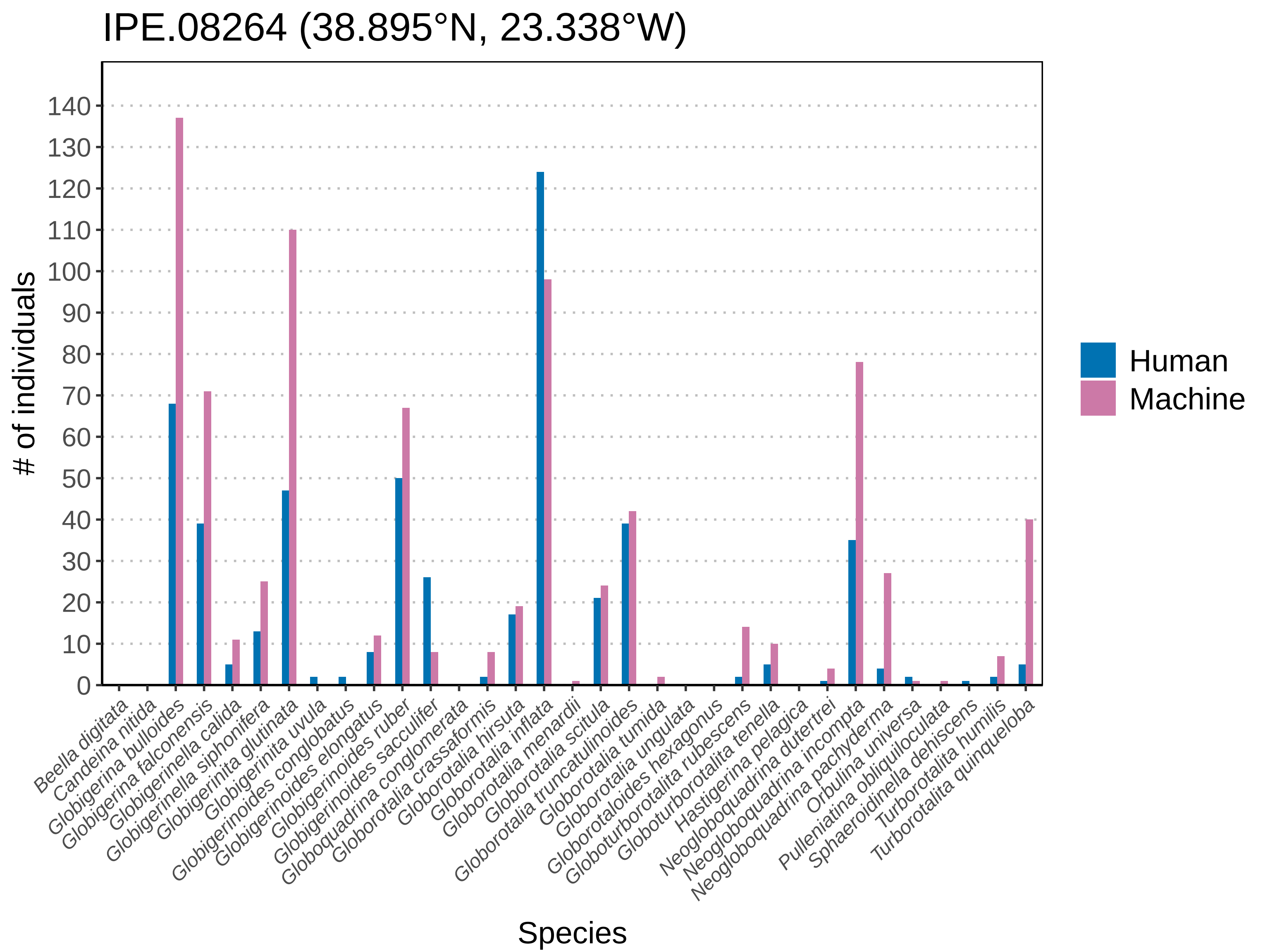

### IPE.08295.pdf

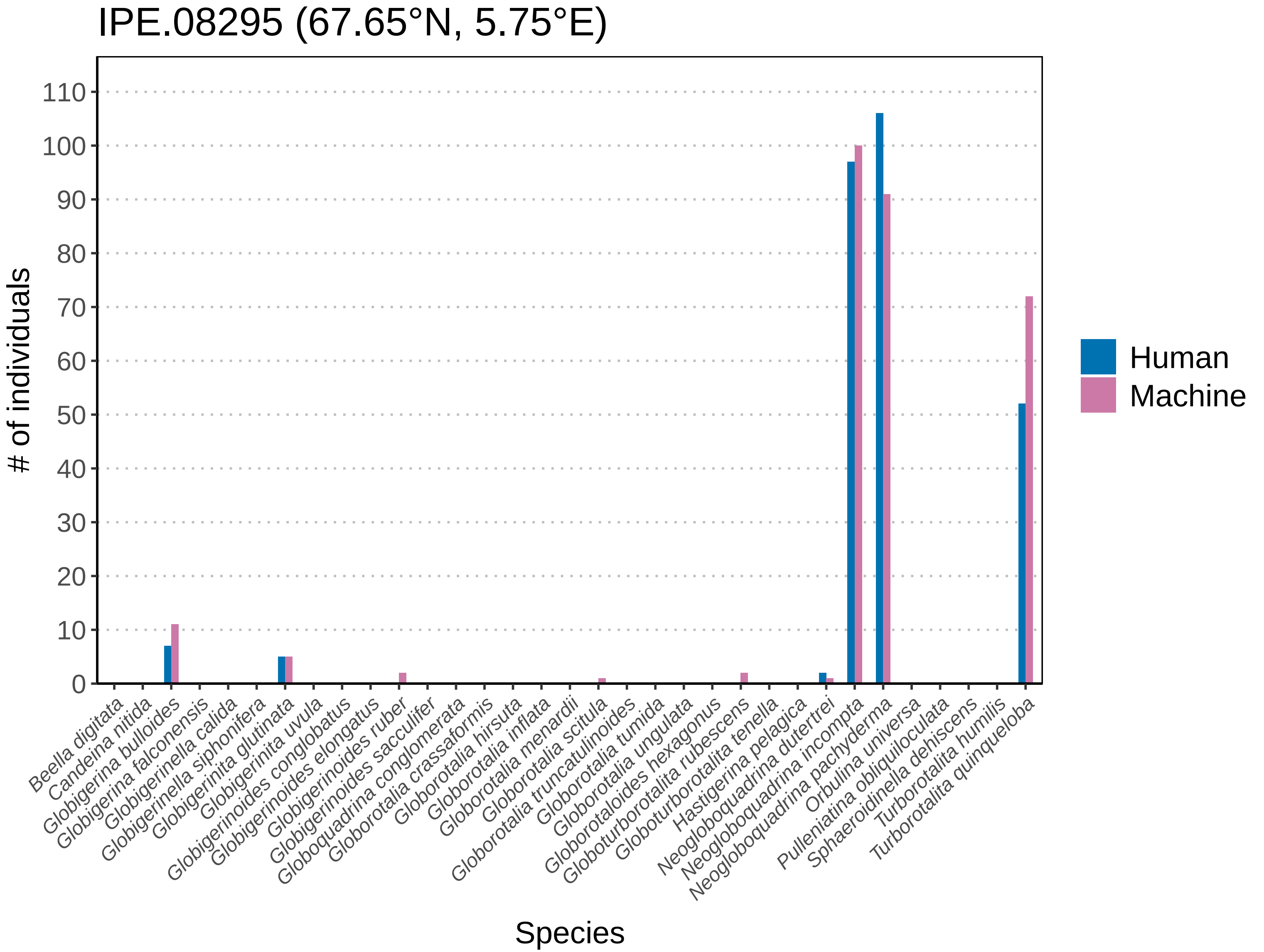

### IPE.08316.pdf

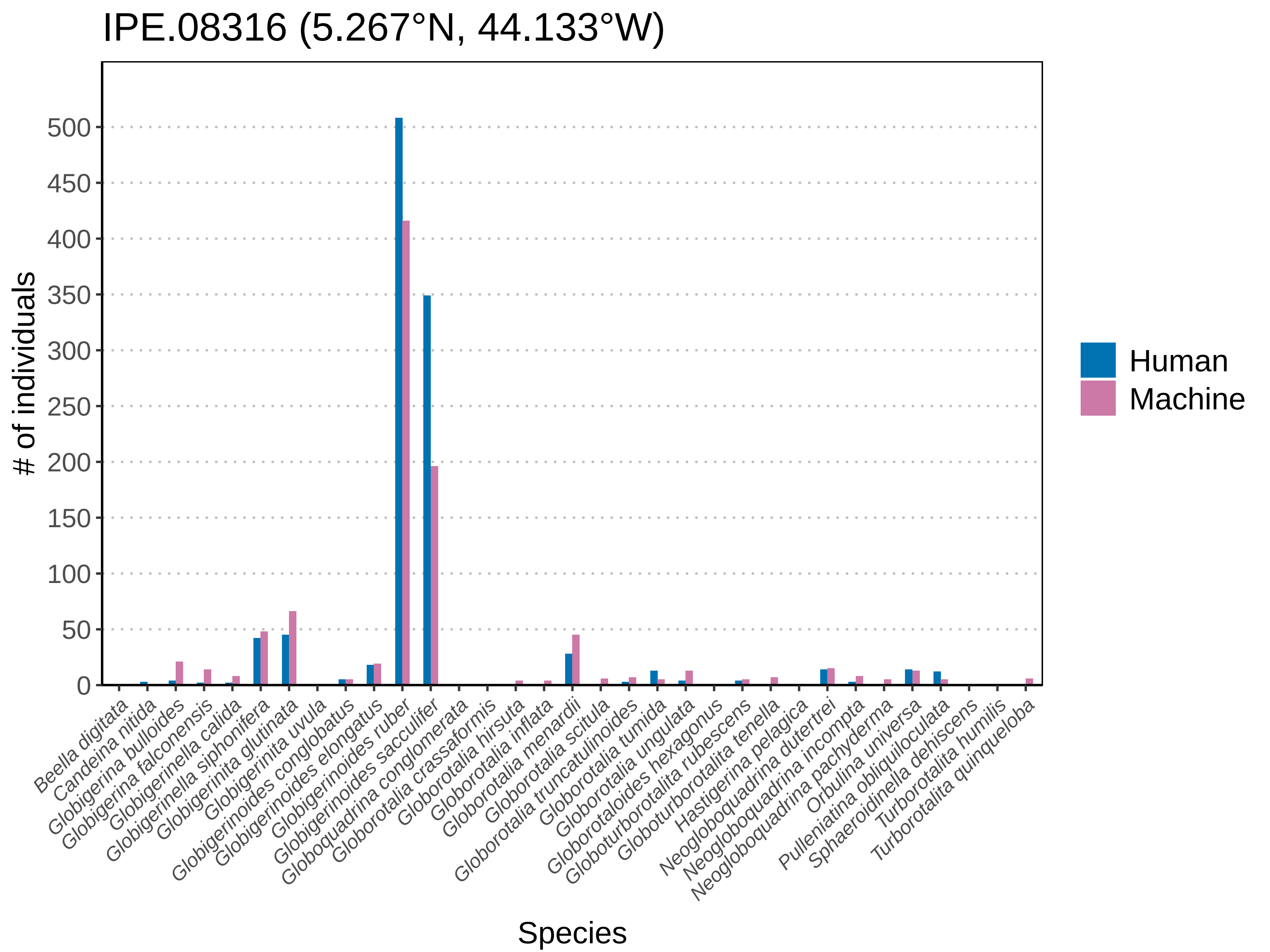

### IPE.08320.pdf

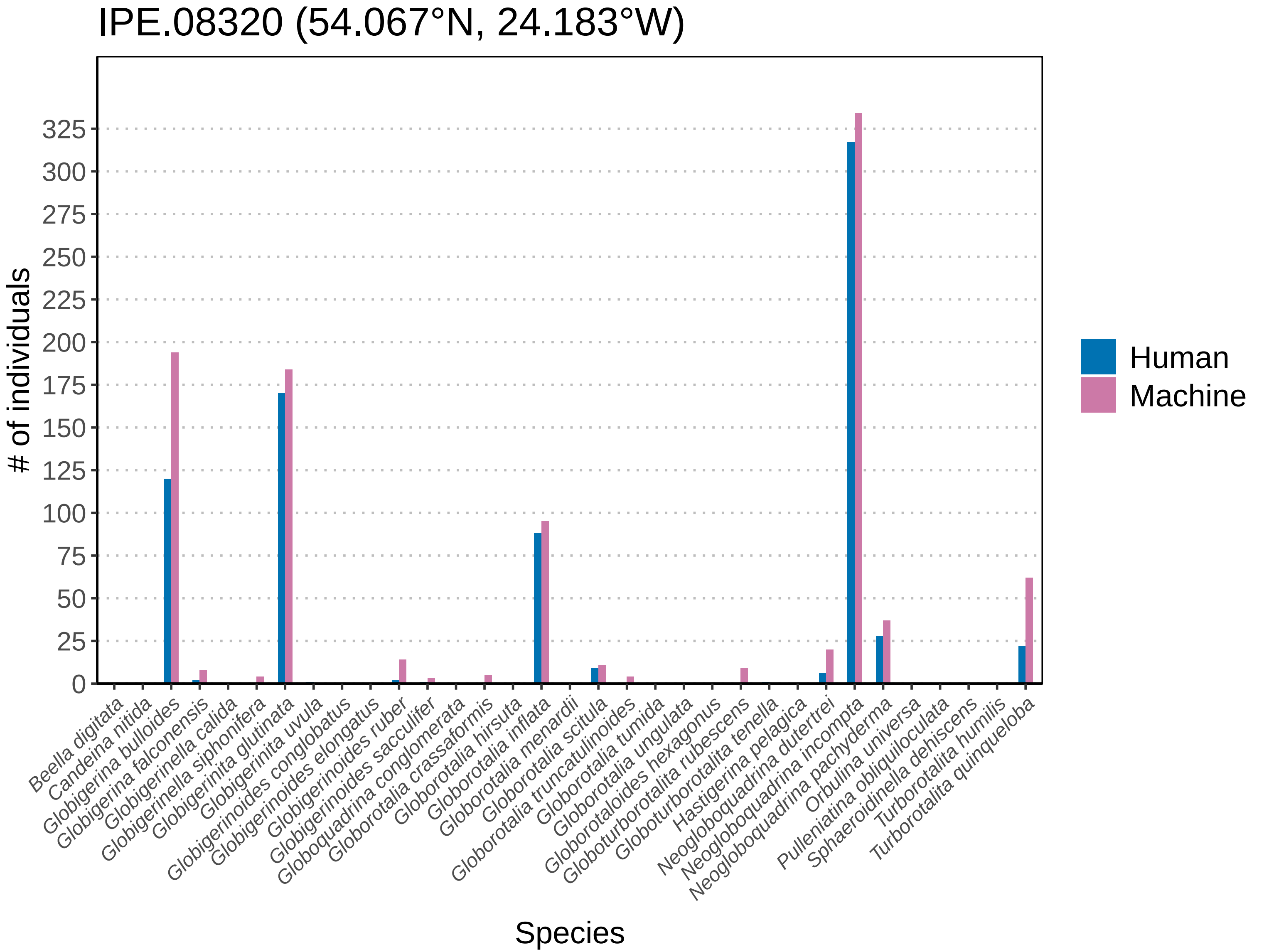

### IPE.08322.pdf

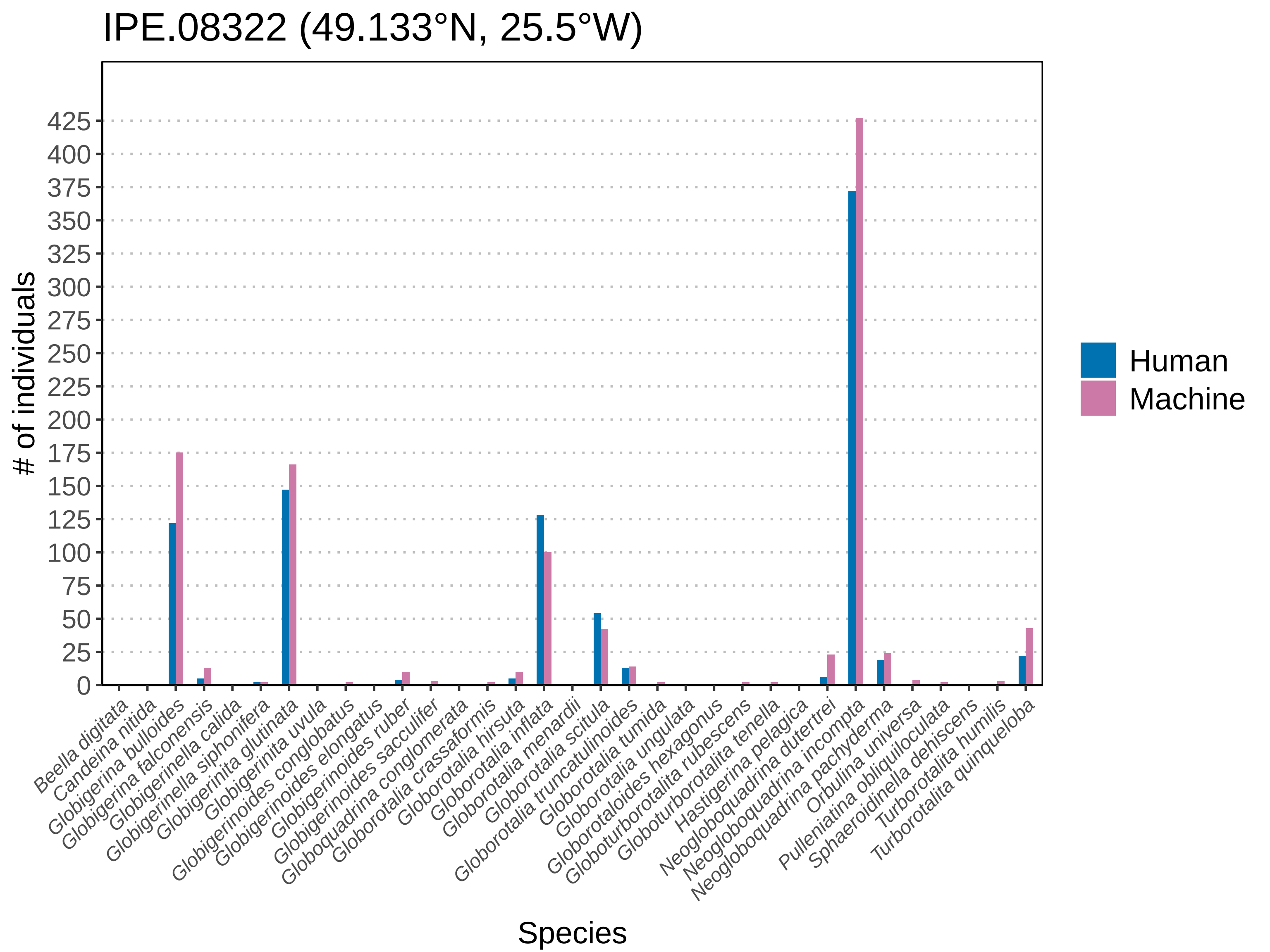

### IPE.08378.pdf

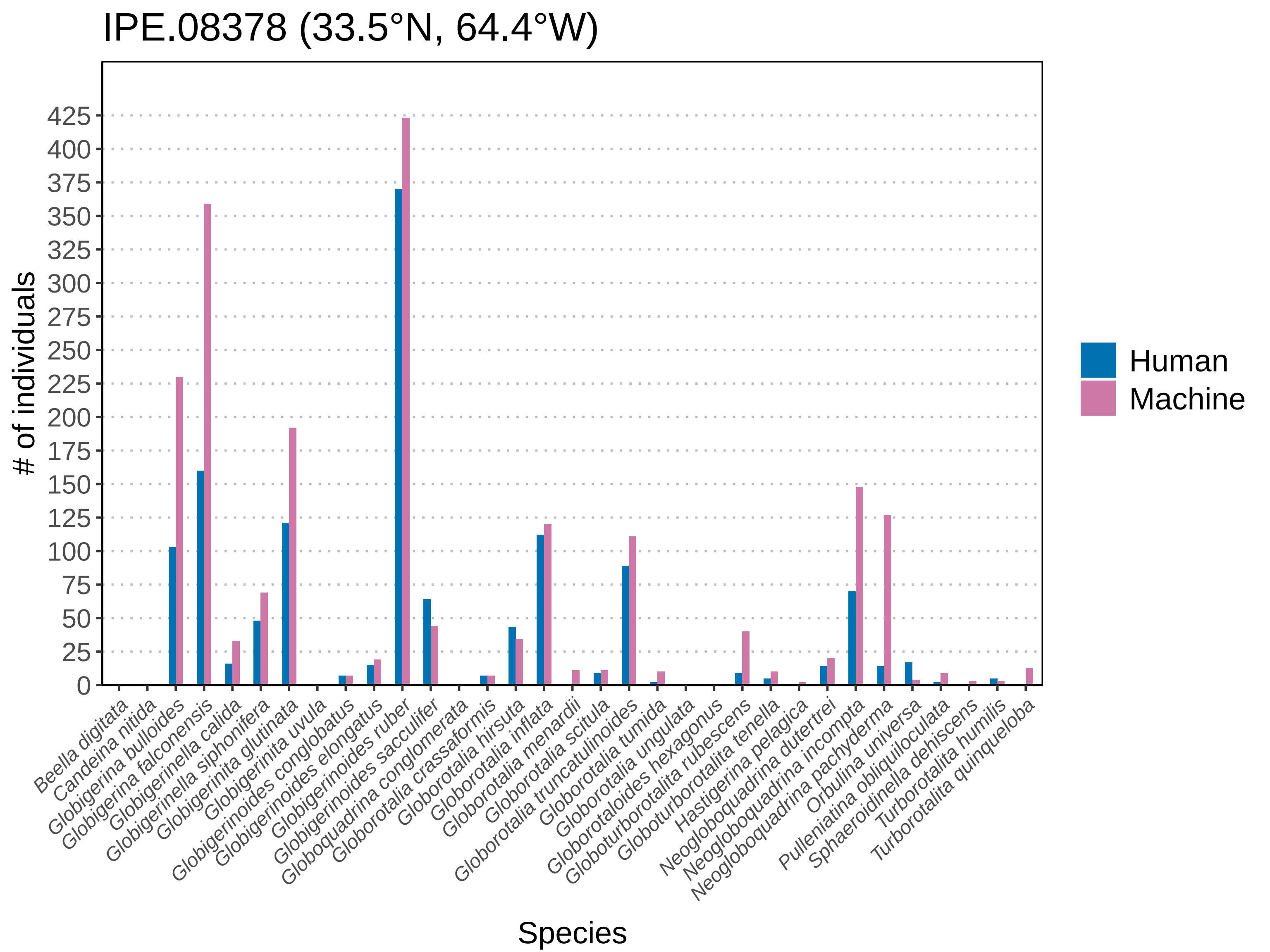

### neogloboquadrina_dutertrei_global_range_plot.pdf

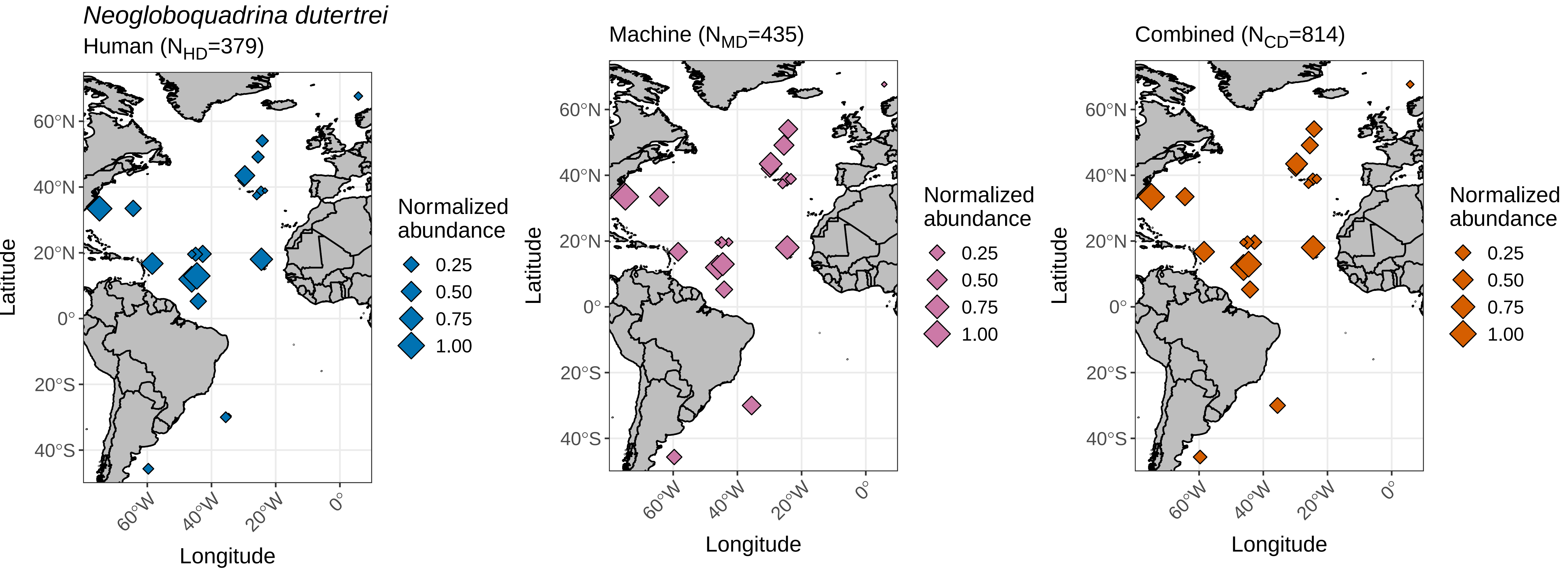

### neogloboquadrina_incompta_global_range_plot.pdf

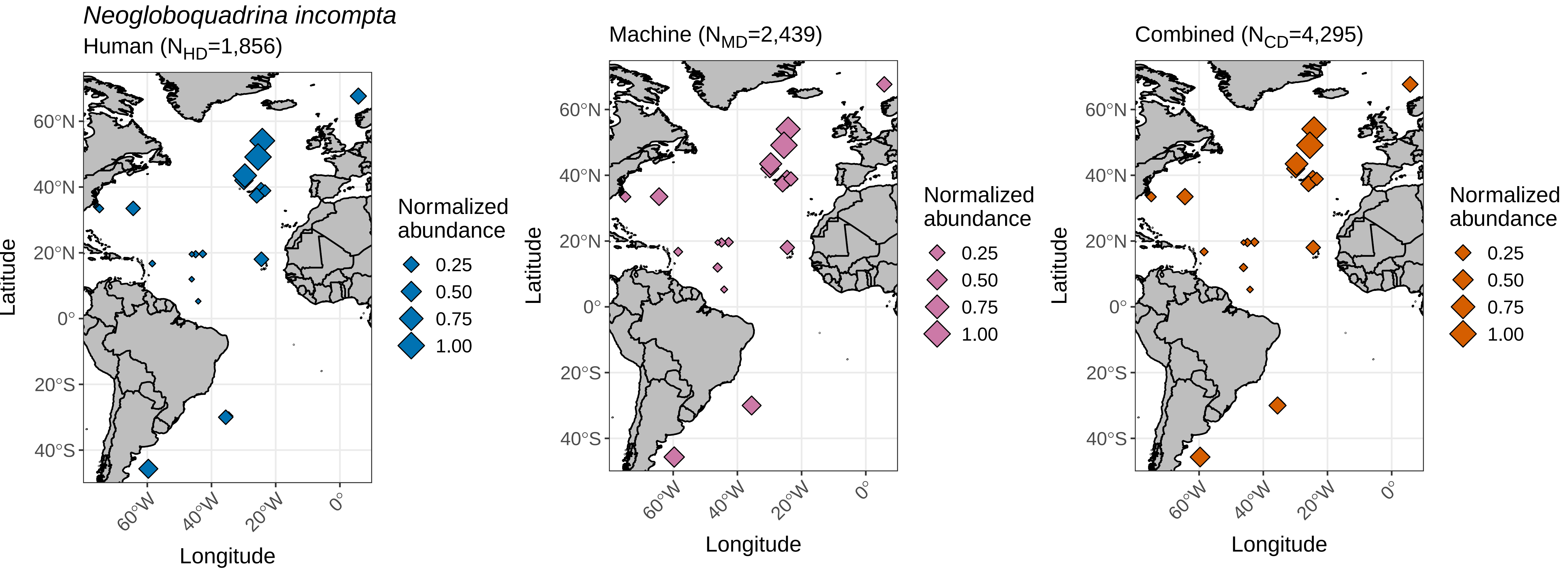

### neogloboquadrina_pachyderma_global_range_plot.pdf

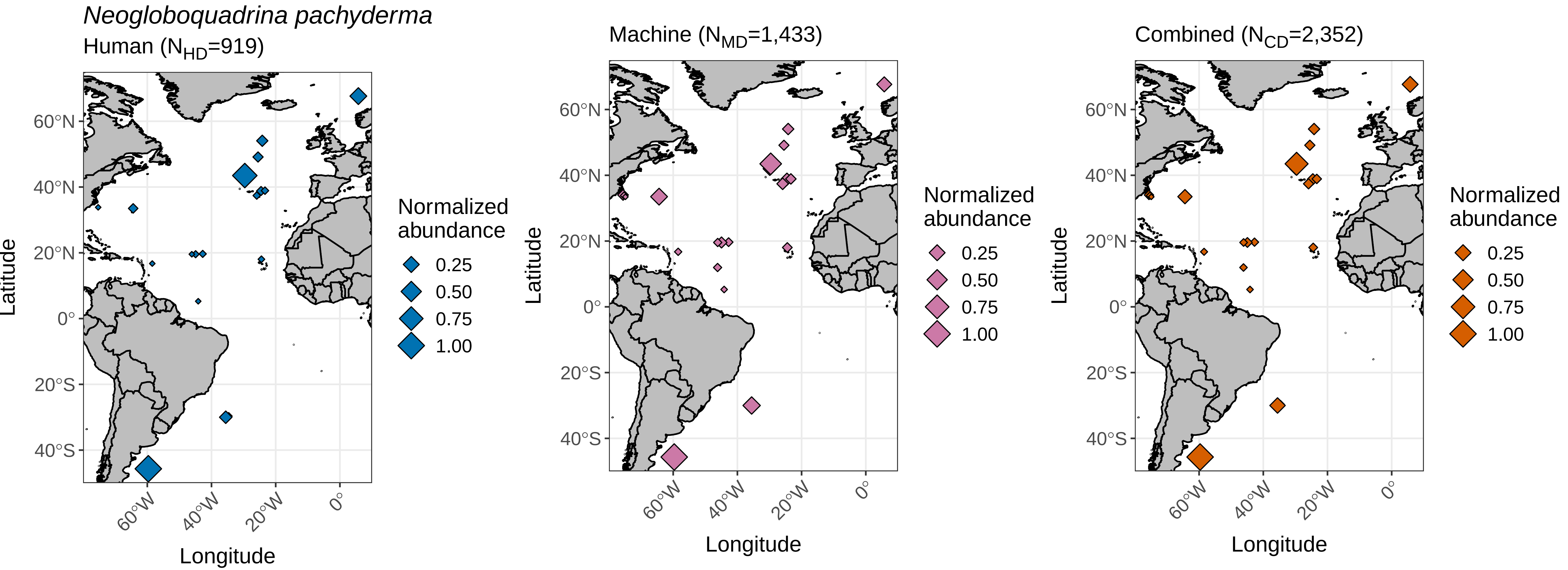

### orbulina_universa_global_range_plot.pdf

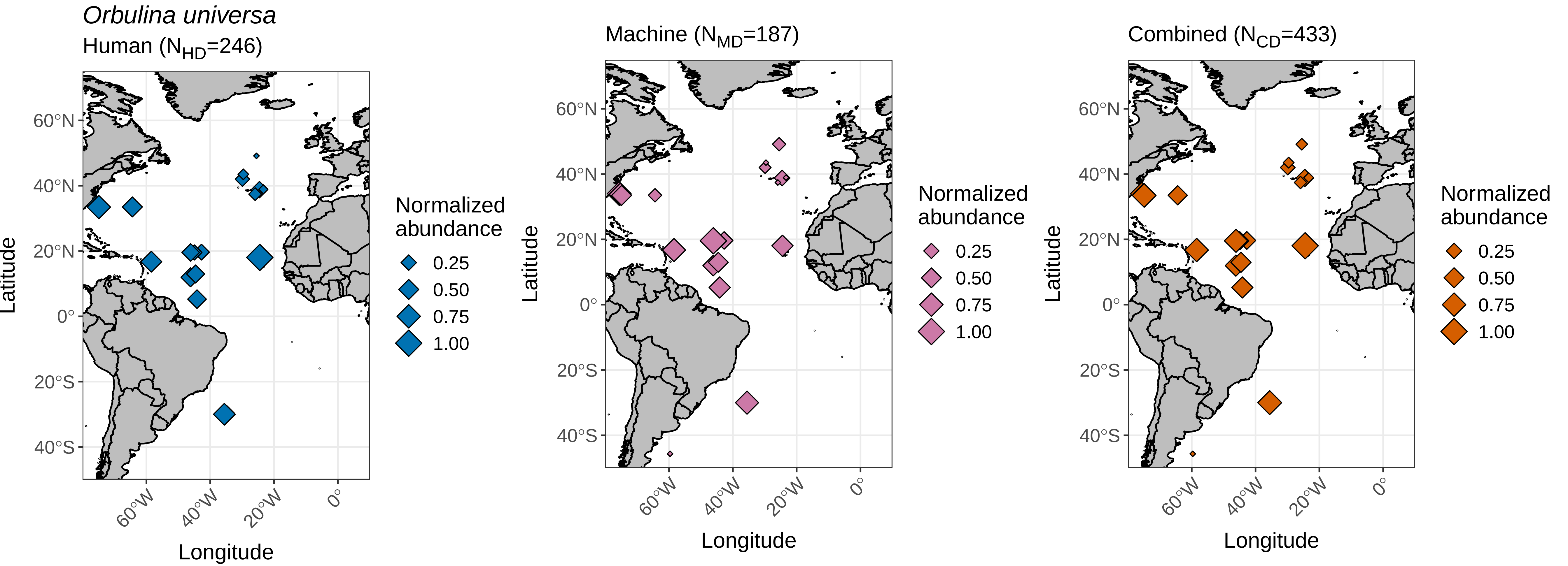

### pulleniatina_obliquiloculata_global_range_plot.pdf

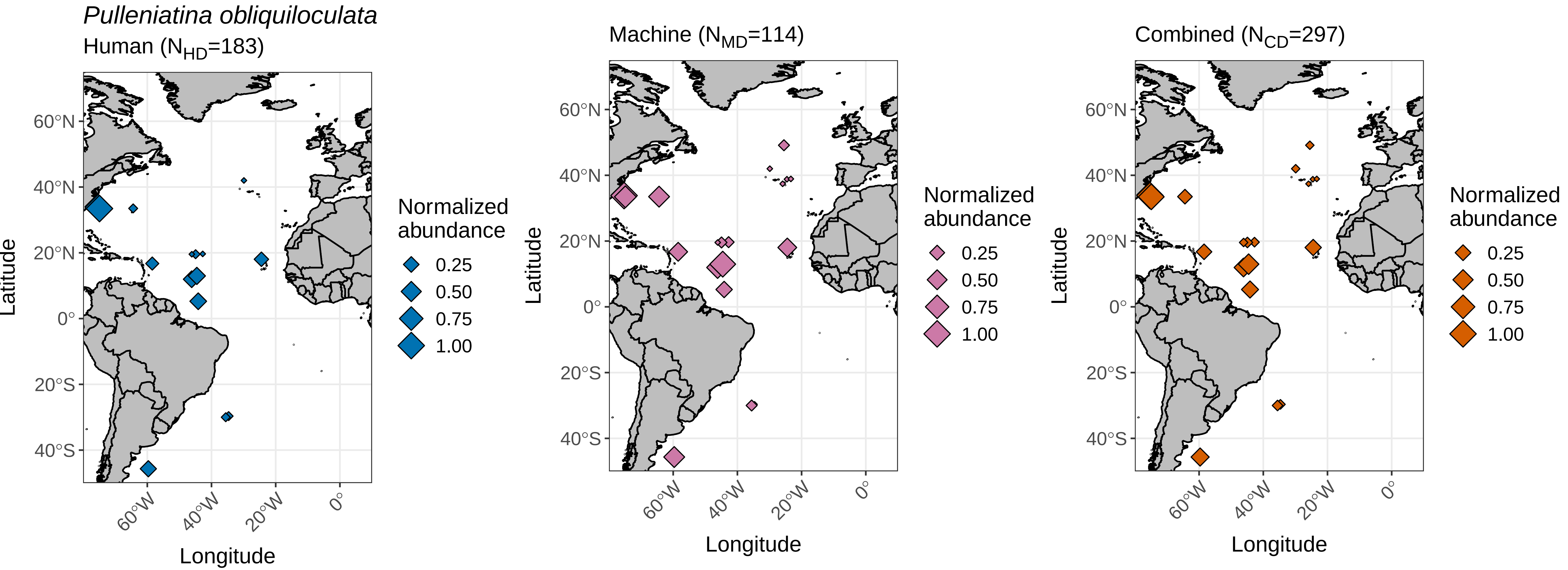

### sphaeroidinella_dehiscens_global_range_plot.pdf

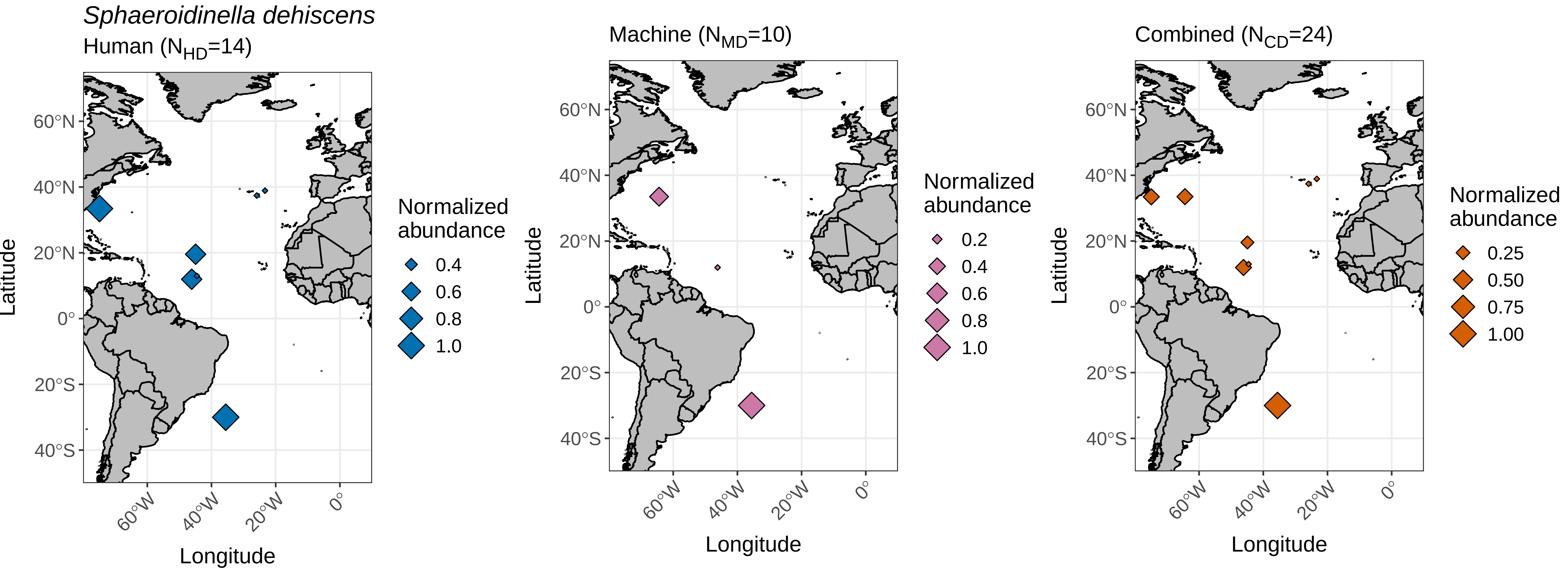

### turborotalita_humilis_global_range_plot.pdf

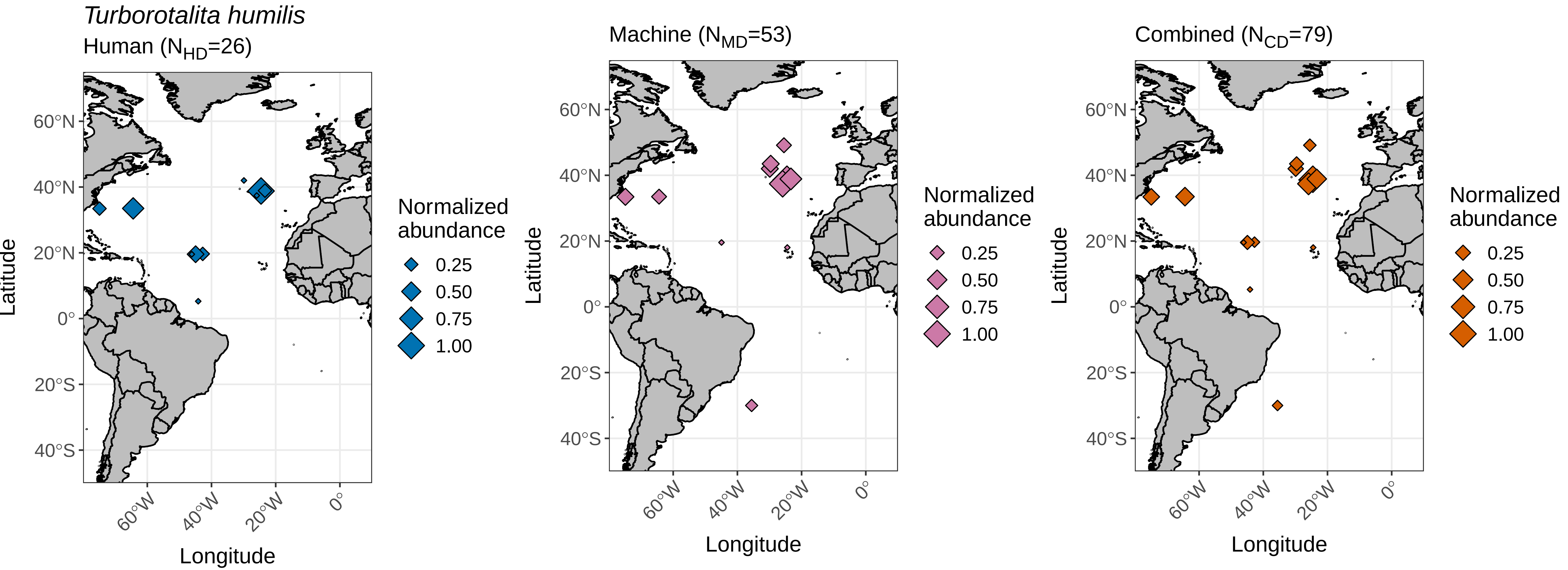

### turborotalita_quinqueloba_global_range_plot.pdf

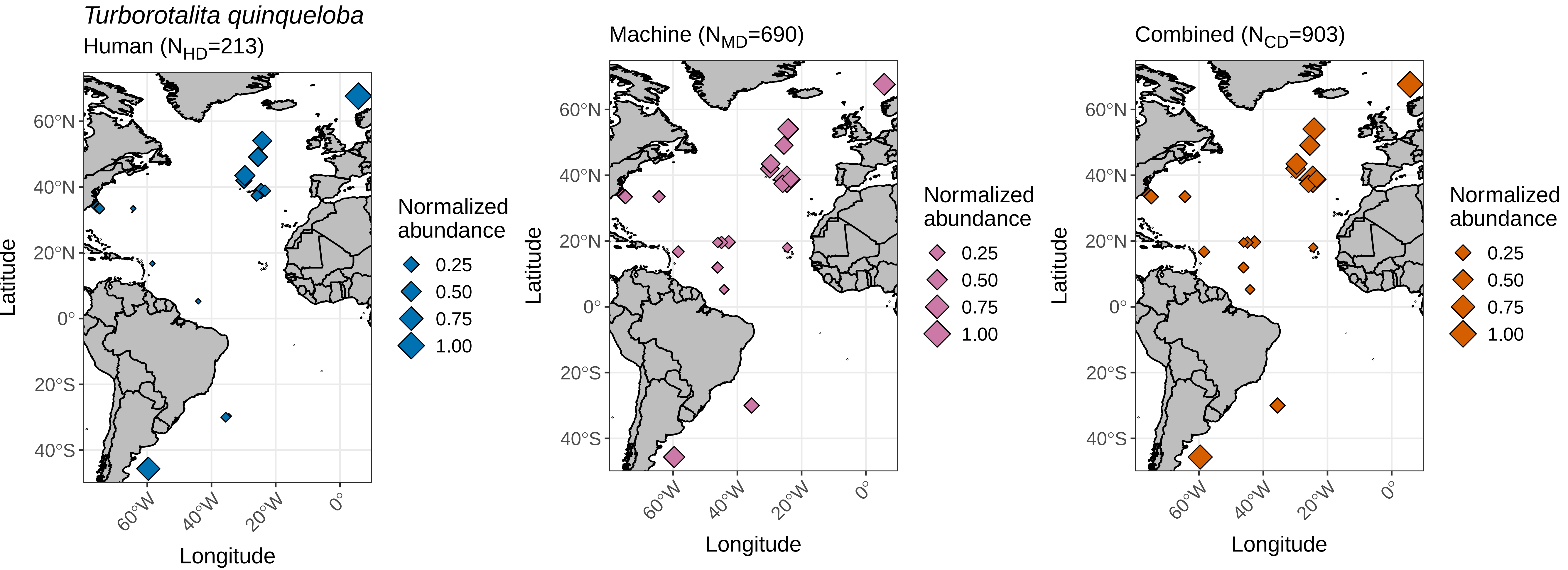
